# Time-dependent Mitochondrial Remodeling in Experimental Atrial Fibrillation and Potential Therapeutic Relevance

**DOI:** 10.1101/2025.01.29.635508

**Authors:** XiaoYan Qi, Feng Xiong, Jiening Xiao, Kalai Mangai Muthukumarasamy, Yasemin Altuntas, Yuan Zhong, Issam Abu-Taha, Florian Bruns, Marcel Tekook, Markus Kamler, Louis Villeneuve, Anna Nozza, Martin Sirois, Jason Karch, Philippe Pasdois, Donald M. Bers, Dobromir Dobrev, Stanley Nattel

## Abstract

**BACKGROUND:** Changes in mitochondria have been implicated in atrial fibrillation (AF), but their manifestations and significance are poorly understood. Here, we studied changes in mitochondrial morphology and function during AF and assessed the effect of a mitochondrial-targeted intervention in a large animal model.

**METHODS AND RESULTS:** Atrial cardiomyocytes (ACMs) were isolated from dogs in electrically-driven AF for periods of 24 hours to 3 weeks and from humans with/without longstanding persistent AF. Mitochondrial Ca^2+^-concentration ([Ca^2+^]_Mito_), reactive oxygen species (mtROS) production, membrane potential (ΔΨ_m_), permeability transition-pore (mPTP) opening and flavin adenine dinucleotide (FAD) were measured via confocal microscopy; nicotine adenine dinucleotide (NADH) under ultraviolet light. mtROS-production increased within 24 hours and superoxide-dismutase type-2 was significantly reduced from 3-day AF. [Ca^2+^]_Mito_ and mPTP-opening frequency/duration increased progressively during AF. Mitochondrial depolarization was detectable 24 hours after AF-onset. NADH increased by 15% at 24-hour AF, concomitant with increased pyruvate-dehydrogenase expression, then gradually decreased. Mitochondria enlarged and elongated at 24-hour and 3-day AF, followed by progressive fragmentation, rupture and shrinkage. Mitochondrial fusion protein-1 (MFN1) was reduced from 3-day to 3-week AF and phosphorylated dynamin-related protein-1 (p-DRP1ser-616) increased after 1 week of canine AF and in human AF. Addition of the mitochondrial antioxidant MitoTempo attenuated action-potential shortening and L-type Ca^2+^-current (I_CaL_)-downregulation in canine and human AF ACMs *in vitro*. Administration of the orally-active mitochondrial-targeted ubiquinone mitoquinone to dogs during 3-week AF prevented mitochondrial Ca^2+^-overload, mtROS-overproduction, structural damage and abnormalities in ΔΨ_m_ and respiration. Functionally, mitoquinone reduced AF-induced Ca^2+^-current downregulation, action-potential abbreviation, contractile dysfunction and fibrosis, preventing AF-substrate development and AF-sustainability.

**CONCLUSIONS:** Mitochondria show a series of changes during AF, with early hyperfunction and enhanced ROS-generation, followed by progressive damage and dysfunction. Mitochondrial-targeted therapy prevents mitochondrial dysfunction and attenuates adverse AF-related remodeling, positioning mitochondrial protection as a potential novel therapeutic target in AF.

## INTRODUCTION

Atrial fibrillation (AF) is the most common sustained cardiac arrhythmia and is a significant source of cardiovascular morbidity and mortality.^1,2^ The mechanisms that initiate and promote AF are unclear and treatment efficacy is suboptimal.^3^ Mitochondria occupy roughly 33% of cardiomyocyte volume and generate more than 95% of cardiac ATP, providing energy for a vast range of intracellular processes, including mechanical work and active ion transport.^4–6^ Mitochondria are the predominant source of intracellular reactive oxygen species (ROS), as well as the main target of ROS-induced signaling, and are a key regulator of local Ca^2+^-concentrations in cellular microdomains.^3,7^ The elevated atrial cardiomyocyte frequency in AF is expected to put added stress on mitochondrial function and modulate mitochondrial Ca^2+^-homeostasis. Furthermore, recent studies have linked mitochondrial dysfunction to AF-promotion,^5,6,8^ raising the possibility of maladaptive positive feedback on mitochondrial function in AF. Excess ROS have been implicated in pathogenesis of AF by affecting ion channels and propagation of the action potential.^9^ However, the time-course of mitochondrial changes and their pathophysiological role during the progression of AF are poorly understood. Here, we aimed to characterize in detail the changes in mitochondrial morphology, Ca^2+^-handling, protein expression and ROS generation over time after the induction of electrically-maintained persistent AF in a canine model. Our results suggested a potentially primary role of mitochondrial dysfunction in AF-progression, so we performed a prospective blinded experimental study with the orally-active mitochondrial-targeted agent mitoquinone (MitoQ) and found that it attenuated AF-progression. Finally, we repeated selected studies on human atrial tissue and cardiomyocytes (ACMs) from sinus-rhythm (SR) and AF patients, establishing evidence for the potential clinical relevance of our findings.

## METHODS

### For detailed methods, see Online Supplement

#### Animal Model and *In Vivo* Study

Animal-care procedures were approved by the Animal Research Ethics Committee of the Montreal Heart Institute (protocol: 2020-47-07) and followed Canadian Council on Animal Care Guidelines. For the initial characterization study, a total of 58 adult Foxhound dogs were studied, divided into control (n=15, 23.4±3.4 kg, 2.0±0.3 years-old, Female/Male ,F/M:7/8); 24-hour (n=11, 21.8±4.1 kg, 2.5±0.5 years, F/M:5/6 ), 3-day (n=10, 22.1±3.3 kg, 2.4±0.5 years, F/M:6/4), 1-week (n=11, 22.4±2.3 kg, 2.1±0.3 years, F/M:5/6) and 3-week AF (n=11, 21.8±2.7 kg, 2.4±0.5 years, F/M:5/6). A unipolar pacing lead was inserted into the right-atrial (RA) appendage under fluoroscopic guidance and connected to a pacemaker in the neck. The pacemaker was programmed to maintain AF by pacing the RA at 600 bpm to maintain AF electrically and mimic the atrial remodeling associated with spontaneous AF.^10^ Hemodynamic data were obtained with fluid-filled catheters and transducers.^11,12^

For the *in vivo* mitochondrial-targeted treatment study, 20 additional Foxhound dogs (21.4±1.7 kg, 2.1±0.3 years, F/M:12/8) were studied in the following 3 groups: 1) Sham group, instrumented but without atrial tachypacing (N=5); 2) AF+Placebo group, maintained in AF by atrial tachypacing and receiving placebo (empty capsule) once a day for 3 weeks (N=8); 3) AF+MitoQ group, receiving MitoQ (5 mg.kg^−1^ orally, once a day beginning 3 days before tachypacing-onset, N=7) during 3-week AF. The experimenter was blinded to dog therapy-assignment until the experiments were completed and data analyzed.

#### Canine ACM Isolation and Culture

ACMs were isolated and maintained with previously described methods.^13,14^

#### Human ACMs

Tissue-specimens from right-atrial appendages were obtained from patients (>18 years) undergoing elective open-heart surgery, with a pre-existing diagnosis of longstanding persistent (chronic) AF (cAF) or SR. Written informed consent was obtained from every patient, with protocols approved by the ethical review board of the University Duisburg-Essen, Germany (#12–5268-BO). Human ACMs were isolated from right-atrial appendage tissue using enzymatic digestion as previously described.^15,16^

### Confocal Imaging

#### Mitochondrial ROS (mtROS)

##### 1) Canine ACMs

For mitochondrial ROS detection, laminin-coated coverslips with adherent myocytes were incubated with MitoSOX Red (2.5 μM, Thermofisher Scientific, M36008) for 70 min at 37° C in the dark^17^ and then washed free of extraneous MitoSOX. Figure S1A shows autofluorescence recordings of unlabeled ACMs. Fluorescence intensity and images were acquired using a confocal microscope, with images analyzed offline using ImageJ software.^18^

##### 2) Human ACMs

After incubation, cells were transferred to a chambered slide (Ibidi) for subsequent confocal imaging. Some cells contained large quantities of lipofuscin granules and high autofluorescence levels (Figure S1B). These regions and the nucleus were excluded from fluorescence analysis.

### Mitochondrial Membrane Potential (ΔΨ_m_) and Permeability Transition Pore (mPTP) Opening

Mitochondrial membrane potential was visualized with tetraethyl benzimidazolyl carbocyanine iodide (JC-1, 5 μM, Thermofisher Scientific, T3168), which is a sensitive marker for mitochondrial membrane potential.^19,20,21^ Images were analyzed offline using ImageJ software. Tetramethylrhodamine methyl ester (TMRM) was used to record rapid changes in Ψ_m_ ^22^ as a decrease in mitochondrial fluorescence.^23^

### Mitochondrial and Cytosolic Ca^2+^ -Transient Measurement

A cold-warm incubation protocol was used to load the mitochondria with Rhod 2-AM.^24–27^ Cytosolic Ca^2+^-transients were measured with the Ca^2+^-indicator Fluo-4 AM.^13^ For details of measurements, see Online supplement.

### Nicotinamide Adenine Dinucleotide (NADH) Autofluorescence

NADH autofluorescence reflects the activity of the mitochondrial electron transport chain (ETC) as well as substrate supply.^27^ The uncoupling agent 1 μM carbonyl cyanide 4-(Trifluoromethoxy phenylhydrazone) (FCCP) was used to stimulate maximal respiration and induce minimum NADH autofluorescence. The complex IV inhibitor 5 mM sodium cyanide (NaCN, Millpore Sigma, # 380970-5), which fully inhibits respiration, was added to record the maximum autofluorescence signal (Figure S2A).^27–29^

### Flavin Adenine Dinucleotide (FAD) Autofluorescence

The mitochondrial uncoupler FCCP (1 μM) was applied to stimulate maximal respiration and record the maximal FAD signal. The subsequent application of NaCN was used to suppress respiration, preventing FADH_2_ oxidation and producing the minimum FAD signal.^27^

### Transmission Electron Microscopy (TEM)

TEM was used to evaluate mitochondrial structure. For details, see Online Methods.

### Patch-clamp recording and cell contraction

Action-potential (AP) and L-type Ca^2+^-current recordings were obtained with patch-clamp.^14,30^ Raw data for sarcomere shortening and relaxation was collected and stored using IonWizard software.^31^ For details, see Online Supplement.

### Optical Mapping

Previously-described methods were used.^32^ Data were processed with custom software written in Matlab (The MathWorks).^33^

### Quantitative Real-Time PCR and Western Blot

See Online Supplement.

### Histology

LA fibrosis was quantified by an observer blinded to experimental group.^34^

### Data Analysis

For details of statistical analysis, see Supplement. Mixed-effects models were used for all analyses, with human data analyzed as previously described.^35^

## RESULTS

### *In Vivo* Experiments

Typical AF-induced remodeling changes were seen *in vivo*. LA effective refractory period (AERP) decreased and lost rate-dependence, with changes reaching steady-state within 24 hours (Figure S3A). Left-ventricular end-diastolic pressure (LVEDP) increased significantly from 3-day to 3-week AF (Figure S3B), whereas LA pressure (LAP) increased significantly only at 3-week AF (Figure S3C). The prevalence of sustained-AF induction increased significantly at 1 and 3 weeks (Figure S3D), as did mean AF-duration (Figure S3E).

### Time-Dependent Changes in Mitochondrial Function Mitochondrial ROS Generation

Figure 1A shows representative images stained with the mitochondrial-ROS (mtROS) indicator MitoSOX Red. Compared to control ACMs (CTL-ACMs), mtROS production was greatly increased in AF-ACMs at 24 hours and remained stable at longer AF durations (Figure 1B). To assess expression of enzymatic ROS scavengers and sources, we quantified the expression of superoxide dismutase-type 1 (SOD1, with predominantly cytosolic localization), SOD2 (a mitochondrial matrix-based antioxidant enzyme), NADPH oxidase 4 (NOX4, ROS-source localized primarily in mitochondria) and monoamine oxidase (MAO-A, predominantly mitochondrial).^36^ The Western-blots are shown, along with total protein controls, in Figure S4A-D. SOD1 (Figure S4A, E) and MAO-A (Figure S4D, F) were not different among groups. SOD2-expression decreased significantly from 3-day AF (Figure 1C, Figure S4B). NOX4 expression increased significantly only at 3-week AF (Figure 1D, Figure S4C). The mitochondrial respiratory chain is a major source of ROS under pathological conditions. Complex I, II and III proteins were significantly upregulated at 1- and 3-week AF (Figure S5). The enzyme protein-expression data point to increased sources and reduced scavengers of mtROS, with significant changes beginning after 3-day AF.

**Figure 1.**
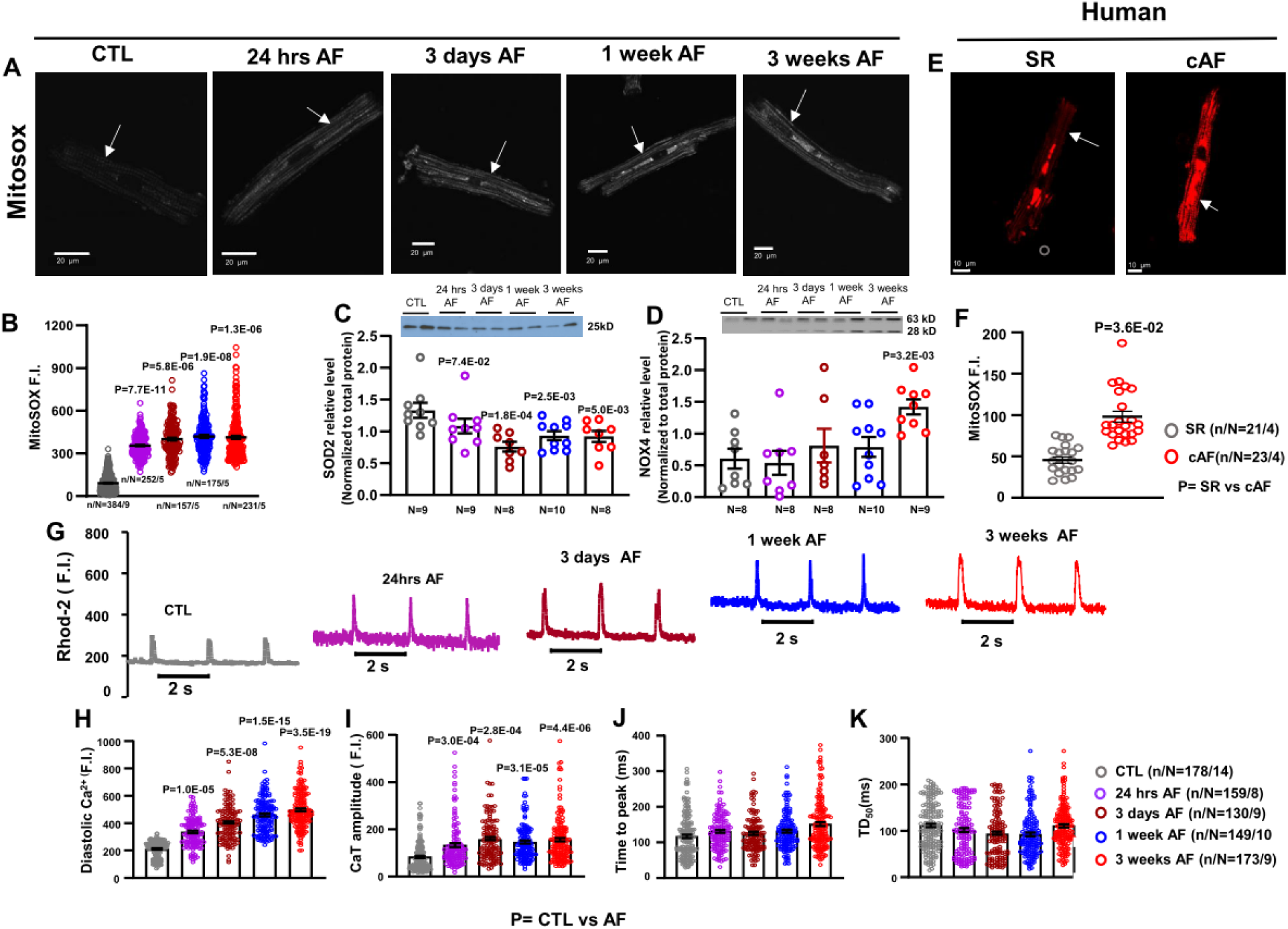
**A.** Representative Confocal image from canine CTL and AF cardiomyocytes labeled with MitoSox Red (arrows indicate individual mitochondria); **B**. Mean±SEM fluorescence intensity of MitoSox; **C.** Mean±SEM SOD2 protein expression level; **D.** Mean±SEM NOX4 protein expression level; **E.** Representative Confocal image of atrial myocytes labeled MitoSox Red from patients with SR and cAF; **F.** Mean±SEM fluorescence intensity of MitoSox ; G. Original recordings of [Ca^2+^] _mito_ transients in 0.5-Hz stimulated CMs; **H**. Mean±SEM diastolic [Ca^2+^] _mito_; **I.** CaT amplitude; **J.** Time to peak; **K.** Time from peak [Ca^2+^] to 50% decline [DT_50_] of mitochondrial CaTs from CTL and AF canine atrial cardiomyocytes.

The results of mtROS assay in human ACMs are shown in Figure 1E, F. mtROS was substantially greater in cAF-ACMs compared to SR-ACMs, in agreement with the canine model. Western-blot assays on human atrial tissue are shown in Figure S6. SOD2 expression was not altered in cAF atria (Figure S6A, C), while NOX4 was upregulated (Figure S6B, D).

### Mitochondrial Ca^2+^

[Ca^2+^]_mito_ plays an important role in controlling mitochondrial function. Figure 1G shows original [Ca^2+^]_mito_ recordings. Diastolic [Ca^2+^]_mito_ (end-cycle value at the onset of each activation at 0.5 Hz) and [Ca^2+^]_mito_ amplitude increased progressively during AF (Figure 1H, I). AF did not significantly affect [Ca^2+^]_mito_-transient kinetics (Figure 1J, K). [Ca^2+^]_mito_ recordings differed importantly from cytosolic [Ca^2+^] (Figure S7B), with much faster [Ca^2+^]_mito_ decay times (compare Figure 1G, with Figure S7B). Additional controls to confirm that the [Ca^2+^]_mito_ signals lack cytoplasmic [Ca^2+^] interference are that Rho-2 colocalizes tightly with mitochondrial indicator MitoTracker Green (Figure S8A-B) and that the [Ca^2+^]_mito_ transients are completely abolished by the mitochondrial Ca^2+^ uniport blocker Ru360 (200 nmol/L; Figure S8C). The protein expression of mitochondrial voltage-dependent anion channel (VDAC) and mitochondrial Ca^2+^ uniporter (MCU), which mediate Ca^2+^ influx into mitochondria, were not altered in both canine AF-ACMs and cAF-patient atrial tissue (Figure S9A-H).

### mPTP-Opening and Mitochondrial Membrane Potential (ΔΨ_m_)

We monitored mPTP-opening in intact ACMs using TMRM, a mitochondrial membrane-potential sensitive dye. Figure 2A shows recordings of mPTP-openings from individual mitochondria. CTL-ACMs did not exhibit spontaneous mPTP-opening. We observed transient mPTP-openings (examples indicated by red arrows)^37^ beginning after 24-hour AF. More prolonged mPTP-openings began to appear after 1-week AF. Both mean duration and frequency of mPTP-opening increased progressively over AF-time (Figure 2B, C).

**Figure 2.**
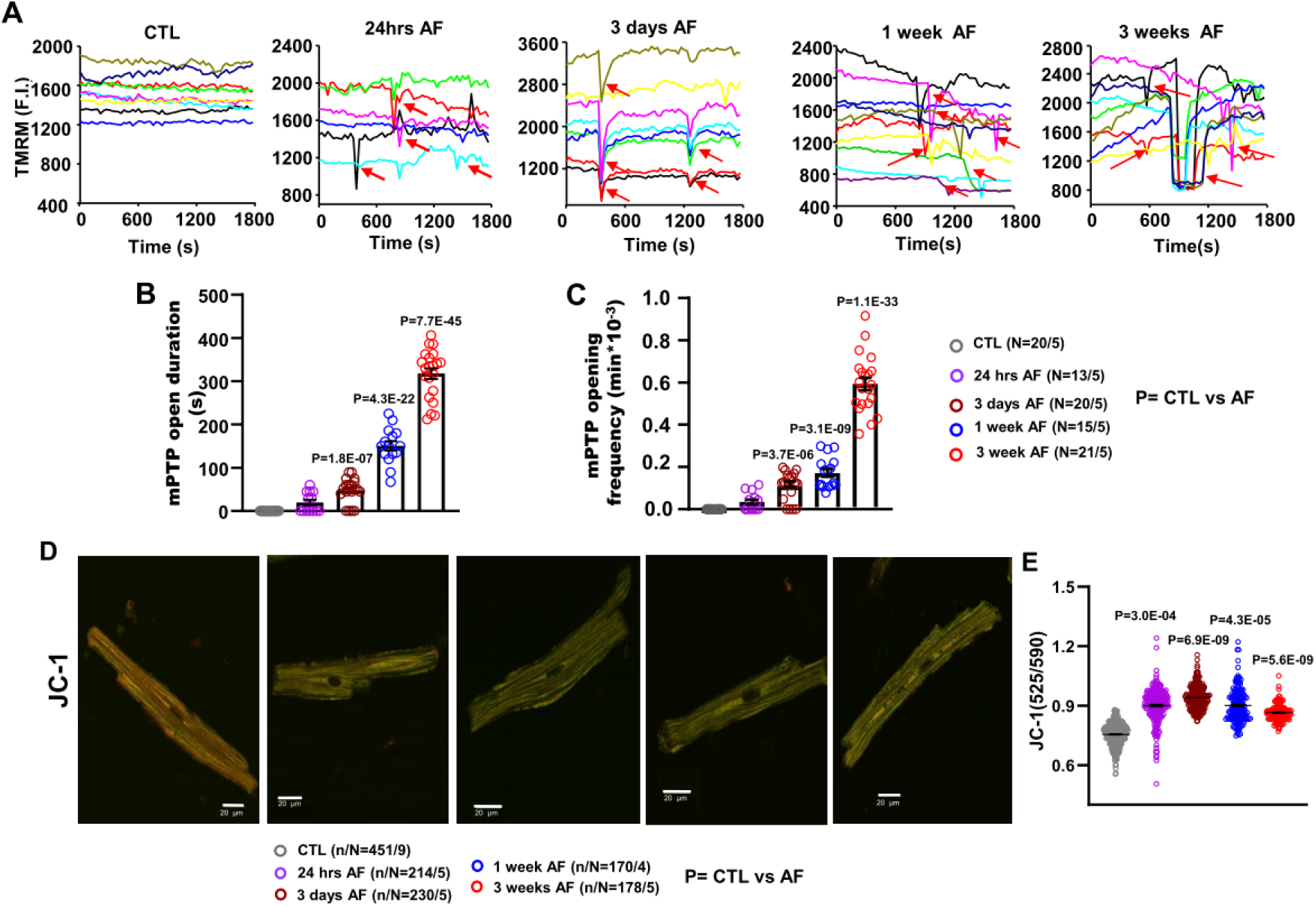
**A.** Traces were obtained from an individual mitochondrion, mPTP indicated by red arrows; **B.** Duration of mPTP-openings (TMRM to assess ΔΨm); **C.** Frequency of mPTP-openings; **D.** Confocal images of JC-1 fluorescence. ΔΨ_m_ of the CMs was measured by JC-1 from CTL and AF canine atrial cardiomyocytes; **E.** Mean data the ratio of aggregated and monomeric JC-1, indicating changes in mitochondrial membrane potential.

ΔΨm generated by the electron transport chain (ETC) is a key indicator of mitochondrial function and provides the driving force for ATP-synthesis.^37^ We used confocal microscopy to quantify JC-1 fluorescence in mitochondria as an indicator of ΔΨm. Figure 2D shows representative JC-1 images from CTL- and AF-ACMs. A shift from red (∼590 nm) to green (∼525 nm) staining reflects mitochondrial depolarization. Within 24 hours of AF onset, significant increases were noted in the average 525/590-nm JC-1 fluorescence ratio, indicating mitochondrial depolarization. These persisted through the 3 weeks of sustained AF, indicating rapid and persistent mitochondrial membrane depolarization during AF (Figure 2E).

### Changes in Mitochondrial Structure

Electron microscopy revealed marked mitochondrial ultrastructural remodeling in AF-atria. CTL-ACMs have regularly distributed sarcomeres and normal mitochondrial morphology (Figure 3A). Mitochondrial elongation and enlargement were observed following 24-hour and 3-day AF (Figure 3A-E), whereas thereafter the mitochondria became rounder and smaller. Individual mitochondria could vary considerably within a cell; Figure S10 shows frequency distribution histograms of mitochondrial morphological properties under control conditions versus various AF time-points. These histograms reflect the early shift towards larger surface area and perimeter, followed by a reversal of this trend, marked increases in roundness and reductions in aspect ratio.

**Figure 3.**
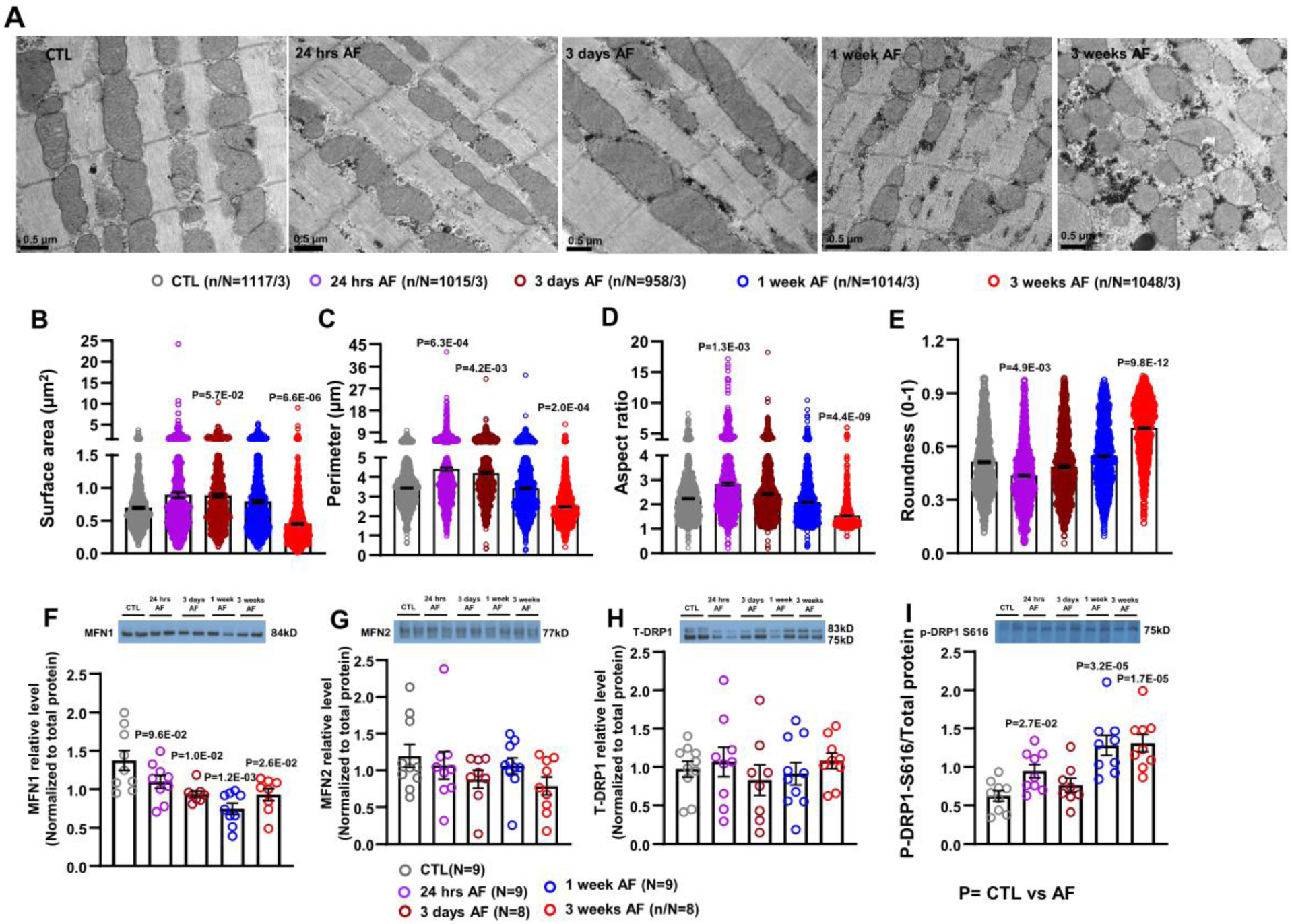
**A.** Representative transmission electron microscopy images of mitochondrial morphology from CTL and AF left atria; **B, C.** Quantitative analysis of mitochondrial morphology, mitochondrial surface area and perimeter; **D.** Aspect ratio (AR) is computed as [(major axis)/ (minor axis)] and reflects the “length to width ratio”; **E.** Roundness: [4 (surface area)/ (π·major axis^2^)] are two-dimensional indexes of sphericity with values of 1 indicating perfect spheroids; **F-I.** Mean±SEM MFN1, MFN2, T-DRP1 and pSRP1S616 protein expression level.

Changes in the ACM expression of proteins controlling mitochondrial dynamics are shown in Figures 3F-I. Expression of the mitochondrial fusion protein mitofusin-1 (MFN1) decreased progressively during AF (Figure 3F, Figure S11A), whereas MFN2 (Figure 3G, Figure S11A) and OPA1 (Figure S11C, E) protein expression did not change during AF. The serine616-phosphorylated form of the fission protein DRP1 (p-DRP1 S616) showed progressively increased expression in AF-ACMs (Figure 3I, Figure S11B). FIS1 (Figure S11C, F), total DRP1 (T-DRP1; Figure 3H, Figure S11B), and p-DRP1 S637 (Figure S11D, G) were not altered during AF. The changes in MFN1 and p-DRP1 S616 likely contribute to reduced mitochondrial size and increased roundness at 1- and 3-week AF, but do not account for the increased size at 24-hour AF.

MFN2 (Figure S12), T-DRP1 (Figure S13A, B) and p-DRP1 S637 (Figure S13A, E, F) were not changed in cAF patients, whereas p-DRP1 S616 (Figure S13A, C, D) was significantly increased in cAF patients, consistent with the results from canine AF-ACMs (Figure 3I). Glycogen-synthase expression increased significantly in canine ACMs at 3-week AF (Figure S14A, B). Increased glycogen synthase expression was observed in cAF patients (Figure S14C, D).

### Mitochondrial Respiration and PDH Complex

We measured the levels of NADH and FAD by their autofluorescence in ACMs.^28^ Mitochondrial NADH and FAD pools and redox index were estimated as illustrated in Figures S2A and S2B. Figures S15A and S15B show average results for NADH and FAD autofluorescence in CTL and at different AF durations. NADH pool increased significantly by 15% at 24-hour AF and then gradually decreased with AF-time to become significantly less than control at 1-week AF (Figure S15C). The NADH redox index (calculated from the maximally oxidized values in the presence of FCCP versus the maximally reduced values in the presence of NaCN) was significantly increased after 1- and 3-week AF (Figure S15D). The mitochondrial FAD pool decreased significantly at 1- and 3-week AF (Figure S15E). FAD redox index increased at 1- and 3-week AF (Figure S15F). This NADH and FADH redox shifts agree with higher ROS at longer AF times.

Pyruvate-dehydrogenase (PDH) protein-expression significantly increased only at 24-hour AF (Figure S16A, C). PDHA1 protein expression was significantly reduced from 3-day to 3-week AF (Figure S16B, D). PDK (1-4) catalyzes phosphorylation and inactivation of the PDH. PDK1 is present in heart.^38^ PDK1 increased significantly during AF-time (Figure S16E, G). The Ca^2+^-sensitive PDP1 is abundant in the heart^39^ and activates PDH.^40–42^ We observed a significant increase in the expression of PDP1 protein during AF-time (Figure S16F, H). Upregulation of total PDH and PDP1 may overcome the increase in PDK1 and be associated with increased NADH production in 24-hour AF-ACMs.

The results of Western-blot on human atrial tissue are shown in Figure S17. PDHA1 expression was significantly downregulated in cAF patients (Figure S17A, C). PDHA2 was not altered in cAF (Figure S17A, D), while PDK1 expression was upregulated in cAF (Figure S17B, E).

### Effects of Mitochondrial-ROS Targeted *In Vitro* Interventions

#### Effects of MitoTempo on AF-Related Changes in ACM Electrophysiology

Given the mtROS accumulation with AF in canine ACMs and the evidence for a role of ROS in AF,^43,44^ it is natural to consider that mitochondrial ROS-suppression might attenuate some of the electrical remodeling induced by AF. MitoTempo is a mitochondria-targeted superoxide dismutase mimetic that consists of piperidine nitroxide conjugated to the lipophilic triphenylphosphonium cation (TPP^+^) that facilitates the membrane potential-dependent accumulation of the compound into the matrix to suppress mitochondrial ROS.^45–47^ We incubated CTL- and AF-ACMs with MitoTempo (5 µmol/L) or vehicle for 7 hours, and then recorded APs and I_CaL_. Figures S18A and B show AP-recordings from canine CTL- and AF (1-week)-ACMs incubated with or without MitoTempo. AF significantly abbreviated APD_90_. MitoTempo application partially reversed AF-induced APD-shortening (Figure S18C). To assess cellular triggered activity, we monitored atrial APs from single ACMs after a 1-minute period of pacing at 2-Hz. Canine CTL-ACMs rarely exhibited spontaneous triggered activity post-pacing. MitoTempo application had no effect on CTL-ACMs (Figure S18E, F). However, 2-Hz-induced trigger activity greatly increased in canine (1-week) AF-ACMs. MitoTempo application reduced 2-Hz-induced trigger activity in AF-ACMs (Figure S18E, G). We then recorded I_CaL_ in ACMs from CTL and AF dogs with or without 5 µmol/L MitoTempo. Figures 4A and S19A show typical I_CaL_ recordings and Figure 4B shows mean current-voltage relations. Atrial tachypacing strongly reduced I_CaL_ in AF-ACMs compared to CTL-ACMs. MitoTempo application partially reversed AF-induced I_CaL_ reduction, but had no effect on I_CaL_ in CTL-ACMs (Figure 4A-B, S19).

**Figure 4.**
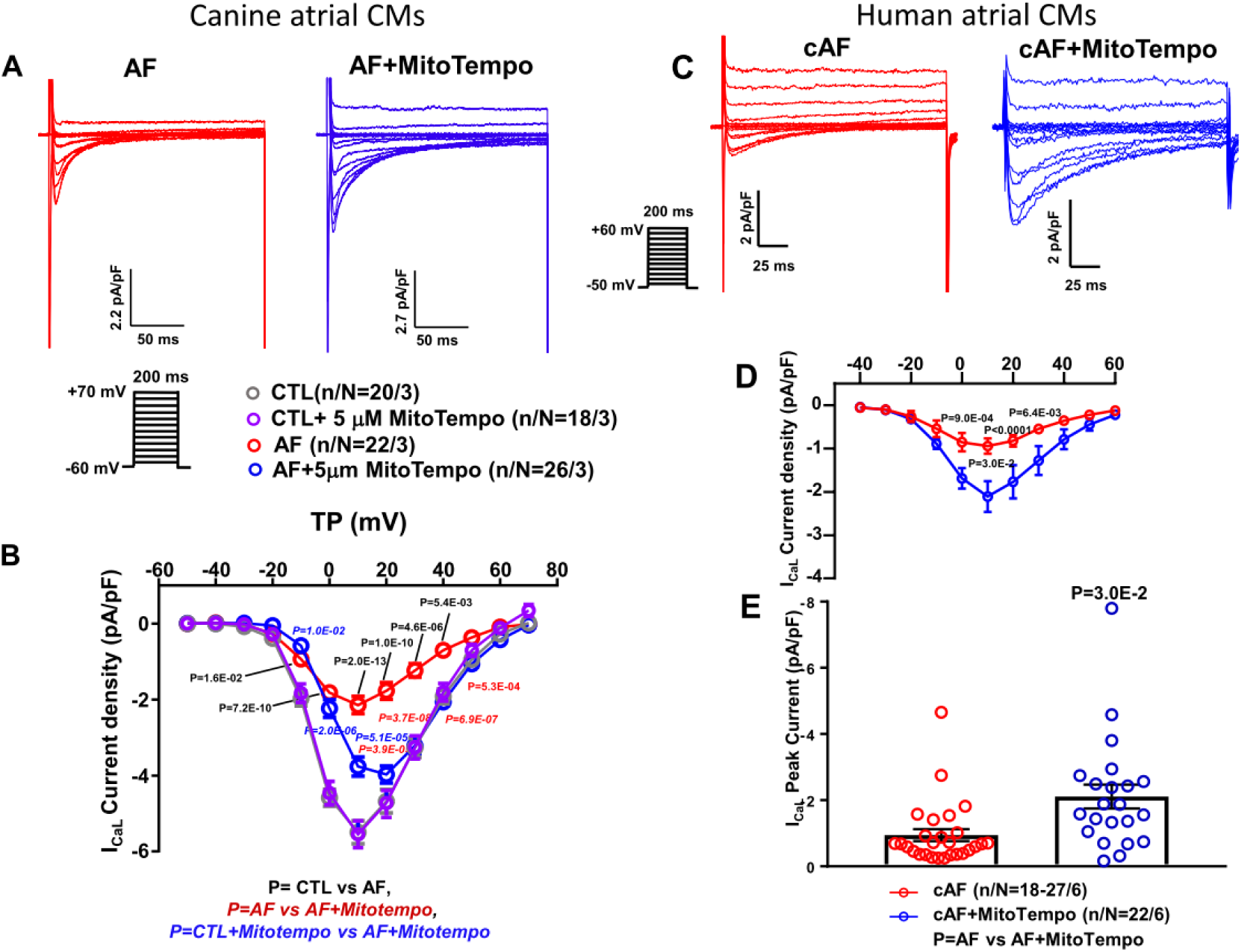
**A.** I_CaL_ recordings at 0.1 Hz from canine CTL and AF atrial CMs with/without MiotoTempo; **B.** Current-density voltage relation of I_CaL_ from canine CTL and AF atrial CMs with/without MitoTempo; **C.** I_CaL_ recordings at 0.1 Hz from atrial CMs of cAF patients with/without MitoTempo; **D.** Current-density voltage relation of I_CaL_ from atrial CMs of cAF patients with/without MitoTempo; **E.** I_CaL_ peak current densities at +10 mV from atrial CMs of cAF patients with/without MitoTempo.

Freshly-isolated human ACMs were divided into two fractions and incubated overnight (15-18 hours) with/without MitoTempo at ambient temperature. Figure S20A shows representative I_CaL_ recordings from human ACMs. I_CaL_ was significantly smaller in ACMs from cAF patients. The peak-I_CaL_ amplitude was 62% smaller in cAF (Figure S20A, C, E). MitoTempo administration significantly increased I_CaL_ in cAF-ACMs (Figures 4C-E). The peak-I_CaL_ amplitude was only 24 % lower in cAF+MitoTempo compared with SR-ACMs. Thus, the mitochondrial-targeted antioxidant MitoTempo reversed AF-induced electrical remodeling in both canine and human ACMs. We therefore decided to pursue studies of the *in vitro* effects of mitochondrial-targeted antioxidants, with the use of an agent that can also be administered orally.

#### Effects of Mitoquinone on Mitochondrial Ca^2+^

Mitoquinone (MitoQ) is a mitochondria-targeted antioxidant that has the ability to target mitochondrial dysfunction.^48^ Our results showed that AF significantly increased [Ca^2+^]_mito_ (Figure 1G,H). To evaluate the potential protective effect of MitoQ against AF-induced [Ca^2+^]_mito_ overload, ACMs from CTL and AF (multiple durations) canines were incubated with 200 nmol/L MitoQ or vehicle for 16 hours at 37°C and then [Ca^2+^]_mito_ transients were measured. *In vitro* MitoQ administration attenuated AF-induced [Ca^2+^]_mito_ overload in AF-ACMs at all AF-durations (Figure S21). MitoQ did not affect mitochondrial [Ca^2+^] in CTL-ACMs.

#### Mitochondrial Metabolism

We next used an *in vitro* tachypaced cultured canine ACM model, which we previously showed to mimic *in vivo* AF-induced cellular electrophysiological remodeling.^13^ We incubated 1-Hz (P1) and 3-Hz (P3) paced ACMs with 200 nmol/L MitoQ. Glucose-uptake was detected by 2NBDG. Glucose-uptake was greatly decreased in P3-ACMs (Figure S22A). MitoQ treatment partially prevented reduction of glucose-uptake induced by 24-hour 3-Hz pacing. We then measured NADH and FAD autofluorescence in 1- and 3-Hz paced ACMs. NADH and FAD pool were significantly decreased by 24-hour 3-Hz pacing, whereas MitoQ treatment prevented 3-Hz pacing induced NADH and FAD pool reduction (Figure S22B, C). Together, our *in vitro* experiments suggested that mitochondrial-targeted antioxidants can suppress AF-induced cardiac remodeling, so we proceeded to test the effects of an orally, clinically-available mitochondrial-targeted agent in AF-related remodeling.

#### Blinded Study of the Effects of Orally Administered Mitochondrial-Directed Therapy on AF-Associated Remodeling

To test the effects of orally-administered mitochondrial-targeted antioxidant therapy *in vivo*, dogs with 3-week AF were randomized to receive MitoQ (5 mg.kg^−1^, once a day)^49–51^ or placebo, with investigators blinded to treatment allocation until the completion of data analysis. Placebo-treated 3-week AF dogs had significantly shorter ERP compared with Sham animals. ERP abbreviation was attenuated by MitoQ, with ERP-values becoming not significantly different from Sham dogs (Figure 5A). Optically mapped RA-conduction velocities were decreased in the placebo-treated AF-group (by a mean of 10.8%). This decrease was attenuated by MitoQ and the mean conduction velocity in AF-mitoQ dogs was not significantly different from the Sham group (Figure 5B). APD_80_ was significantly abbreviated in placebo-treated animals, an effect that was significantly attenuated by MitoQ (Figure 5C). Placebo-treated dogs showed significantly greater AF duration (Figure 5E). Mean AF duration in MitoQ-treated AF-dogs averaged nearly 66% less than that in the placebo group, and not significantly different from Sham. AF was induced in 88% (7/8) animals of the placebo-treated group, versus 20% (1/5) of Sham and 43% (3/7) in MitoQ-treated AF-dogs (Figure 5F). MitoQ-treatment attenuated the cytosolic [Ca^2+^]-transient abnormalities induced by AF (Figure S23A-D), as well as the increase in spontaneous [Ca^2+^]-spark events resulting from AF (Figure S23E-F).

**Figure 5.**
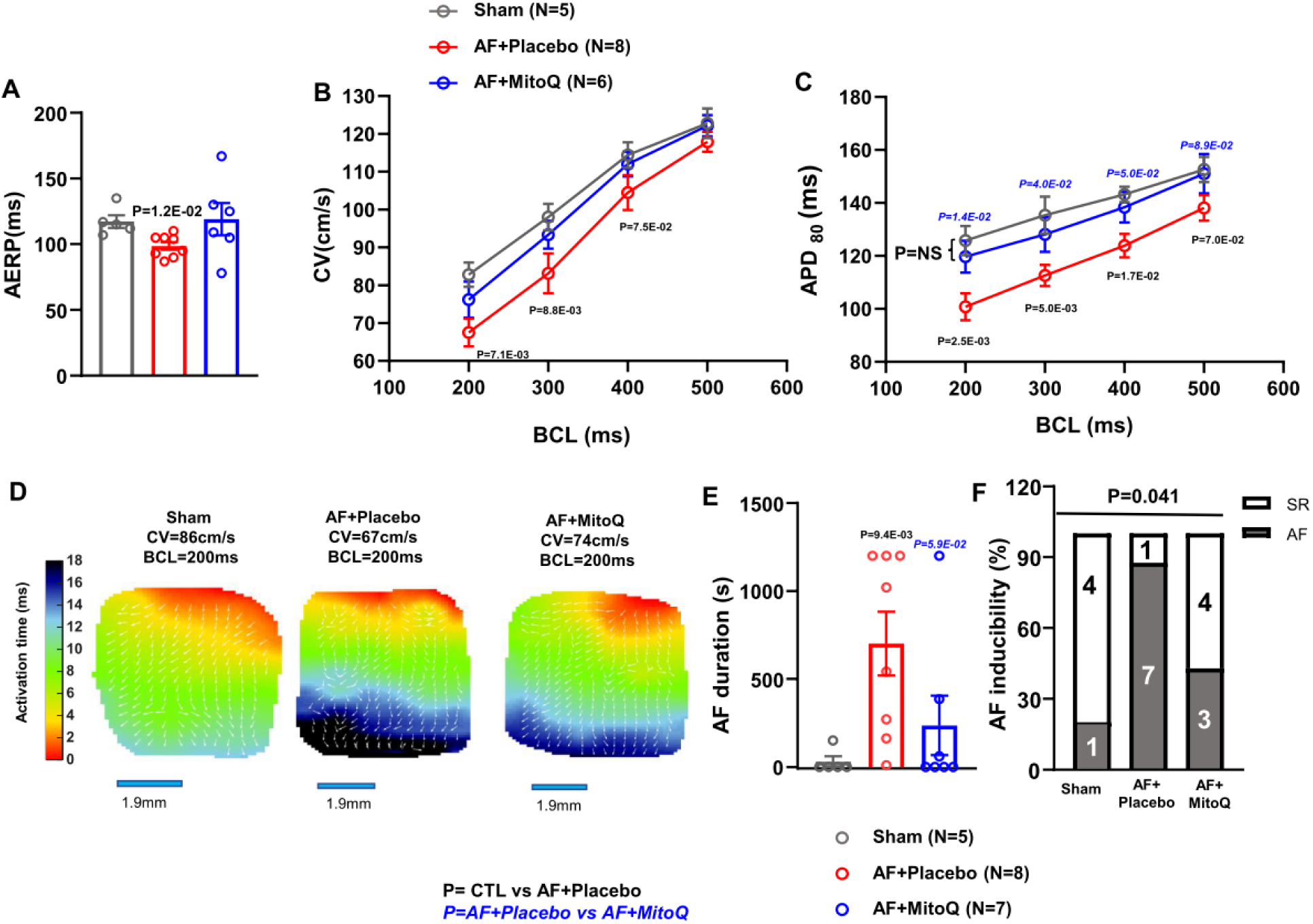
**A.** Mean±SEM effective refractory periods; **B.** Mean±SEM conduction velocities (CV); **C.** Mean±SEM APD_80_; **D.** Representative activation maps from Sham, AF+Placebo and AF+MitoQ canine right atria; **E.** Mean±SEM AF duration; **F.** Inducibility of AF from Sham, AF+Placebo and AF+MitoQ canine right atria.

#### *In Vivo* MitoQ Therapy Attenuates AF-Induced Oxidative Stress, Ca^2+^ Overload and ΔΨ_m_ Changes

To assess effects on AF-related oxidative stress and mitochondrial electrophysiological changes, we examined ACM mtROS in ACMs from AF-dogs treated with MitoQ or placebo. Figure 6A shows representative images of MitoSOX-Red staining in ACMs isolated from Sham, AF+placebo and AF+MitoQ dogs. Placebo-treated AF-ACMs showed a significantly increased mtROS production compared to Sham-ACMs, whereas MitoQ treated AF-ACMs showed significantly reduced mtROS production compared to placebo (Figure 6B). ETC Complex II contributes to ROS production in both physiological and pathophysiological condition.^52^ We observed a 116% increase in complex II protein expression in placebo-treated AF-ACMs compared to Sham. MitoQ treatment reversed AF-induced complex II overexpression (Figure S24), while not significantly affecting the expression of complexes I, III and IV.

**Figure 6.**
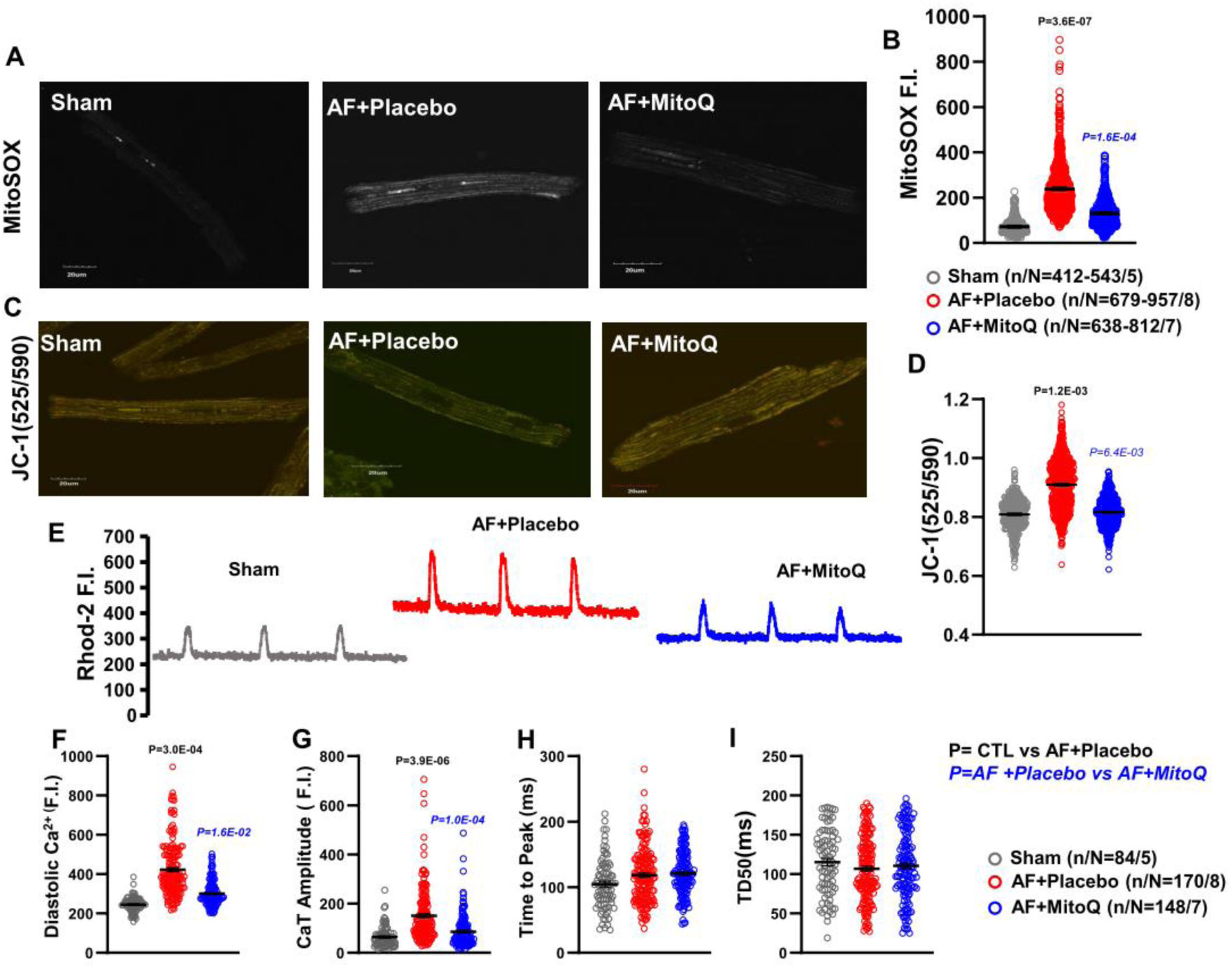
**A.** Representative image of MitoSOX red staining of superoxide radicals from Sham, AF+Placebo and AF+MitoQ canine atrial cardiomyocytes; **B.** Mean±SEM fluorescence intensity of MitoSox; **C.** Representative images of JC-1 fluorescence; **D** Mean±SEM ratio of monomeric and aggregated JC-1; **E.** Original recordings of [Ca^2+^] _mito_ transients in 0.5-Hz stimulated atrial cardiomyocytes from Sham, AF+Placebo and AF+MitoQ dogs; **F-I.** Mean±SEM diastolic [Ca^2+^] _mito_, CaT amplitude, time to peak and time from peak [Ca^2+^] to 50% decline [DT_50_].

Figure 6E shows mitochondrial Ca^2+^-transient recordings in 3 week AF dogs treated with MitoQ, versus placebo-treated and sham dogs. AF greatly increased mitochondrial diastolic [Ca^2+^] and Ca^2+^-transient amplitude but had no significant effect on kinetics. MitoQ-treatment suppressed AF-induced mitochondrial Ca^2+^-loading (Figure 6F-I). The mitochondrial transport proteins MCU and VDAC were not directly affected by AF (Figure S25).

The effect of MitoQ on ΔΨ_m_ of ACMs from CTL and AF dogs was assessed by live-cell imaging with JC-1. ACMs from placebo-treated AF dogs showed an increased JC-1 (green/red) fluorescence ratio compared to Sham-ACMs (Figure 6C, D), indicating ΔΨ_m_ depolarization, which was prevented by oral MitoQ treatment.

#### *In Vivo* MitoQ Therapy Ameliorates AF-Induced Mitochondrial Morphological and Biochemical Indices

Figure 7A shows representative transmission electron microscopy micrographs of Sham, AF+Placebo and AF+MitoQ canine LA tissue. MitoQ-treatment prevented mitochondrial morphological changes in AF-ACMs: the reductions in size and aspect ratio, the fragmentation, and accumulation of glycogen were greatly attenuated (Figure 7A-E). Analysis of the frequency distribution histograms also revealed that oral MitoQ resulted in preservation of mitochondrial surface area, perimeter, aspect ratio and roundness distributions (Figure S26). Western-blot analysis revealed that the expression levels of MFN1 (Figure 7F, Figure S27A) and MFN2 (Figure 7G, Figure S27B) protein declined significantly in AF-Placebo compared to Shams. MitoQ-administration significantly increased the level of MFN1 protein in AF-ACMs compared to Placebo-treated. MFN2 values were slightly but not significantly greater in AF+MitoQ vs AF+Placebo groups. Neither OPA1 nor DRP1 expression varied among groups (Figure S28).

**Figure 7.**
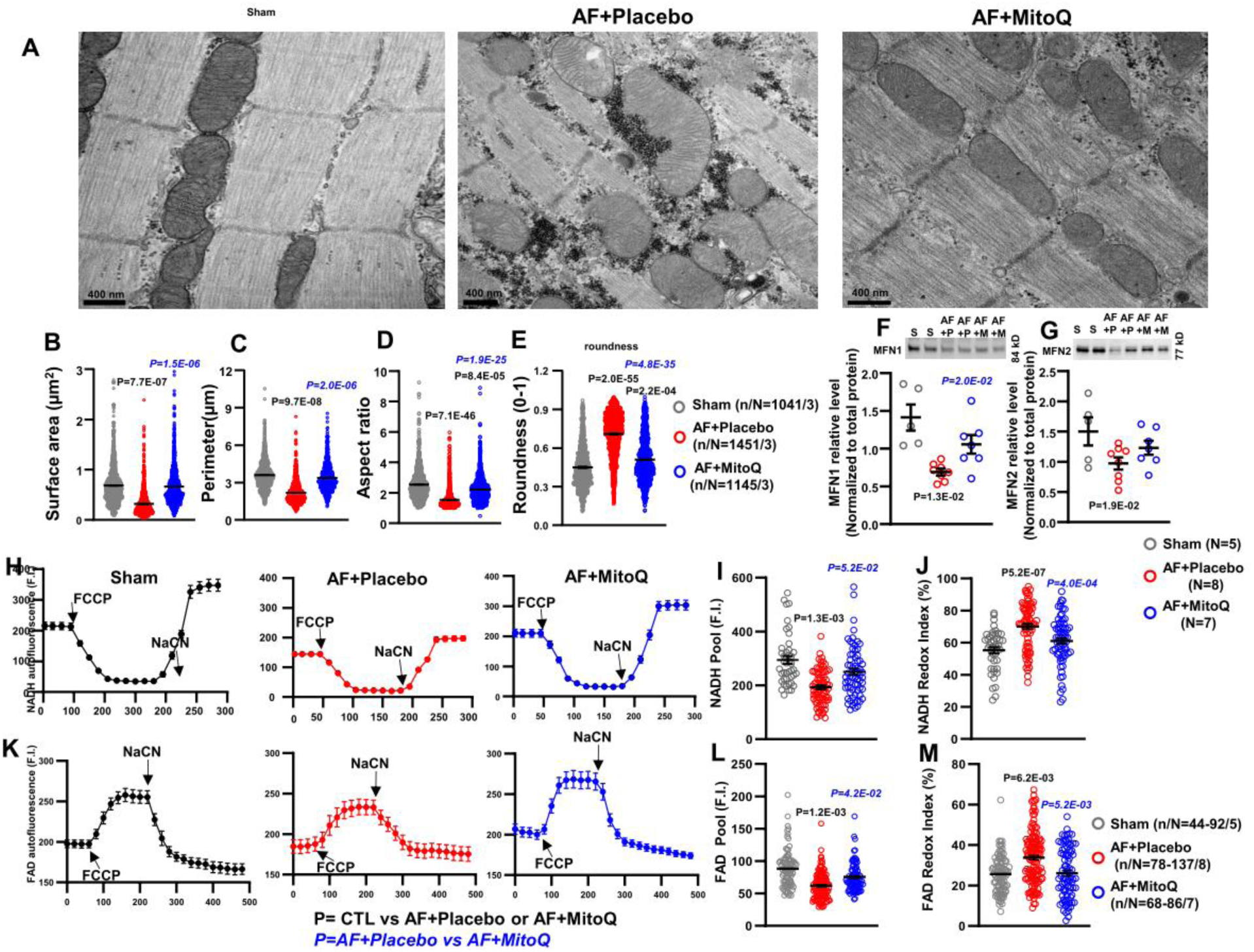
**A.** Representative transmission electron microscopy images from Sham, AF+Placebo and AF+MitoQ canine left atria; **B-E.** Mean±SEM mitochondrial surface area, perimeter, aspect ratio and roundness; **F, G.** MFN1 and MFN2 immunoblot band and Mean±SEM protein expression level from Sham, AF+Placebo and AF+MitoQ canine atrial CMs; **H.** Representative average traces for NADH autofluorescence; **I**, **J.** Mean±SEM mitochondrial NADH pool and NADH redox index; **K.** Representative average traces for FAD autofluorescence; **L, M.** Mean±SEM mitochondrial FAD pool and FAD redox index.

Figures 7H and 7K show average recordings of NADH and FAD autofluorescence in Sham, AF+Placebo and AF+MitoQ ACMs. Compared to Sham-ACMs, NADH and FAD pools were significantly reduced in AF+Placebo ACMs but partially restored in AF+MitoQ (Figure 7I and L). The redox indexes of NADH and FAD were significantly higher in AF+Placebo ACMs and again partially to fully restored in AF+MitoQ (Figure 7J and M). The PDHA1 unit of pyruvate dehydroxylase was significantly downregulated in AF (Figure S16B, D). While overall PDH expression was not affected by AF, with or without MitoQ (Figure S29A, B), the expression of the PDHA1 subunit, which was suppressed in AF+Placebo, was partially preserved in MitoQ-treated dogs (Figure S29C, D), which may have contributed to improved respiration as reflected by preserved NADH and FAD pools.

#### *In Vivo* MitoQ Therapy Prevents Atrial Hypocontractility, I_CaL_ Downregulation and Fibrosis in AF-dogs

AF-induced atrial contractile dysfunction has significant clinical implications by contributing to thromboembolic risk. Figure 8A shows representative recordings of sarcomere shortening in ACMs paced at 1-Hz. Overall sarcomere shortening, contraction and relaxation velocity were significantly reduced in AF-ACMs from placebo-treated canines compared to Sham-ACMs. MitoQ prevented AF-induced reductions in sarcomere shortening, while preserving the velocity of shortening and relaxation (Figure 8B-D).

**Figure 8.**
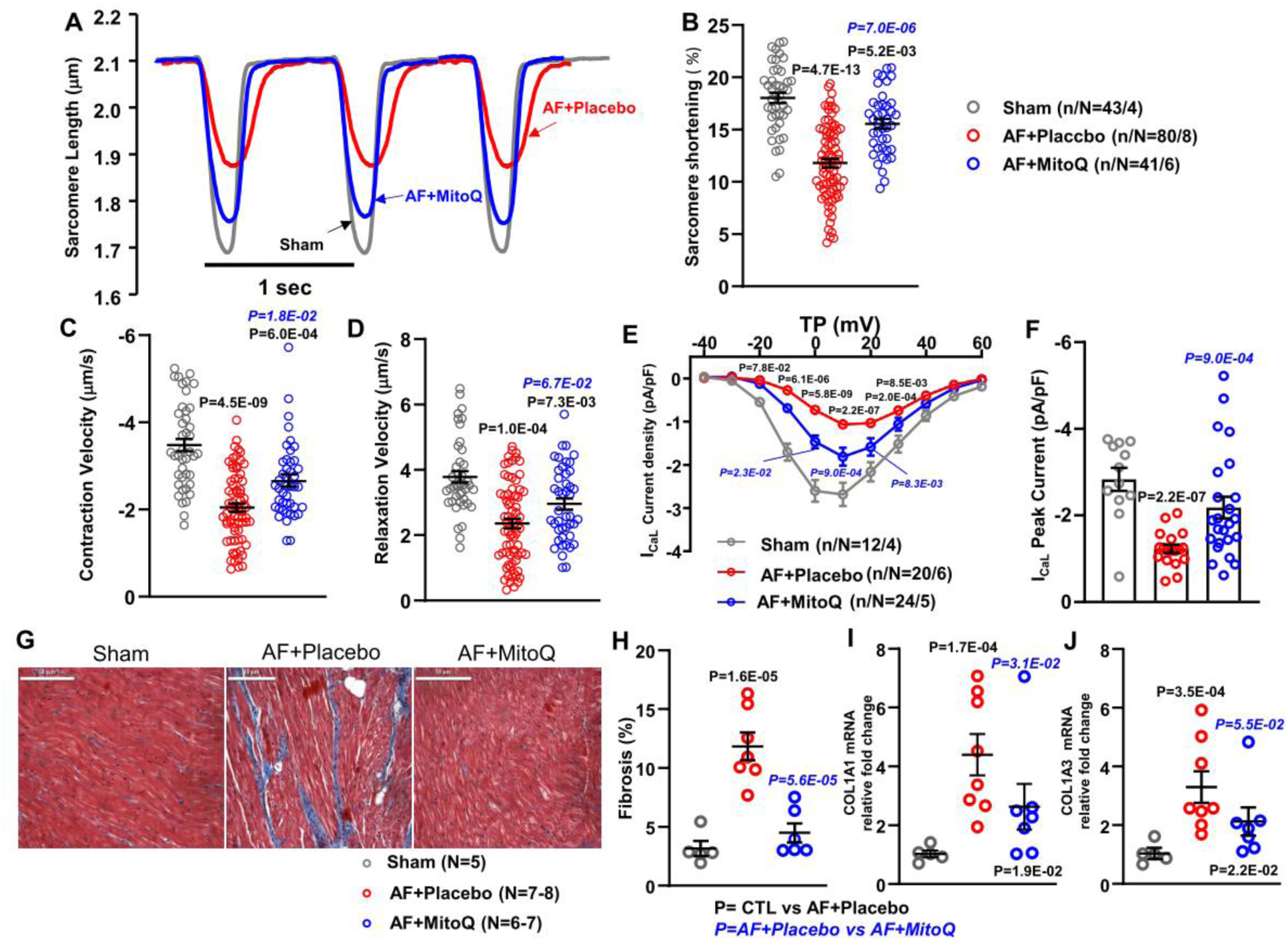
**A.** Typical example of recorded sarcomere shortening at 1-Hz; **B.** Mean±SEM percentage sarcomere shortening; **C.** Mean±SEM velocity of contraction; **D.** Mean±SEM relaxation velocity, **E.** Current-density voltage relation of I_CaL_ from Sham, AF+Placebo and AF+MitoQ canine atrial CMs; **F.** Peak *I*_CaL_ densities (at +10 mV); **G.** Representative images of Masson’s trichrome–stained left atria; **H.** Mean±SEM fibrous tissue content; **I.** Mean ± SEM gene expression of COL1A1 (fold changes); **J.** Mean±SEM gene expression of COL1A3 (fold changes).

I_CaL_ downregulation is an important contributor to the electrical remodeling caused by AF. Figure 8E shows data from Sham, AF+Placebo and AF+MitoQ ACMs. MitoQ administration attenuated I_CaL_ reduction in AF-ACMs (Figure 8E, F).

Fibrous-tissue content was significantly increased in AF+Placebo atria (Figure 8G, H) compared with Sham and this effect was significantly attenuated by MitoQ. Atrial *COL1A1* and *COL1A3* mRNA expression increased markedly in placebo-treated AF atria, and MitoQ treatment prevented AF-induced upregulation of collagen-gene expression (Figure 8I, J). Similar trends were observed for *COL1A2* and *FN*, but the changes were not statistically significant (Figure S30).

## DISCUSSION

In this study, we show progressive alterations in mitochondrial properties during the course of AF. We found that mitochondria show evidence of early activation (enlargement, increased NADH pool and mtROS production) followed by progressive damage (reduced size and increased roundness, glycogen accumulation, reduced NADH and FAD pools, increased MPTP-opening, and loss of mitochondrial membrane potential). Qualitatively similar changes are seen in atria of cAF-patients. Mitochondria-targeted antioxidant therapy *in vitro* alleviated AF-induced mitochondrial dysfunction and electrical remodeling in both canine and human ACMs, and oral treatment with a mitochondrial-targeted drug (MitoQ) attenuated adverse mitochondrial, cellular and electrical/structural remodeling, along with AF-promotion, in a chronic dog model. Our data establish mitochondrial ROS production and dysfunction as a very early and a potential causal contributor to AF-promoting atrial remodeling and position mitochondria-targeted interventions as a potential novel therapeutic option for AF treatment.

### Mitochondrial Dysfunction and AF

Oxidative stress plays a major role in the development and perpetuation of AF in patients and animal models.^53–55^ Mitochondria are the major source of cardiomyocyte ROS, and there is evidence that mitochondrial oxidative stress promotes AF.^8^ Mitochondrial dysfunction has been implicated in post-operative AF.^56,57^ Mitochondrial ROS production is a by-product of normal metabolism and electron leak from the ETC at complexes I, II, and III.^58,59^ Complex I catalyzes the transfer of two electrons from NADH to ubiquinone in a reaction that is coupled with the translocation of four protons across the membrane, contributing to the proton motive force.^60^ Mitochondrial complex III catalyzes the electron transfer from ubiquinol (QH2) to ferricytochrome c, which is coupled to proton translocation for ATP synthesis.^60^ Under physiological conditions, about 0.2–2% of the electrons that pass through the ETC leak from the system, generating superoxide (O_2_^•–^),^61^ the major source of mtROS. Mitochondria-localized proteins that contribute to the mtROS pool include p66^Shc^, monoamine oxidases (MAOs) and NOX4.^62^ Ausma et al. first noted reductions in phosphocreatinine and mitochondrial shrinkage in an AF goat model.^63^ Subsequently, Mihm et al. demonstrated reduced myofibrillar creatine kinase activity and increased protein oxidation in the atria of AF-patients, suggesting that myofibrillar energetic dysfunction and oxidative stress might be important.^43^ Mitochondrial dysfunction was more directly implicated subsequently by work from Wiersma et al.^6^ and Ozcan et al.^64^ Mouse atrial-tumor derived cardiomyocytes (HL-1 cells) paced at 6-Hz showed impaired mitochondrial Ca^2+^-handling and decreased mitochondrial membrane potential, respiratory-function and ATP-synthesis, while atrial tissue from cAF-patients showed abnormal ATP-content and fragmentation of mitochondria.^6^ Atrial samples from patients with terminal heart failure and AF showed further energy depletion and metabolic stress compared to those with SR, and an AF mouse model (LKB1-knockout) showed mitochondrial dysfunction and increased ROS production.^63^ Fossier et al. recently demonstrated mitochondrial Ca^2+^-handling abnormalities, enhanced expression and reduced single-channel activity of the MCU in mice with experimentally-induced metabolic syndromes as well as in patients with metabolic syndrome.^65^ The authors noted improvement of mitochondrial abnormalities and suppression of AF-inducibility in metabolic-syndrome mouse-preparations on exposure to the MCU-agonist kaempferol, leading to suggestions for the possible repurposing of ezetimibe to target mitochondrial MCU-mediated Ca^2+^-handling dysfunction in AF.^66^ We did not observe changes in MCU-expression in the present study; however, [Ca^2+^]_mito_ significantly increased in the AF model, with attenuation upon treatment with MitoQ.

AF increases atrial oxygen consumption about 4-fold and atrial blood-flow by up to 3-fold.^67^ At 24-hour AF mitochondrial hyperfunction, presumably compensatory, was reflected by mitochondrial enlargement, elongation and markedly increased ROS production. Thereafter, mitochondrial dysfunction developed progressively, manifested by structural changes (reduced size, rounding), changes in mitochondrial redox-related enzymes (downregulation of SOD2 from 3-day AF, upregulation of NOX4 at 3-week AF) and ETC proteins (upregulation of complexes I-III from 1-week AF), increases in mitochondrial Ca^2+^ (from 24-hour AF), enhanced mPTP-opening and loss of mitochondrial transmembrane potential. Production of mtROS remained elevated throughout the time course of AF, suggesting a potential role in cellular and more specifically mitochondrial damage associated with AF. Thus, our findings expand previous knowledge by determining the precise time course of mitochondrial damage during AF; and by showing that mitochondrial redox dysfunction is a causal contributor to AF-induced atrial remodeling and the evolution of the proarrhythmic atrial substrate, and as such a potential novel anti-AF drug target.

### Mechanistic Contribution of Mitochondrial Dysfunction and Potential Therapeutic Relevance

Pathophysiological contributions of mitochondrial dysfunction and ROS accumulation in AF have long been suggested.^43^ We obtained evidence of a potentially-significant pathophysiological role of mtROS accumulation in terms of the ability of the mitochondrial-targeting SOD-mimetic MitoTempo to attenuate abnormalities in I_CaL_ in canine and human AF-ACMs. I_CaL_-reduction is believed to be an important contributor to the atrial AP-abbreviation, refractoriness-shortening and arrhythmogenesis associated with atrial remodeling.^68^

Further evidence was provided by the ability of oral MitoQ-administration to prevent AF-induced mitochondrial dysfunction and adverse atrial remodeling. MitoQ is a ubiquinone that accumulates within mitochondria, containing a conjugated lipophilic triphenylphosphonium cation (TPP^+^)^69^ that does not require a specific uptake mechanism.^70^ The lipophilic TPP^+^ cation allows MitoQ to cross the phospholipid bilayer and to accumulate within the mitochondrial inner membrane, driven by the mitochondrial membrane potential.^48^ The positively charged residue (TPP^+^) of MitoQ is adsorbed to the matrix surface, while the hydrophobic end (ubiquinone) inserts into the hydrophobic core of the mitochondrial inner membrane.^71^ MitoQ is then reduced to its active ubiquinol form within mitochondria only. After detoxifying oxidants, the inactive ubiquinone form of MitoQ is recycled back to the active ubiquinol antioxidant form by complex II.^48^ Chronic oral MitoQ is under investigation in humans and a 6-week course has been shown to improve indices of endothelial function.^49^ Longer courses of therapy (≥6 months) have been administered to treat Parkinson’s disease, for which it proved to be ineffective.^72^ To the best of our knowledge, the effect of MitoQ on AF has not been previously studied, but our findings suggests that it may be a valuable lead compound for the development of mitochondria-targeted therapy to prevent development or progression of the AF-promoting substrate. MitoQ-treatment attenuated AF-induced ERP-abbreviation, impaired cell-contraction, cellular triggered activity and tissue-fibrosis, and is therefore effective against electrical, structural, Ca^2+^-handling and contractile remodeling, all of which are key elements of AF promotion.

### Potential Limitations

We carefully evaluated the changes in mitochondrial properties and the effects of mitochondria-targeted therapy in a specific large-animal model of AF, but no animal model perfectly mimics the clinical condition and further assessment in other large- and small-animal AF models would be of interest to validate our findings. Our studies in human atrial tissues and ACMs support the translational relevance of our findings, but further studies in both experimental and clinical models will be needed before the full applicability of our findings becomes clear. Abnormal ROS production and mitochondrial dysfunction might contribute to AF pathophysiology by mechanisms not addressed in present study. Further work is required to delineate the precise mechanisms through which oxidative stress and mitochondrial dysfunction induce atrial remodeling and related arrhythmogenesis.

### Conclusions

Mitochondria show time-dependent changes during AF, which contribute to the progression of the arrhythmia. Mitochondria-targeted therapy merits further consideration as a therapeutic avenue, with mitoquinone being a potentially-interesting lead compound.

## Nonstandard Abbreviations and Acronyms

AERPs: Atrial effective refractory periods
AF: Atrial fibrillation
APD: Action potential duration
cAF: Chronic atrial fibrillation
CaT: Ca^2+^ transient
CMs: Cardiomyocytes
CTL: Control
CV: Conduction velocity
DRP1: Dynamin-related protein 1
ETC: Mitochondrial electron transport chain
FAD: Flavin adenine dinucleotide
FIS1: Fission protein 1
I_CaL_: L-type Ca^2+^-current
LA: Left atrial
LAP: Left atrial filling pressure
LVEDP: Left ventricular end-diastolic pressure
MAO-A: Monoamine oxidases
MCU: Mitochondrial Ca^2+^ uniport
MFN1: Mitofusin-1
MFN2: Mitofusin-2
MitoQ: Mitoquinone mesylate
mPTP: Mitochondrial permeability transition pore
mtROS: Mitochondrial reactive oxygen species
NADH: Nicotinamide adenine dinucleotide
NADPH: Nicotinamide adenine dinucleotide phosphate
NOX4: NADPH oxidase 4
OPA1: Optic atrophy 1
OXPHOS: Oxidative phosphorylation
PDH: Pyruvate dehydrogenase
PDHA1: Pyruvate dehydrogenase E1 component subunit alpha
PDK1: Pyruvate dehydrogenase kinase 1
PDP1: Pyruvate dehydrogenase phosphatase 1
RA: Right atrial
RP: Resting membrane potential
ROS: Reactive oxygen species
SOD1: Superoxide dismutase 1
SOD2: Superoxide dismutase 2
SR: Sinus rhythm
TCA: Tricarboxylic acid
TEM: Transmission electron microscopy
TMRM: Tetramethylrhodamine methyl ester
VDAC: Mitochondrial voltage-dependent anion channel
ΔΨm: Mitochondrial membrane potential

## Acknowledgments

The authors thank Nathalie L’Heureux, Chantal St. Cyr, Ramona Löcker, Sylvia Metze, Simone Olesch, Dennis Hoffmann, Lu Yang, and the Imaging Center Campus Essen—University Duisburg Essen for excellent technical assistance, and Lucie Lefebvre for secretarial assistance.

## Novelty and Significance

### What Is Known?

- AF is associated with progressive remodeling of tissue properties, which makes the arrhythmia more persistent and resistant to therapy.
- There is evidence that disturbed mitochondrial function is implicated in AF.
- How mitochondria remodel as a function of time in AF, and the potential value of targeting mitochondrial changes in preventing AF-progression, are unknown.

### What New Information Does This Article Contribute?

- Mitochondria show a sequence of changes in response to AF, with initial adaptation followed by progressive deterioration in mitochondrial properties and function.
- Mitochondrial ROS production increases early in AF and continues to remain elevated over time; *in vitro* mitochondrial antioxidants attenuate AF-induced cellular remodeling.
- In vivo oral therapy with a mitochondrial-targeted drug suppresses AF-related adverse mitochondrial remodeling, deleterious changes in atrial cellular and electrical dysfunction and AF-progression, meriting consideration as a novel approach to treating AF.

