## Supplementary material for "Time-dependent Mitochondrial Remodeling in Experimental Atrial Fibrillation and Potential Therapeutic Relevance": Online Figure

**Online Figure S1**

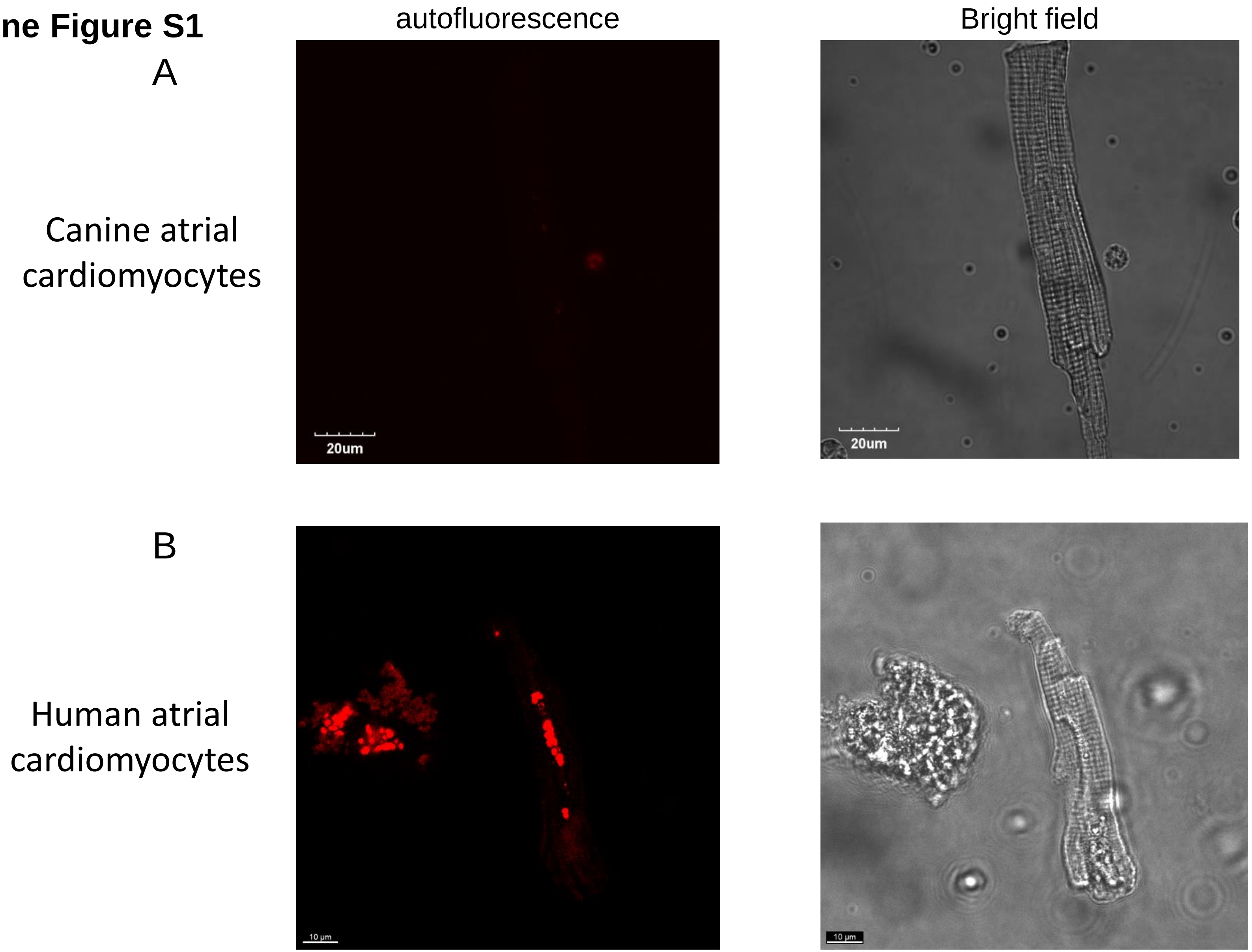


**Figure S1.** Examples of confocal microscopic observation of autofluorescence in canine (**A**) and human (**B**) atrial cardiomyocytes. Images were acquired at excitation wavelength of 500 nm. Human cardiomyocytes showed spontaneous autofluorescence.

### Online Figure S2

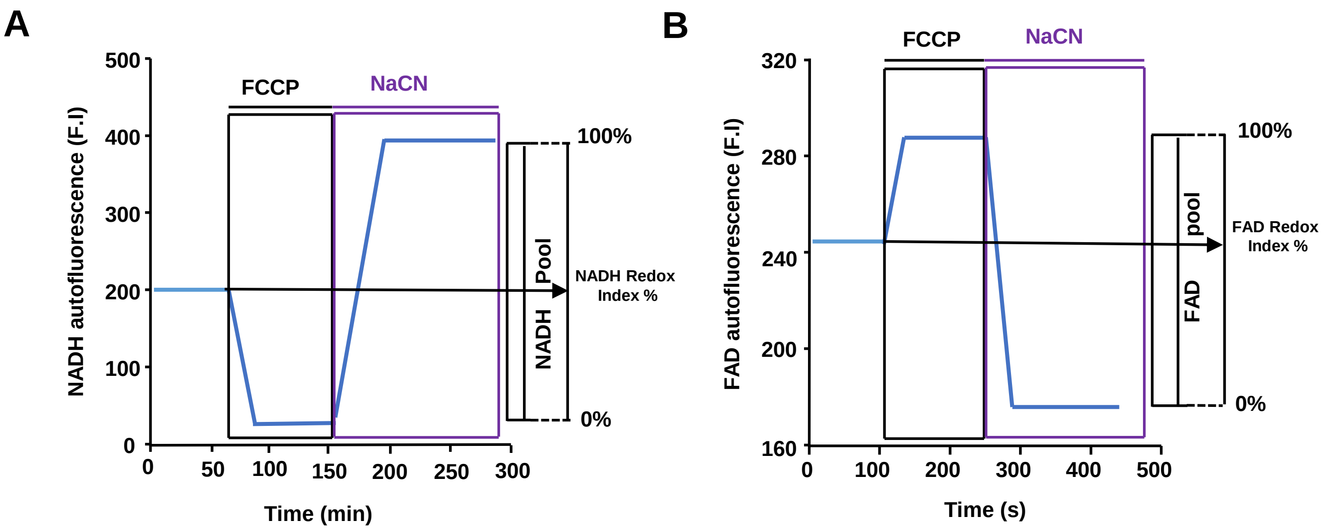


**Figure S2.** Schematic representation of the methods to analyze NADH (**A**) and FAD (**B**) in atrial cardiomyocytes (CMs). NADH redox indexes were obtained by measuring the initial NADH autofluorescence, followed by minimum NADH autofluorescence in the presence of the mitochondrial oxidative-phosphorylation uncoupler FCCP and then maximum NADH values in the presence of the NADH-oxidation blocker NaCN. FAD autofluorescence using confocal microscopy, FAD redox indexes were obtained under the same conditions, with a response reciprocal to that of NADH.

### Online Figure S3

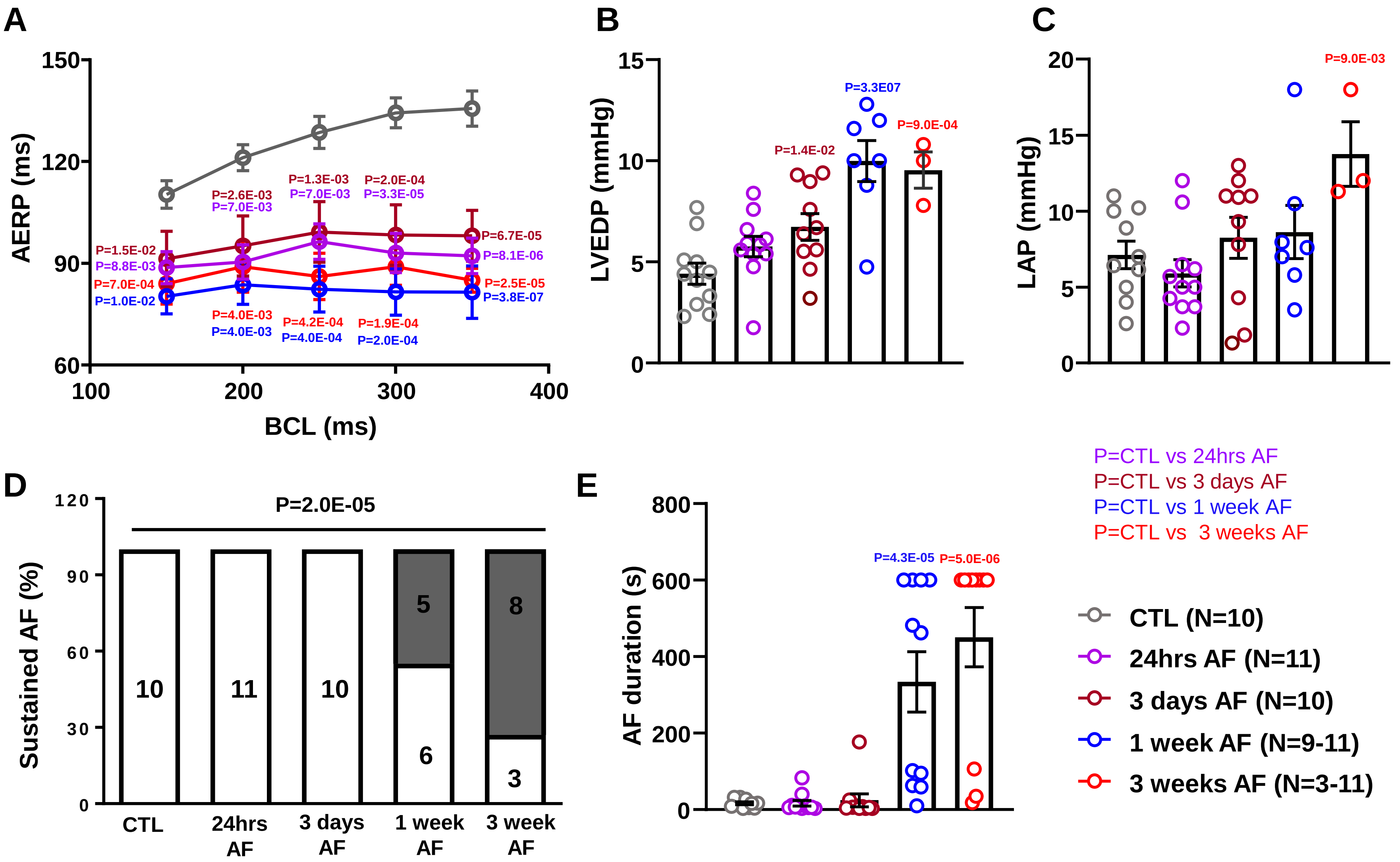


**Figure S3. A.** Mean ± SEM atrial effective refractory period (aERP; grey open circles: control (CTL); purple: 24-hour AF, dark red: 3-day AF, blue: 1 week AF, red: 3-weeks AF; BCL indicates basic cycle length); **B.** Mean ± SEM left ventricular end-diastolic pressure (LVEDP); **C.** Mean ± SEM left atrial pressure (LAP); **D.** incidence of sustained AF; **E.** Mean ± SEM duration of induced atrial fibrillation (AF) in CTL and AF dogs.

### Online Figure S4

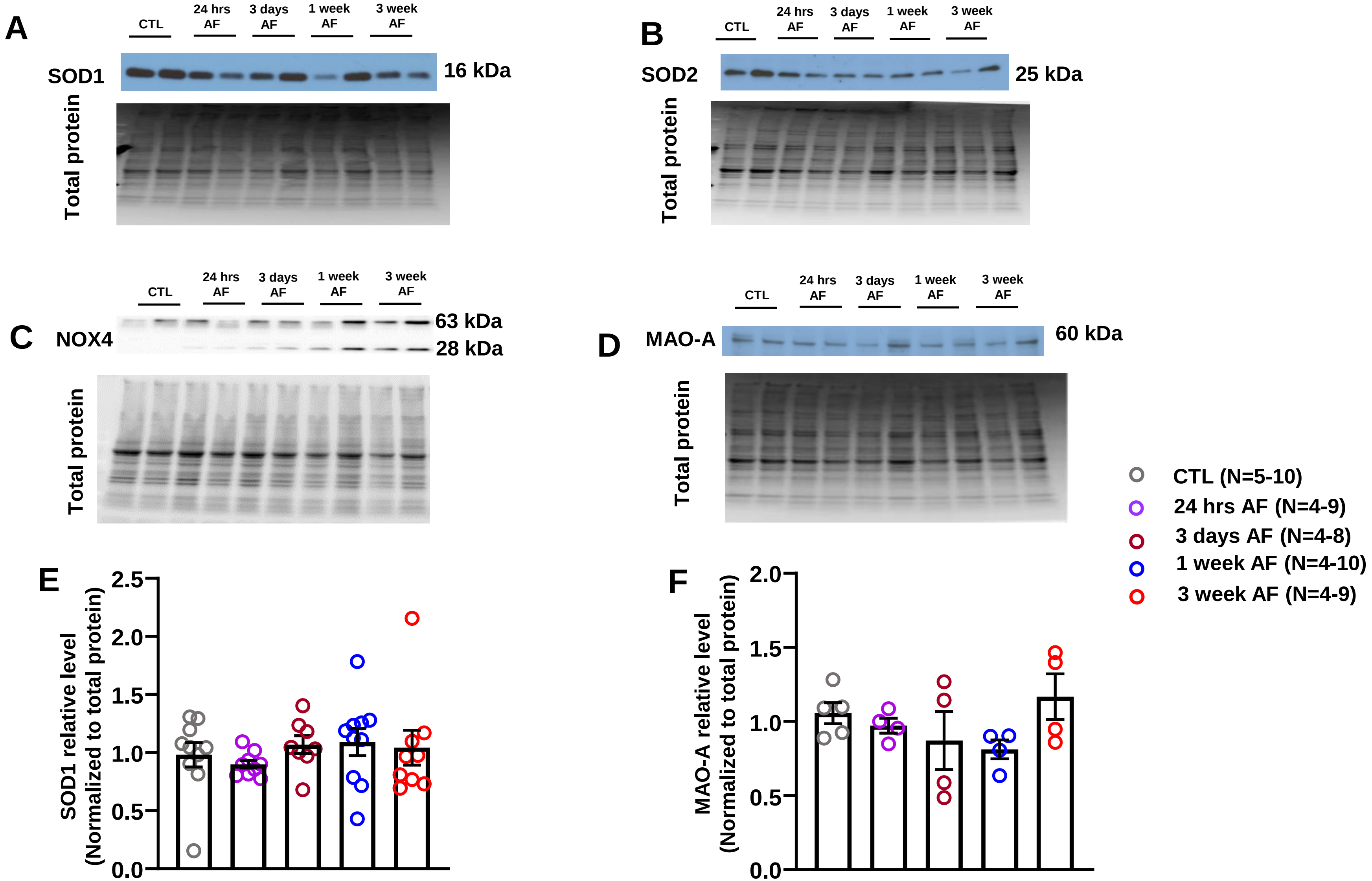


**Figure S4. A-D.** SOD1, SOD2, NOX4 and MAO-A immunoblot and total protein loading control images from CTL and AF atrial CMs; **E-F.** Mean±SEM SOD1 and MAO-A protein expression level.

### Online Figure S5

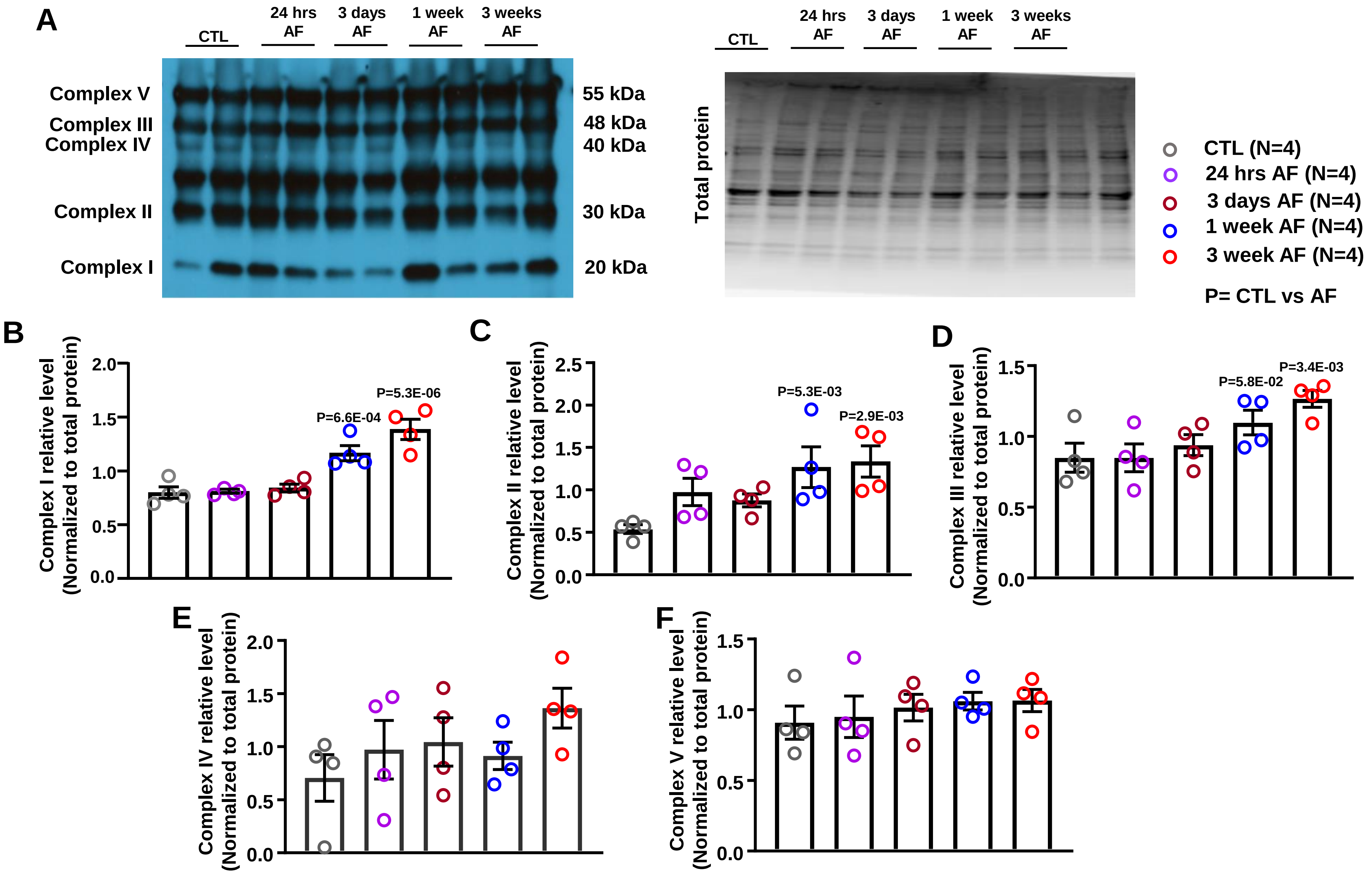


**Figure S5. A**. Western blot images of mitochondrial OXPHOS respiratory complex protein expression and total protein loading controls. Antibodies against proteins representing the five mitochondrial oxidative phosphorylation complexes were used simultaneously to examine the expression of mitochondrial proteins in CTL and AF atrial cardiomyocytes; **B-F**. Mean ± SEM protein expression level.

### Online Figure S6

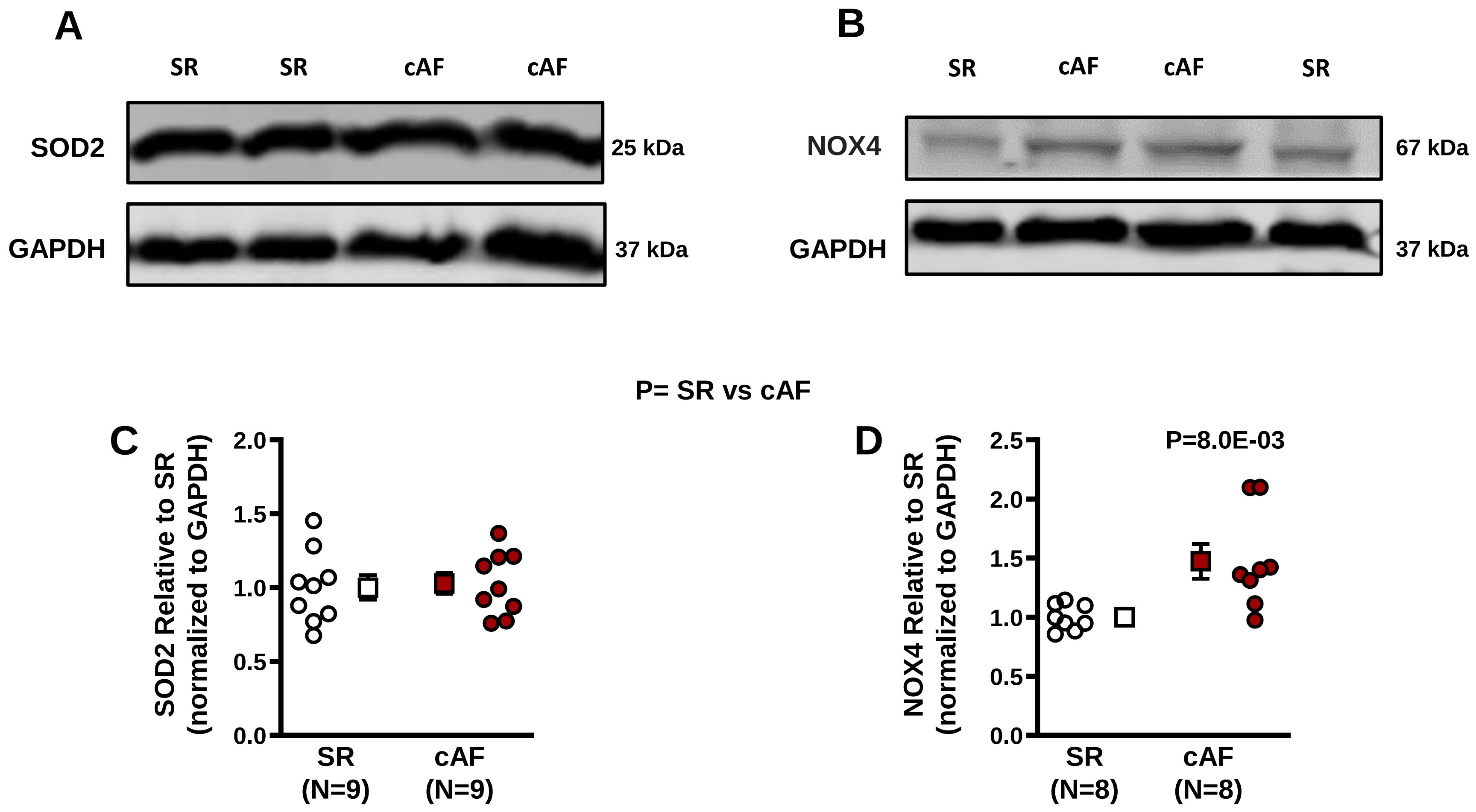


**Figure S6. A, B.** SOD2, NOX4 and GAPDH immunoblot band images from SR and cAF patient samples; **C, D.** Mean±SEM SOD2 and NOX4 protein expression levels.

### Online Figure S7

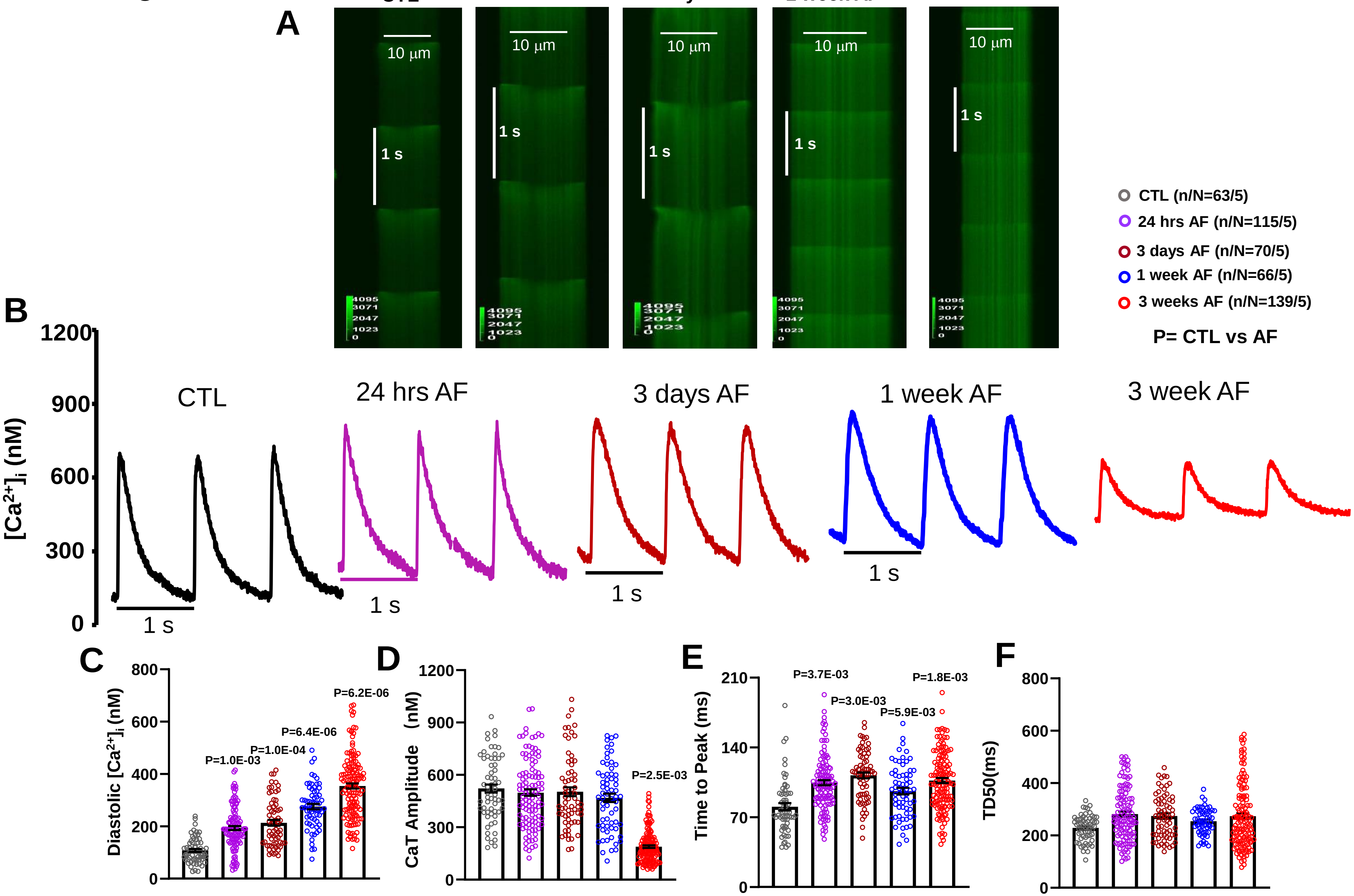


**Figure S7.A-B**. Confocal line scan of cytosolic [$Ca^{2+}$] and corresponding [$Ca^{2+}$] transient (CaT) of intracellular cytosolic [$Ca^{2+}$] transients (CaT) in 1Hz stimulated atrial CMs; **C-F**. Mean ± SEM cytosolic [$Ca^{2+}$] transient properties. TD50=time for 50% decay of CaT.

#### Online Figure S8

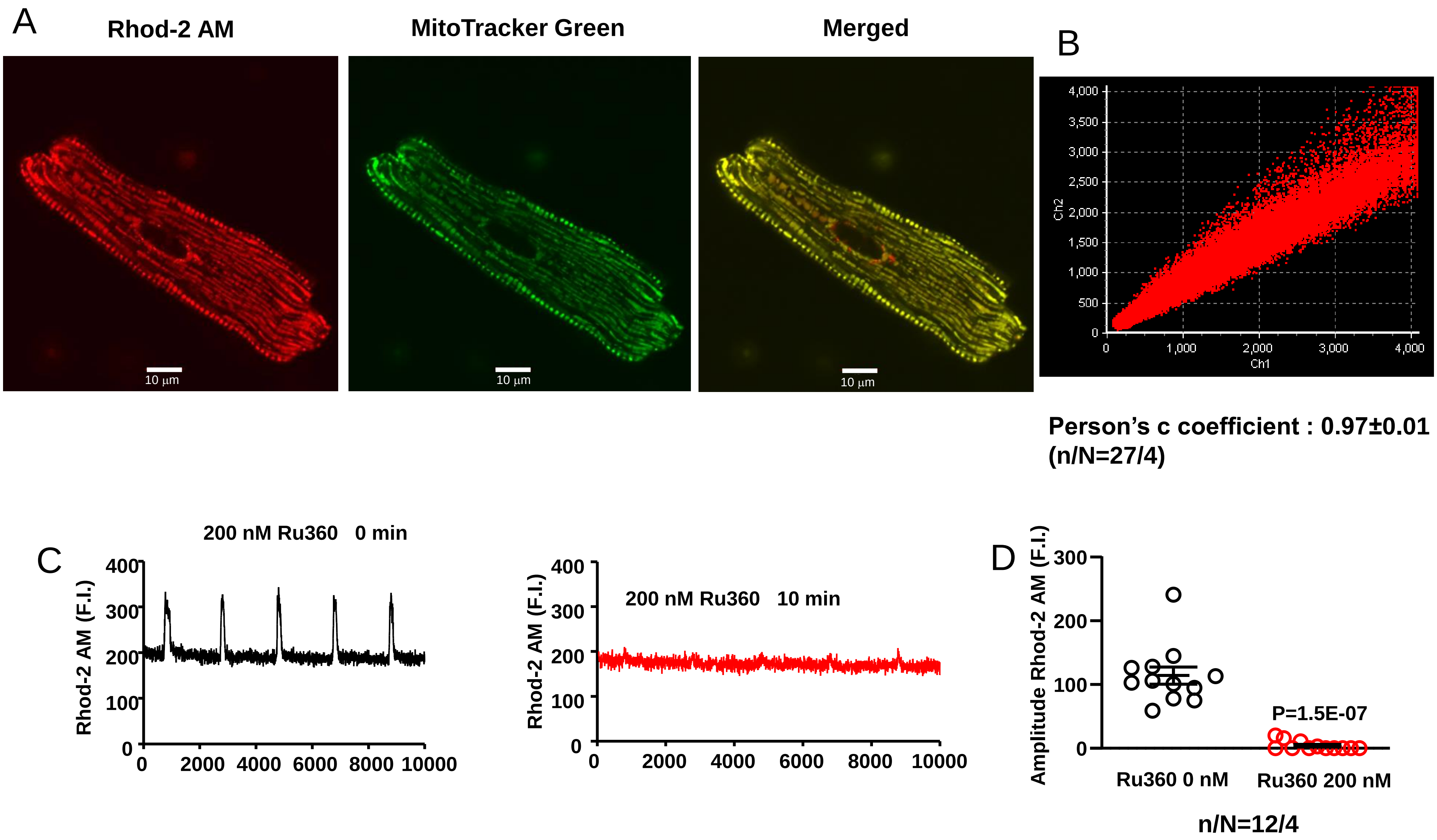


**Figure S8. A.** Examples of microscopic cellular images of staining from Rhod-2 AM and MitoTracker Green, as well as merged image. B. Pearson's c-coefficient for co-localization of Rhod-2 and MitoTracker staining, indicating localization to mitochondria. C. $[Ca^{2+}]_{mito}$ transient recordings before and exposure to 200 nM Ru360 for 10 min. D. Rhod-2 fluorescence amplitudes before and after Ru360 exposure in 12 cells from 4 dogs.

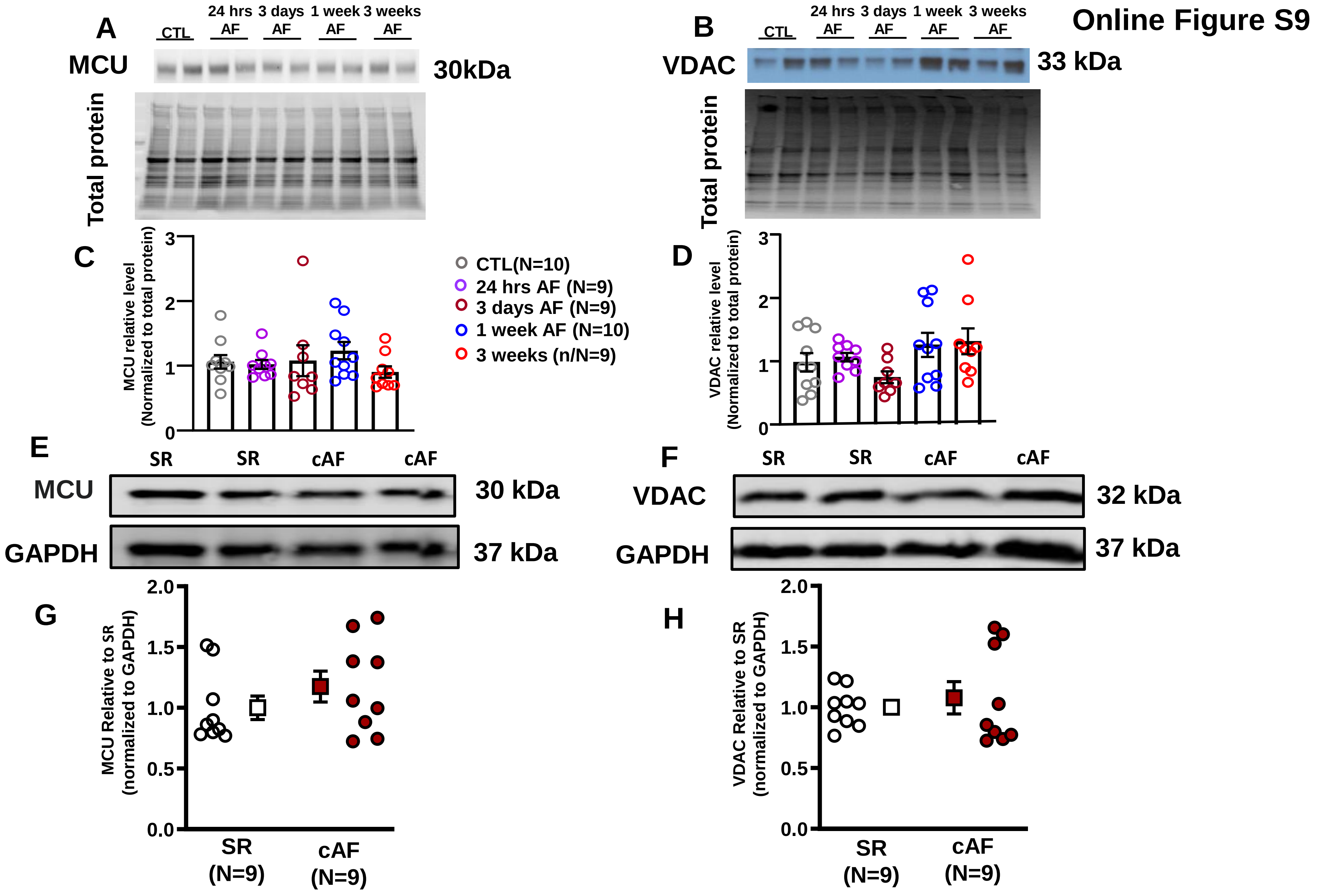


**Figure S9. A, B.** Western blot images of MCU, VDAC and total protein from canine CTL and AF atrial CMs; **C, D.** Mean±SEM protein expression of MCU and VDAC; **E, F.** Western blot images of MCU, VDAC and GAPDH from SR and cAF patient samples; **G, H.** Mean±SEM MCU and VDAC protein expression level.

### Online Figure S10

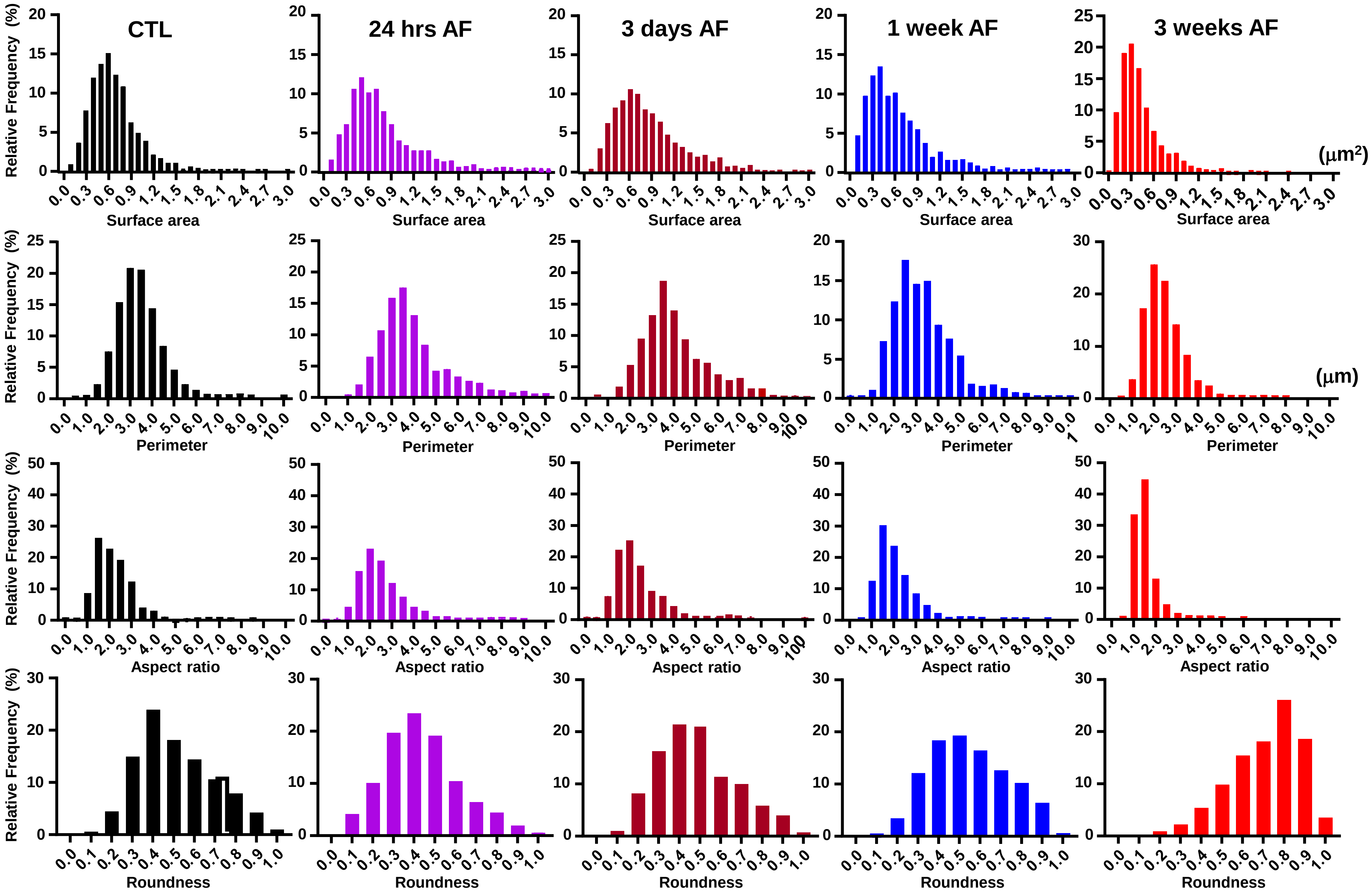


**Online Figure S10 .** Histograms showing distribution frequency (% total mitochondria) of mitochondrial surface area, perimeter, aspect ratio and roundness from CTL and AF left atria.

### Online Figure S11

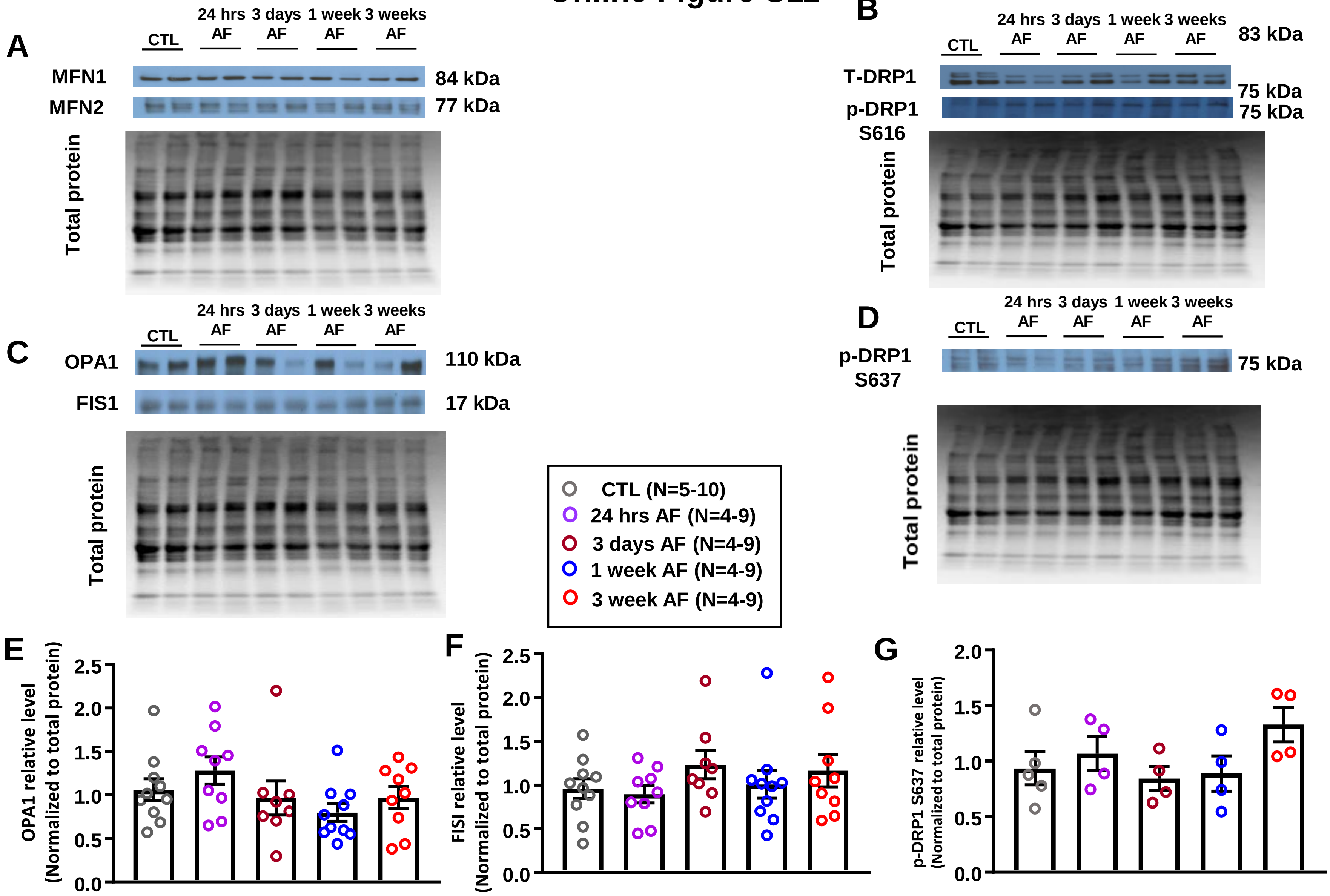


**Figure S11. A.** Western blot images of MFN1, MFN2 and total protein from CTL and AF canine ACMs; **B.** Western blot images of T-DRP1, p-DRP1 S616 and total protein from CTL and AF canine ACMs; **C, D.** Western blot images of OPA1, FIS1, p-DRP1 S637 and total protein from CTL and AF canine ACMs; **E-G.** Mean ± SEM OPA1, FIS1 and p-DRP1 S637 protein expression level in CTL and AF canine ACMs.

### Online Figure S12

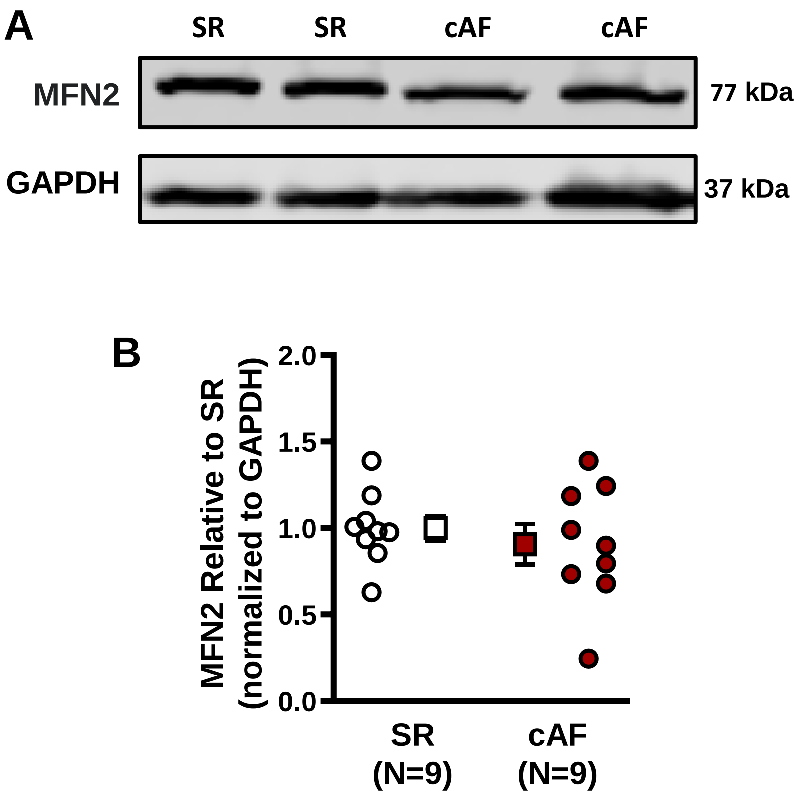


**Figure S12. A.** Western blot images of MFN2 and GAPDH from SR and cAF patient samples; **B**. Mean±SEM MFN2 protein expression level.

#### Online Figure S13

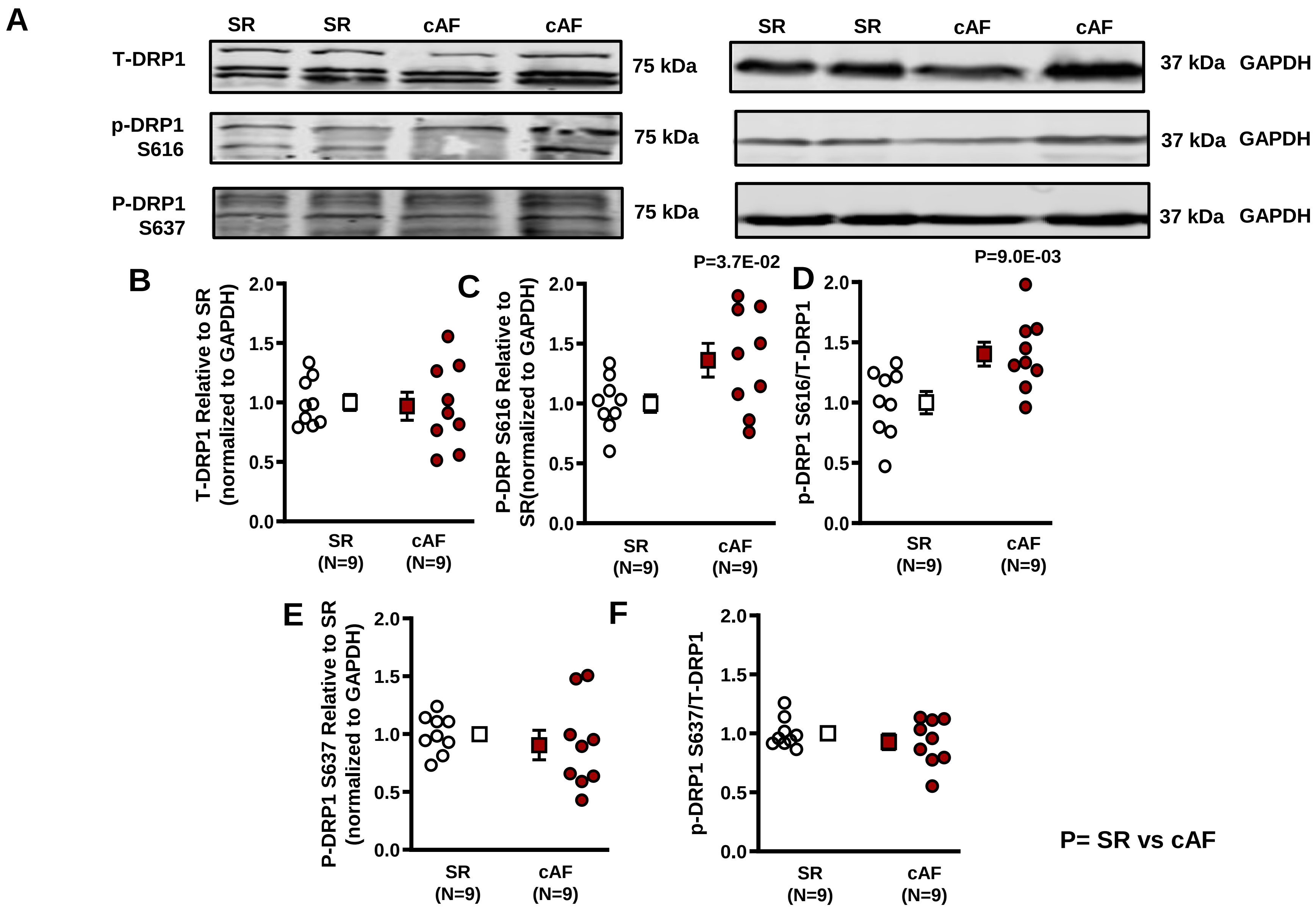


**Figure S13. A**. Western blot images of T-DRP1, p-DRP1 S616, p-DRP1 S637 and GAPDH from SR and cAF patient samples; **B, C.** Mean±SEM T-DRP1 and p-DRP1 S616 protein expression level; **D.** p-DRP1 S616/T-DRP1 ratio; **E,** Mean±SEM p-DRP1 S637 protein expression level; **F.** p-DRP1 S637/T-DRP1 ratio.

### Online Figure S14

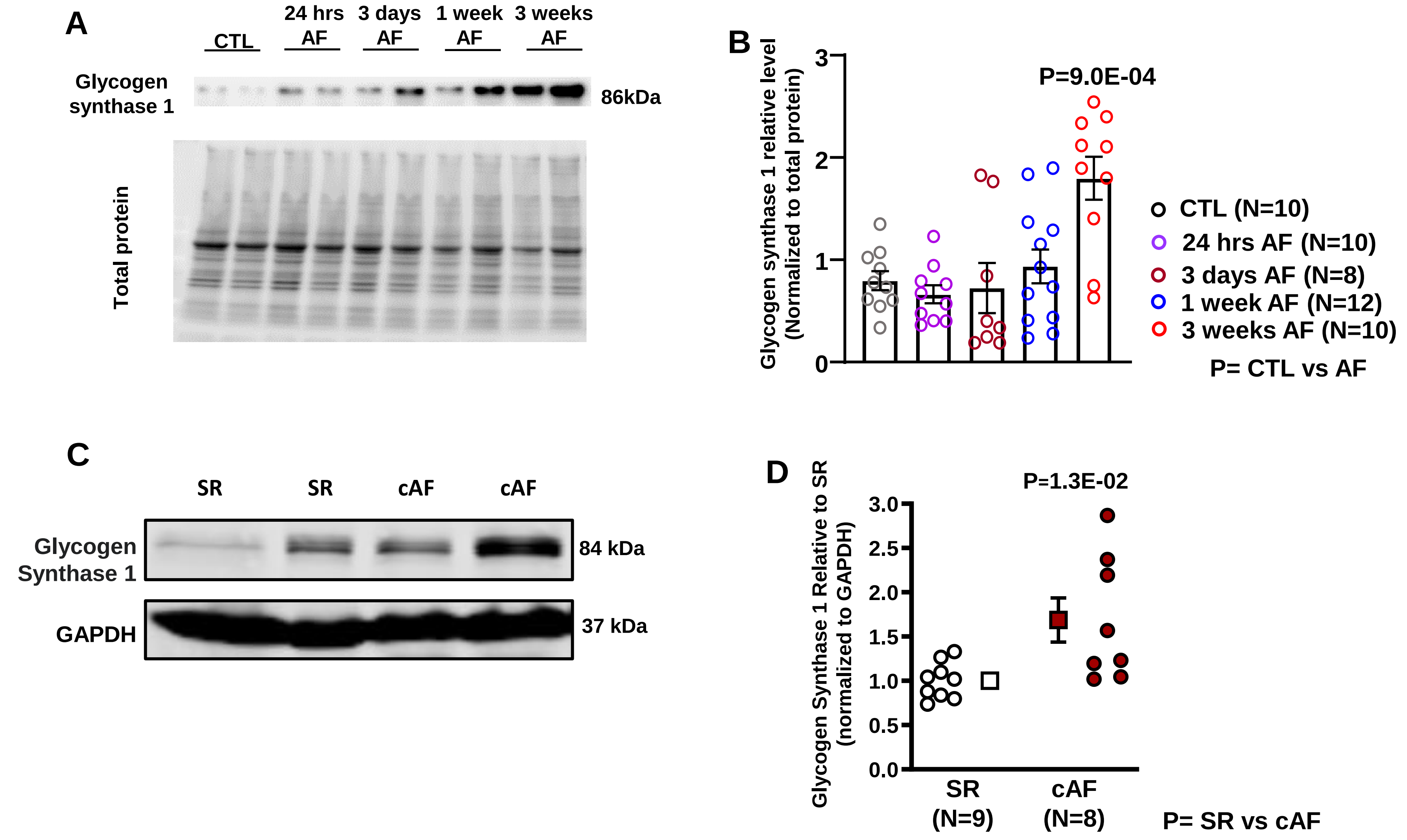


**Figure S14. A.** Western blot images of Glycogen Synthase1 and total protein from CTL and AF atrial CMs; **B**. Mean±SEM Glycogen Synthase1 protein expression level; **C.** Western blot images of Glycogen Synthase1 and GAPDH from SR and cAF patient samples; **D.** Mean±SEM Glycogen Synthase1 protein expression level.

### Online Figure S15

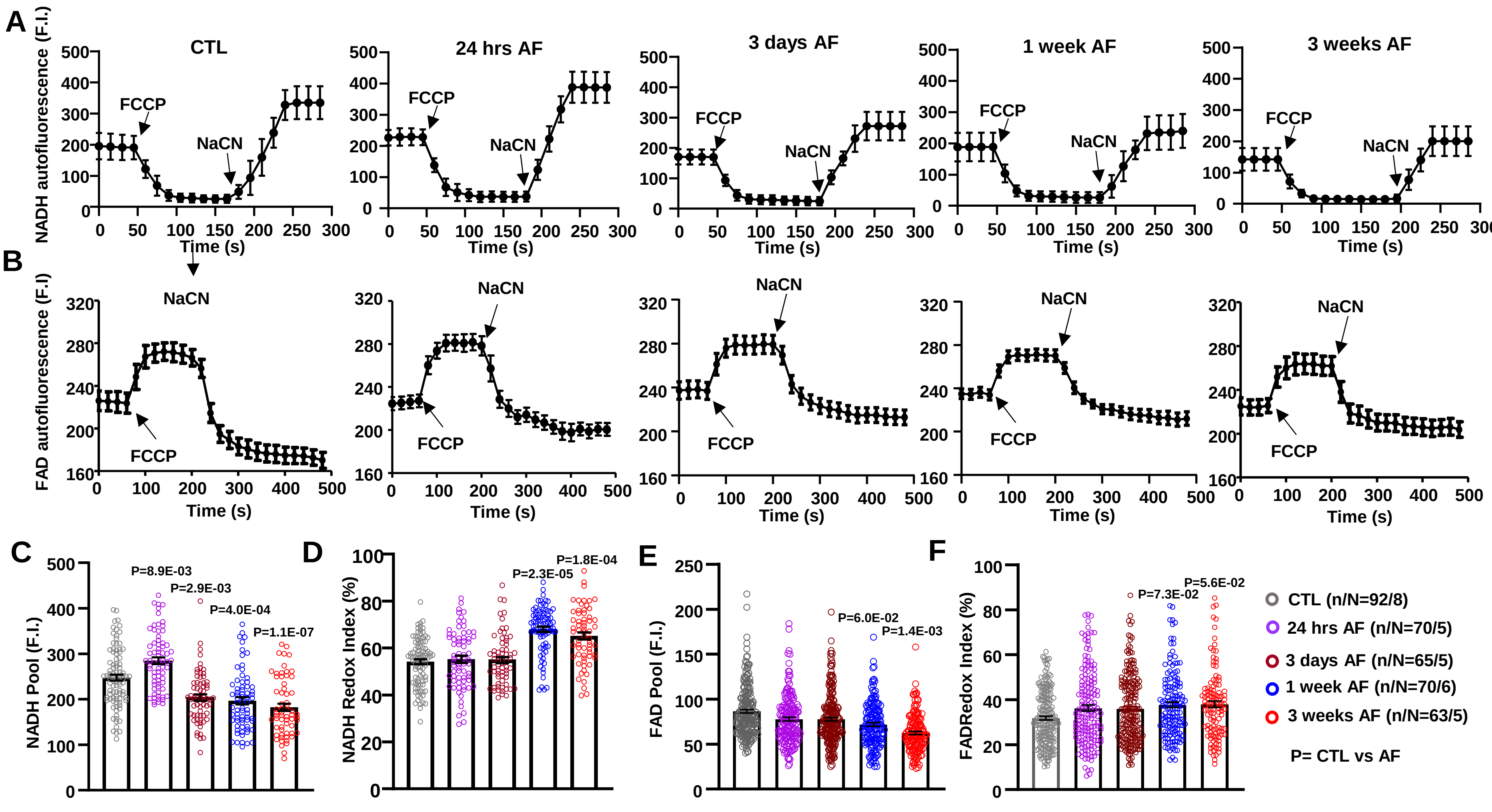


**Figure S15. A, B.** Representative average values for NADH and FAD autofluorescence from CTL and AF CMs obtained with protocols in Figure S2; **C, D.** Mean±SEM mitochondrial NADH pool and NADH redox index from CTL and AF atrial CM; **E, F.** Mean±SEM mitochondrial FAD pool and FAD redox index from CTL and AF atrial CMs.

### Online Figure S16

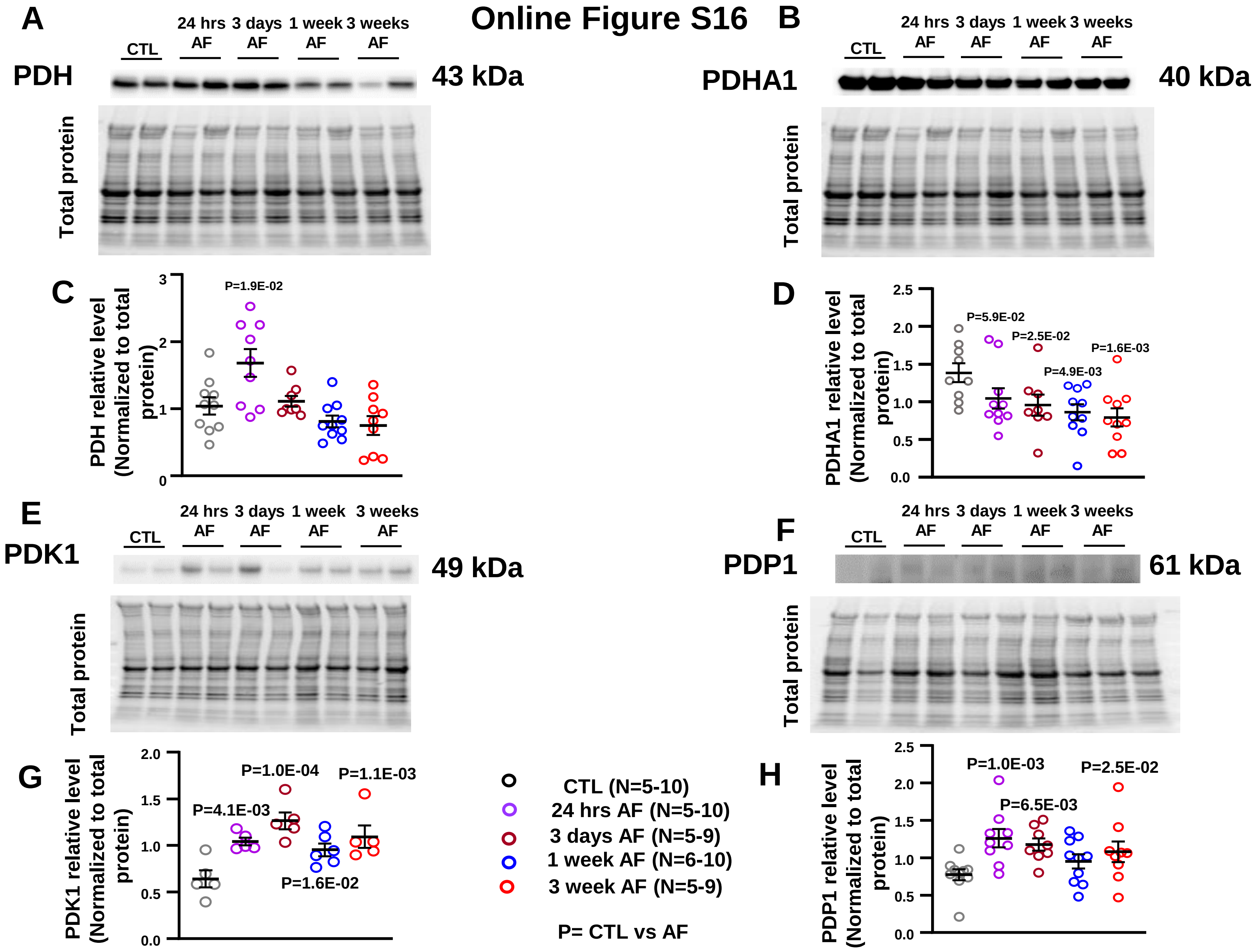


**Figure S16. A, B.** Western blot images of PDH, PDHA1 and total protein from CTL and AF canine atrial CMs; **C, D.** Mean±SEM PDH and PDHA1 protein expression level; **E, F.** Western blot images of PDK1, PDP1 and total protein from CTL and AF atrial CMs; **G, H.** Mean±SEM PDK1 and PDP1 protein expression level.

### Online Figure S17

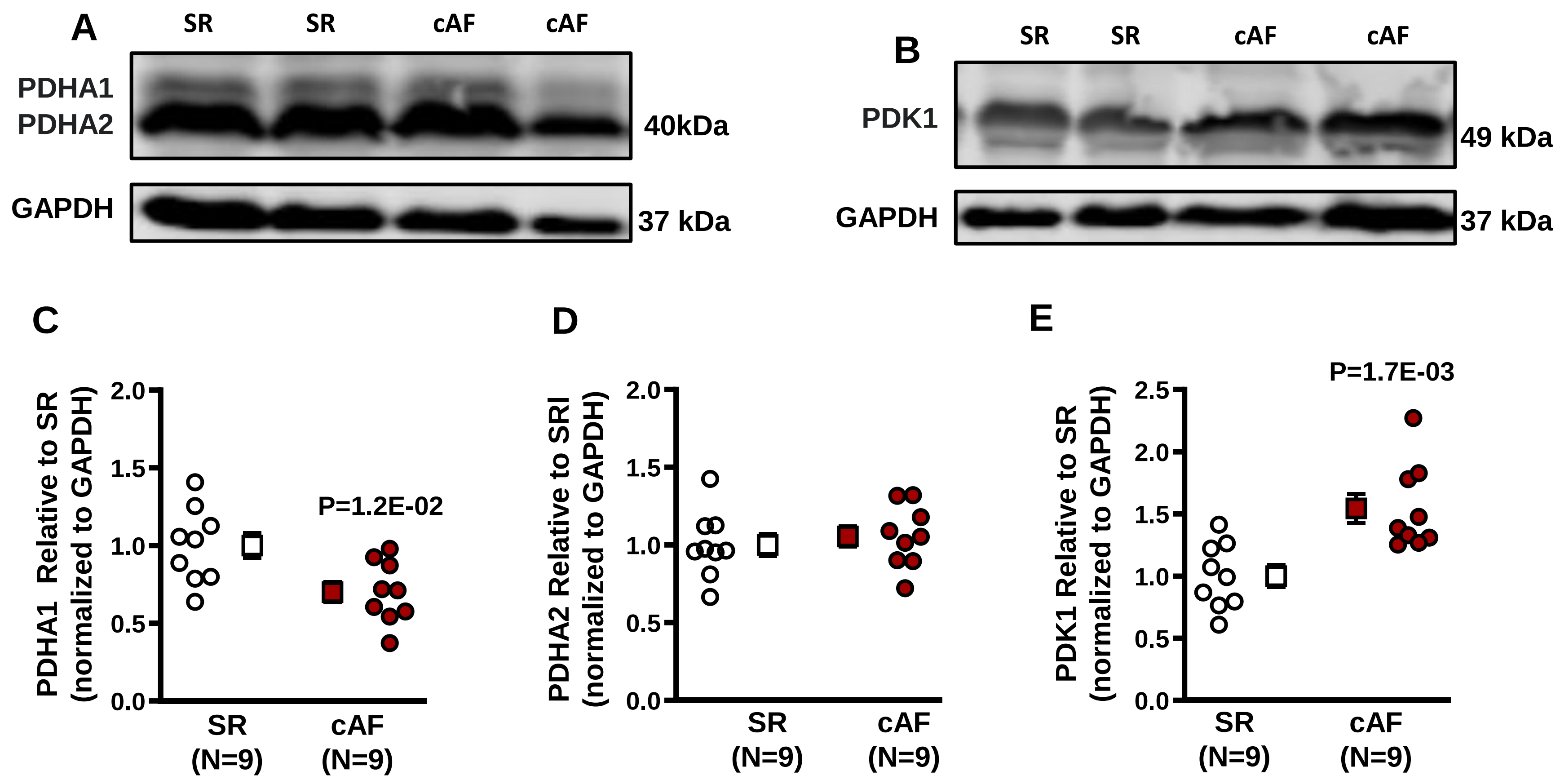


**Figure S17. A.** Western blot images of PDHA1, PDHA2 and GAPDH from SR and cAF patient samples; **B.** Western blot images of PDK1 and GAPDH from SR and cAF patient samples; **C-E.** Mean±SEM PDHA1, PDHA2 and PDK1protein expression level.

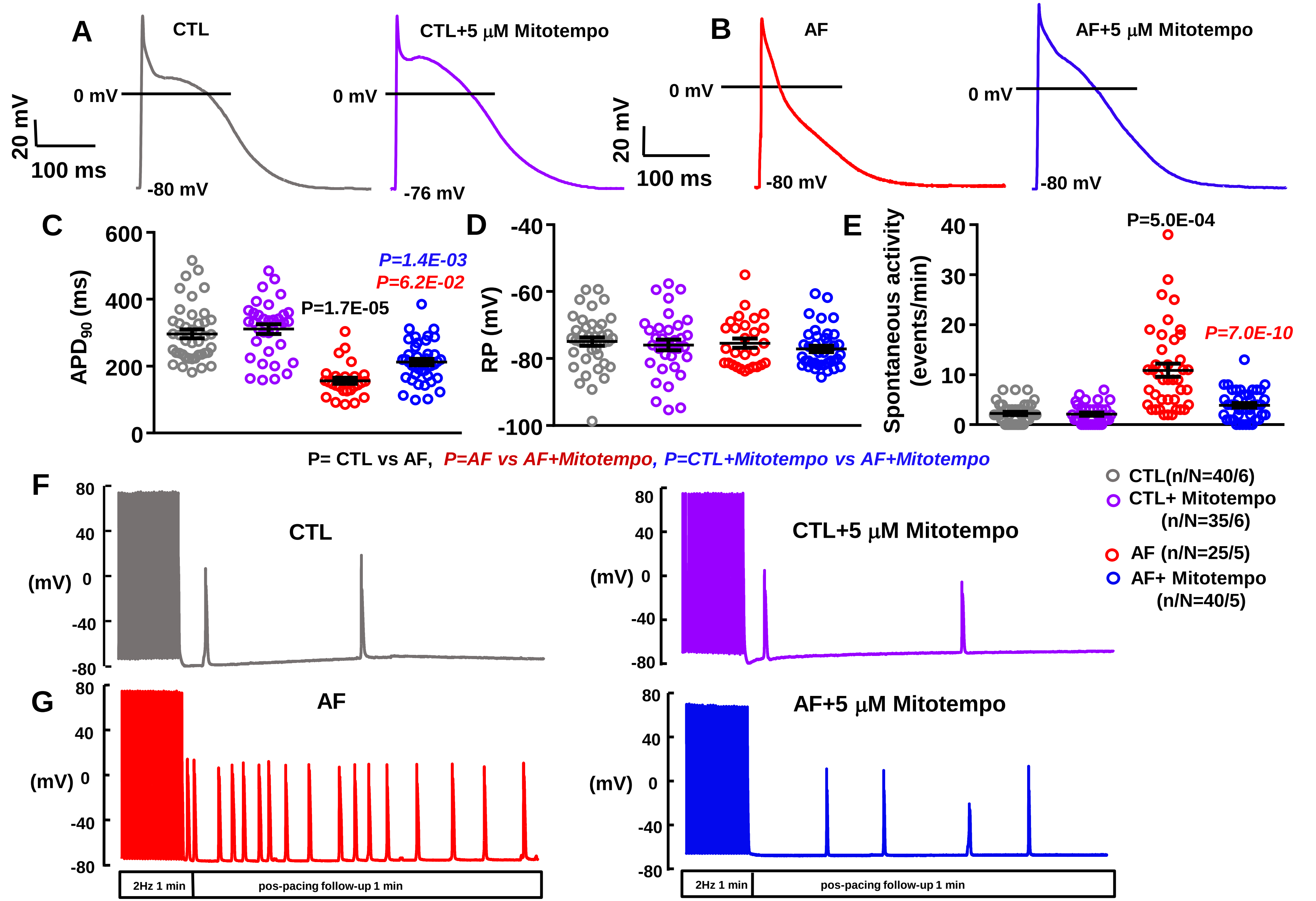


**Online Figure S18. A, B.** Action potential recordings at 1-Hz from canine CTL and AF (1-wk) canine atrial cardiomyocytes with/without MitoTempo; **C, D.** Mean±SEM $APD_{90}$ and RP; **E.** Frequency of spontaneous activity following $Ca^{2+}$-loading at 2 Hz; **F, G.** Triggered APs, after $Ca^{2+}$ loading by 1 minute of pacing at 2 Hz from canine CTL and AF atrial CMs with/without MitoTempo.

#### Online Figure S19

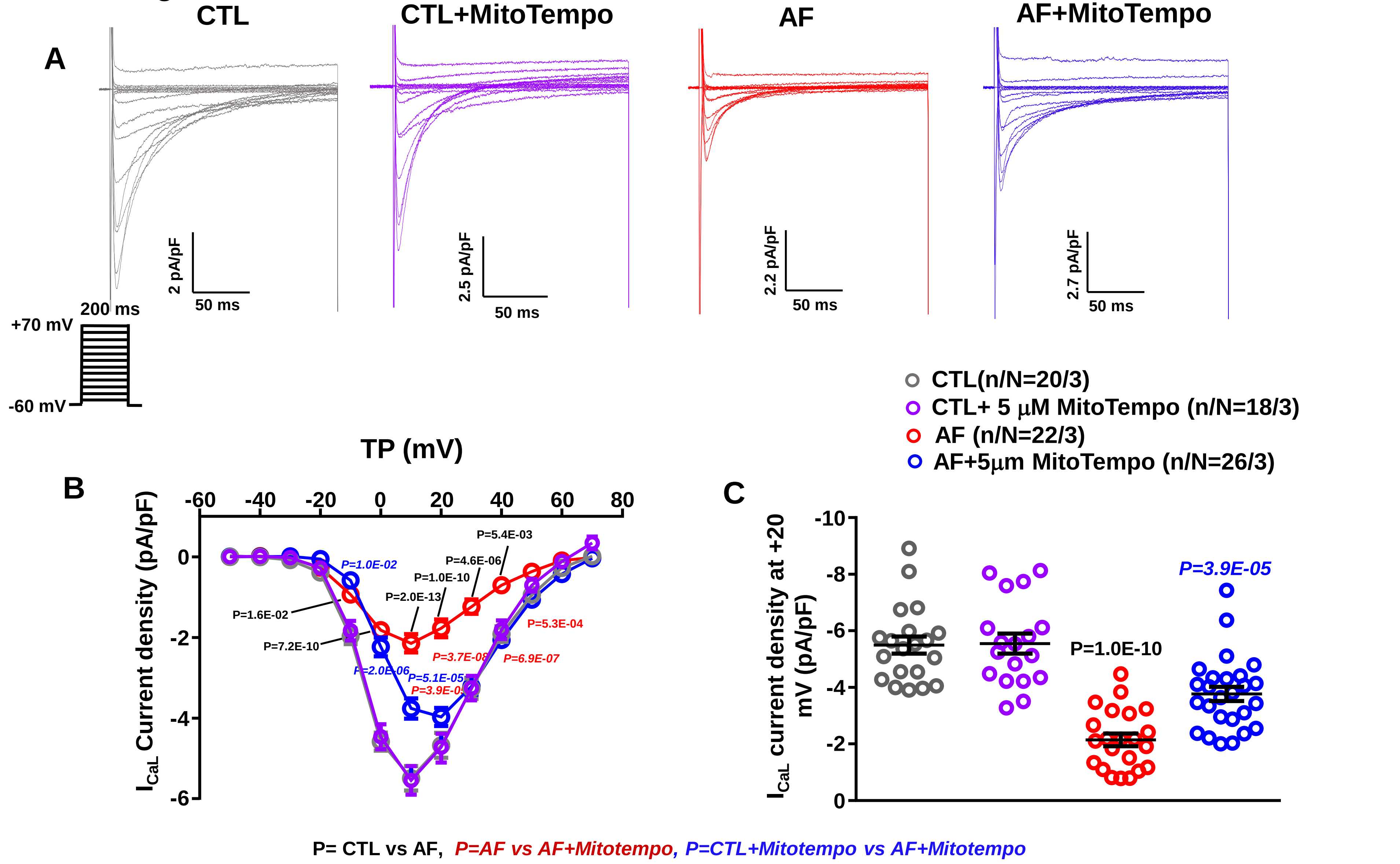


**Figure S19. A.** $I_{CaL}$ recordings at 0.1 Hz from canine CTL and AF (1-wk) atrial cardiomyocytes with/without MitoTempo; **B.** Current-density voltage relation of $I_{CaL}$ from CTL and AF atrial CMs**; C.** $I_{CaL}$ current densities at +20 mV from canine CTL and AF atrial CMs.

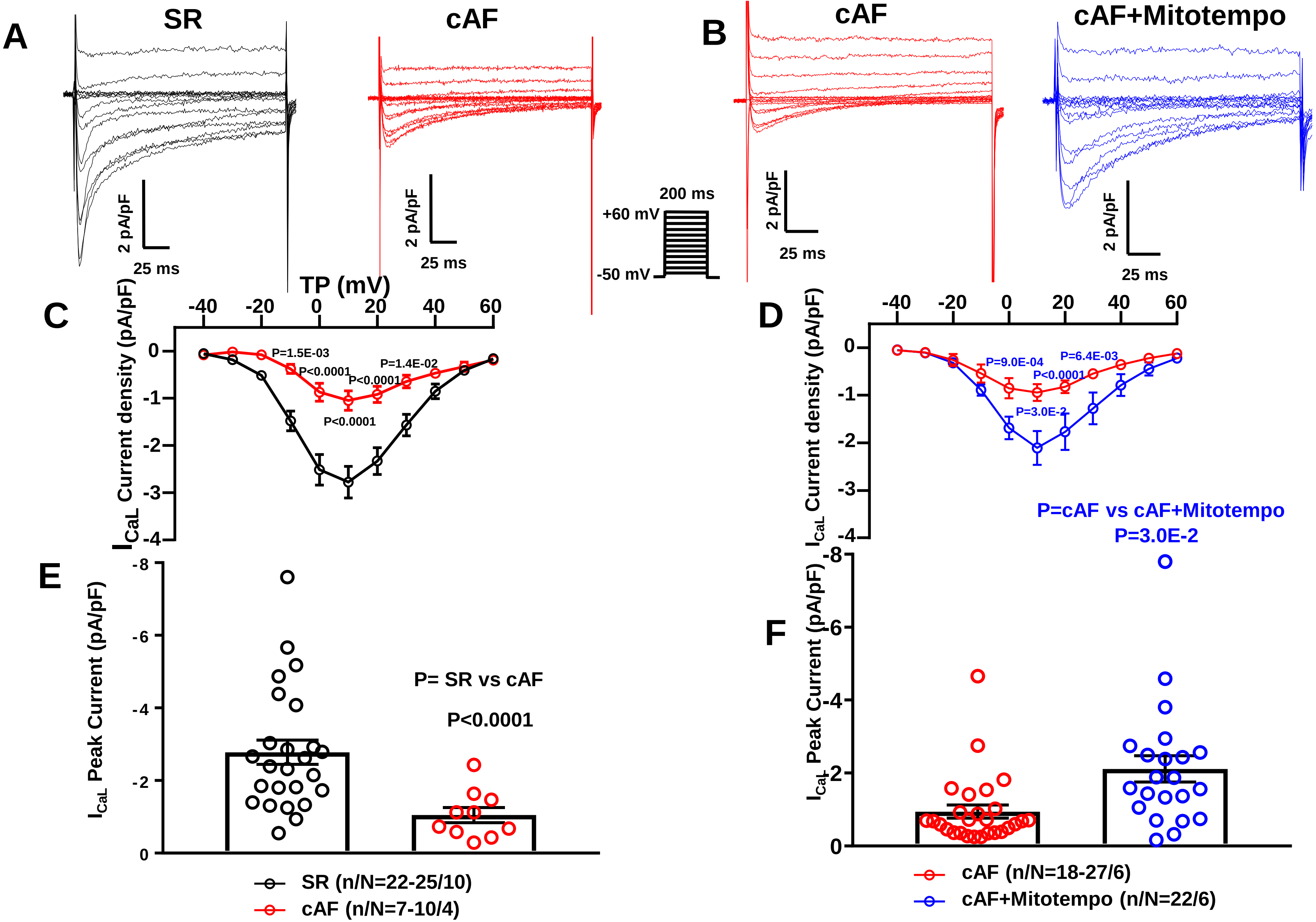


**Online Figure S20. A.** $I_{CaL}$ recordings at 0.1 Hz from SR and cAF patient atrial CMs; **B.** $I_{CaL}$ recordings at 0.1 Hz from cAF and cAF+MitoTempo patient atrial CMs; **C.** Current-density voltage relation of $I_{CaL}$ from SR and cAF patient atrial CMs; **D.** Current-density voltage relation of $I_{CaL}$ from cAF and cAF+MitoTempo patient atrial CMs; **E.** $I_{CaL}$ peak current densities at +10 mV from SR and cAF atrial CMs; **F.** $I_{CaL}$ peak current densities at +10 mV from cAF and cAF+ MitoTempo atrial CMs.

### Online Figure S21

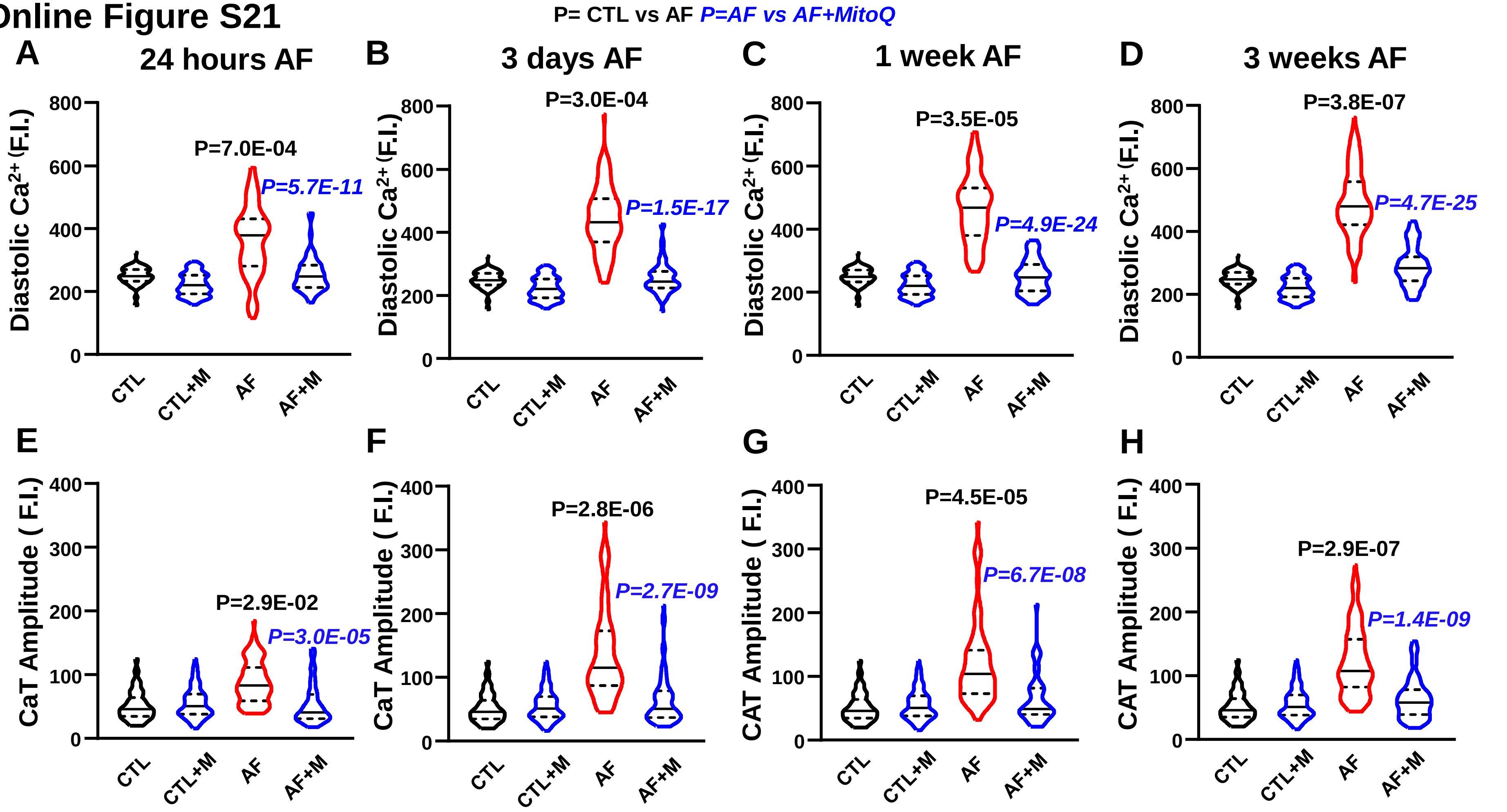


**CTL(n/N)=98/5**
**CTL+MitoQ(n/N)=109/5**
**AF(n/N)=95/5**
**AF+MitoQ(n/N)=58/5**

**CTL(n/N)=98/5**
**CTL+MitoQ(n/N)=109/5**
**AF(n/N)=82/5**
**AF+MitoQ(n/N)=70/5**

**CTL(n/N)=98/5**
**CTL+MitoQ(n/N)=109/5**
**AF(n/N)=76/5**
**AF+MitoQ(n/N)=46/5**

**CTL(n/N)=98/5**
**CTL+MitoQ(n/N)=109/5**
**AF(n/N)=104/5**
**AF+MitoQ(n/N)=62/5**

**Figure S21. A-D.** Mean±SEM diastolic $[Ca^{2+}]_{mito}$ from CTL and AF atrial CMs with/without 200 nM MitoQ; **E-H.** Mean±SEM $[Ca^{2+}]_{mito}$ transient amplitude in CTL and AF atrial CMs with/without 200 nM MitoQ. n/N=cells/dogs.

### Online Figure S22

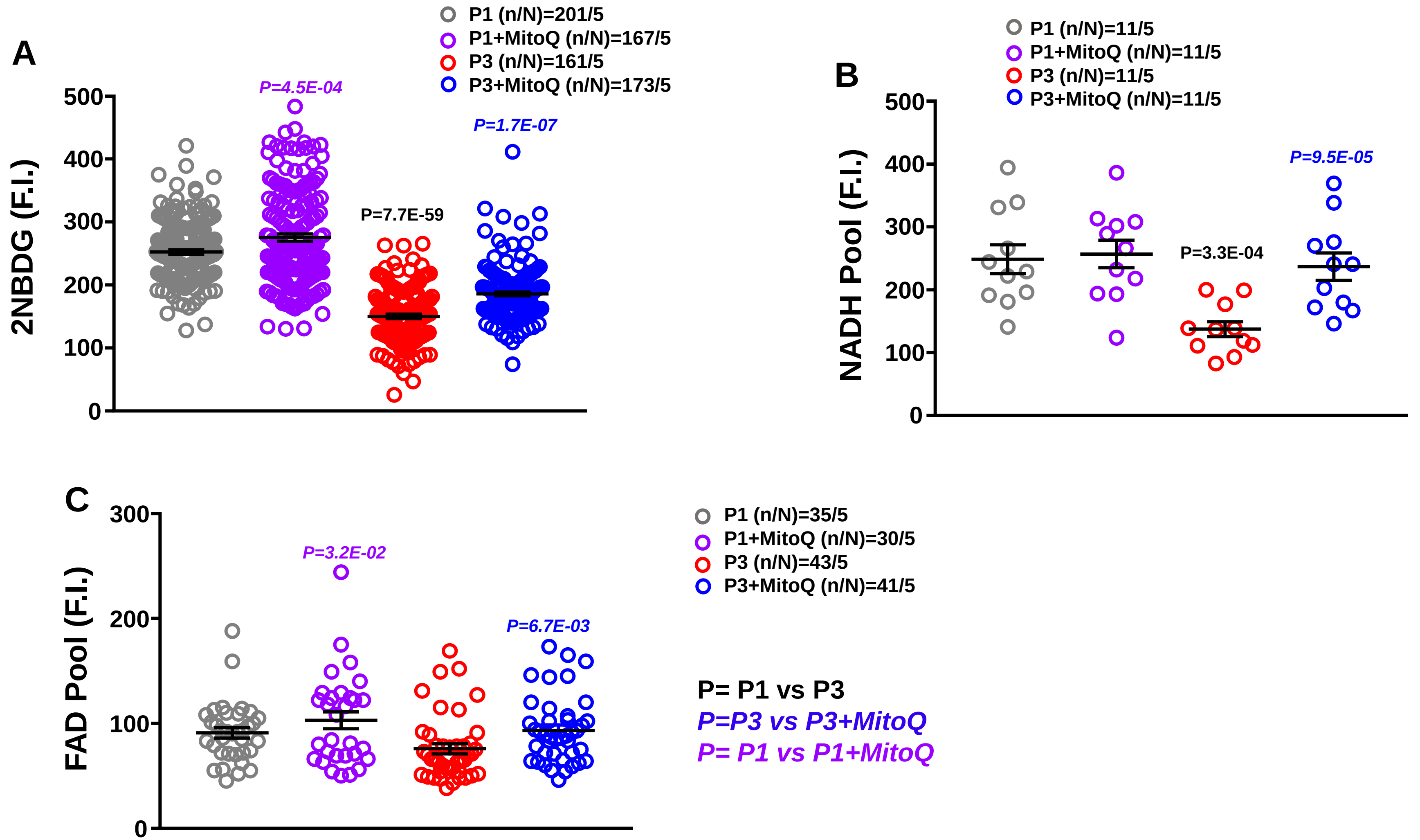


**Figure S22. A**. Mean±SEM 2-NBDG fluorescence intensity from P1 and P3 atrial CMs with/without MitoQ; **B.** Mean±SEM mitochondrial NADH pool from P1 and P3 atrial CMs with/without MitoQ; **C.** Mean±SEM mitochondrial FAD pool from P1 and P3 atrial CMs with/without MitoQ. Individual-cell values are shown along with Mean±SEM. n/N=cells/dogs.

### Online Figure S23

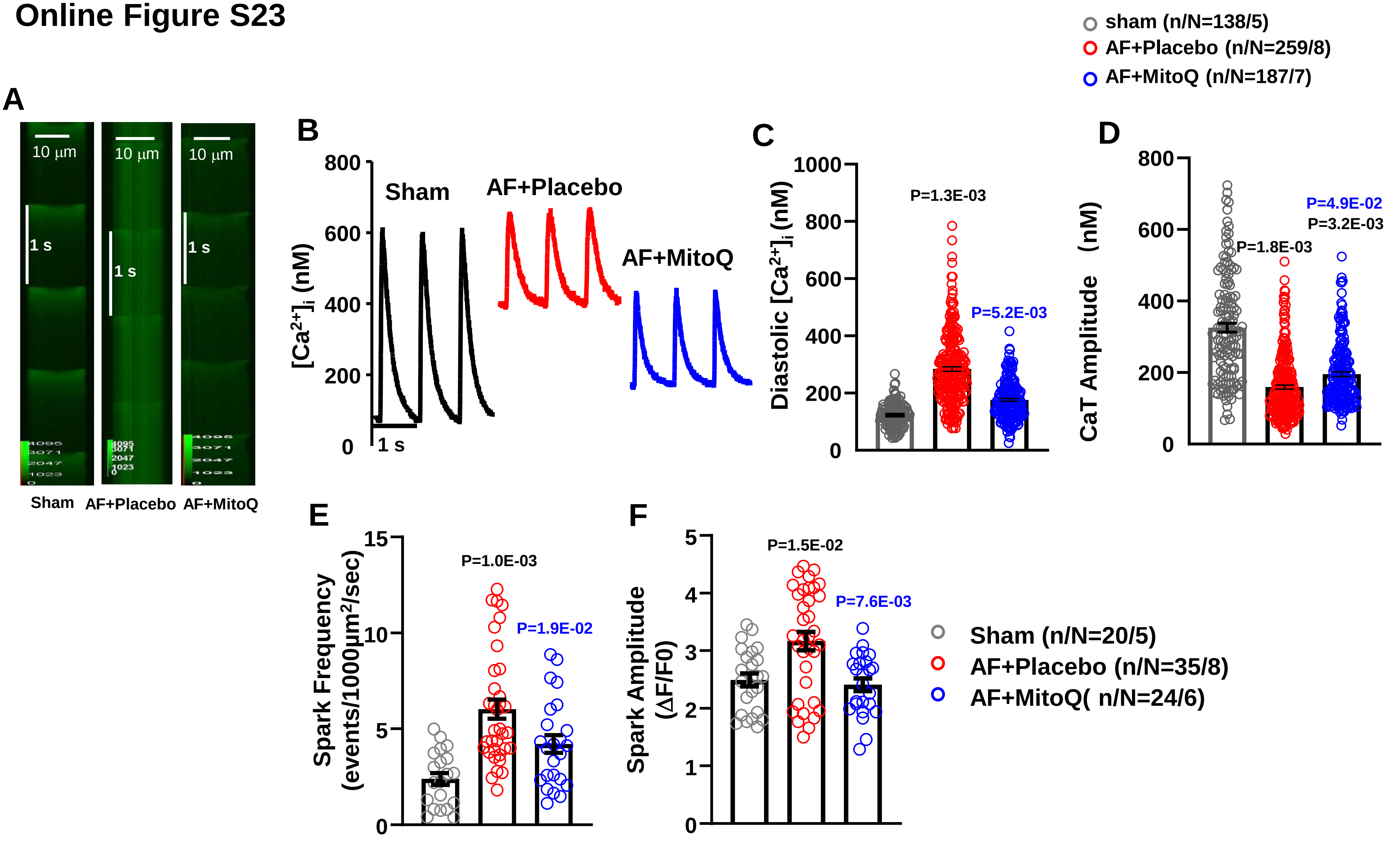


**Figure S23. A.** Confocal line scan of cytosolic [$Ca^{2+}$] and corresponding [$Ca^{2+}$] transient (CaT) recordings; **B.** Diastolic [$Ca^{2+}$]; **C.** CaT amplitude; **D, E.** $Ca^{2+}$ spark frequency (D) and amplitude (E). Data points in B-D are results from individual cells from 5, 8 and 7 dogs for Sham, AF+Placebo and AF+MitoQ respectively.

#### Online Figure S24

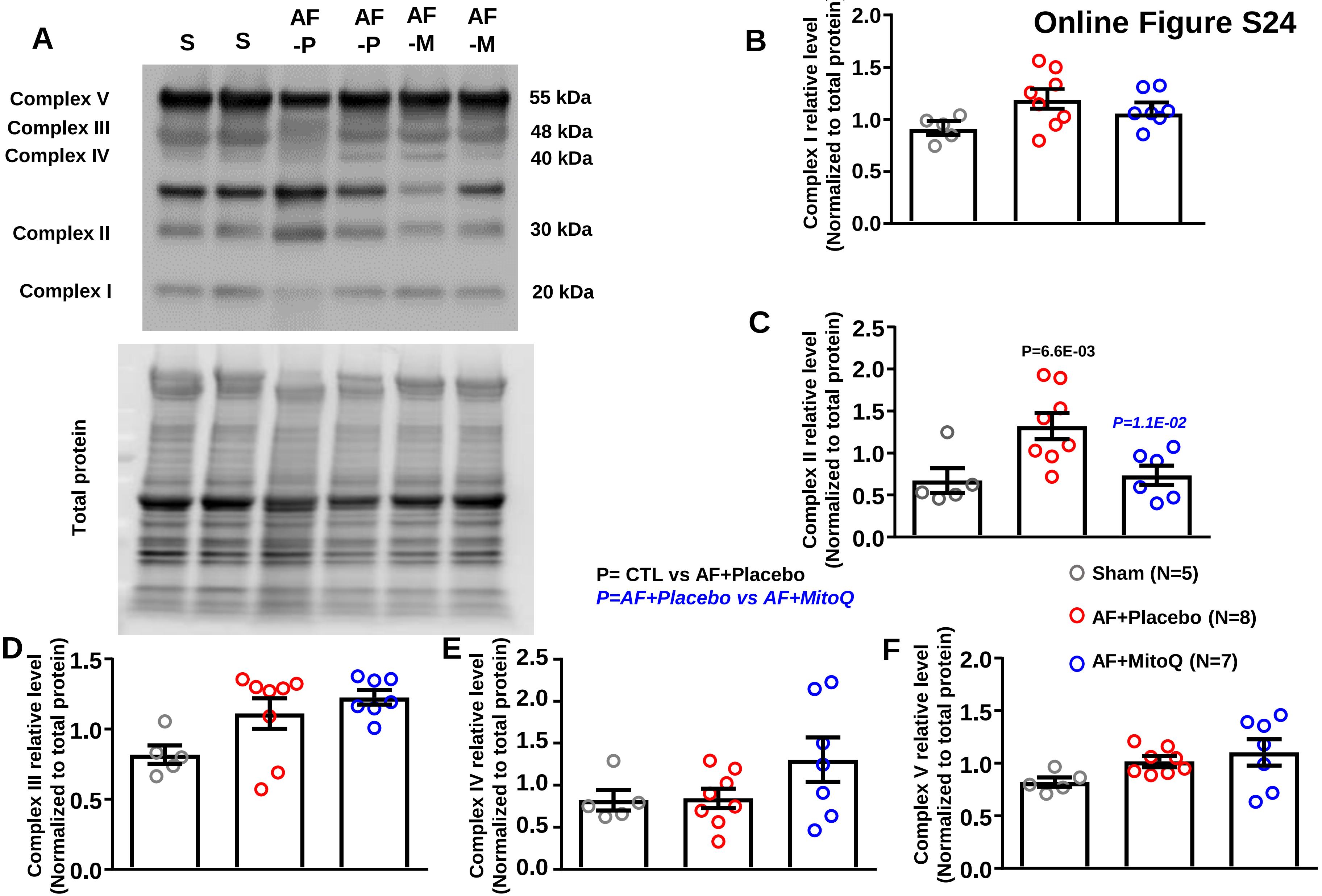


**Figure S24. A.** Western blot images of mitochondrial respiratory chain complex protein levels and total protein from sham, AF, AF+MitoQ canine atrial CMs. An antibody cocktail against proteins representing the five mitochondrial oxidative phosphorylation complexes was used to examine the expression of mitochondrial proteins in canine atrial cardiomyocytes from Sham, AF+Placebo and AF+MitoQ groups; **B-F**. Individual-data point and mean±SEM protein expression level.

#### Online Figure S25

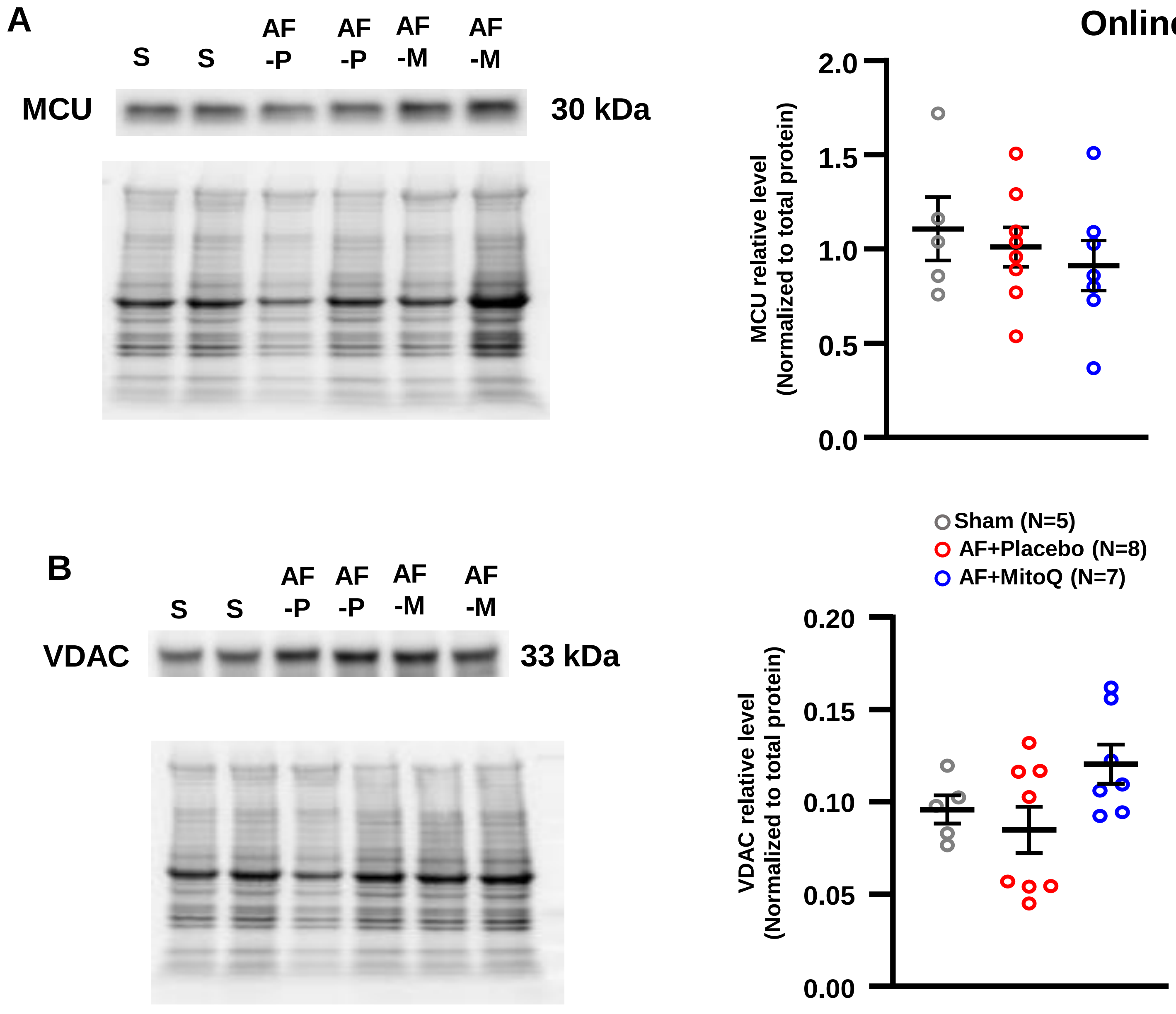


**Figure S25. A, B.** Western blot images of MCU, VDAC and total protein from Sham, AF+Placebo and AF+MitoQ atrial CMs, along with individual-sample (each sample is from a separate dog) and mean±SEM data.

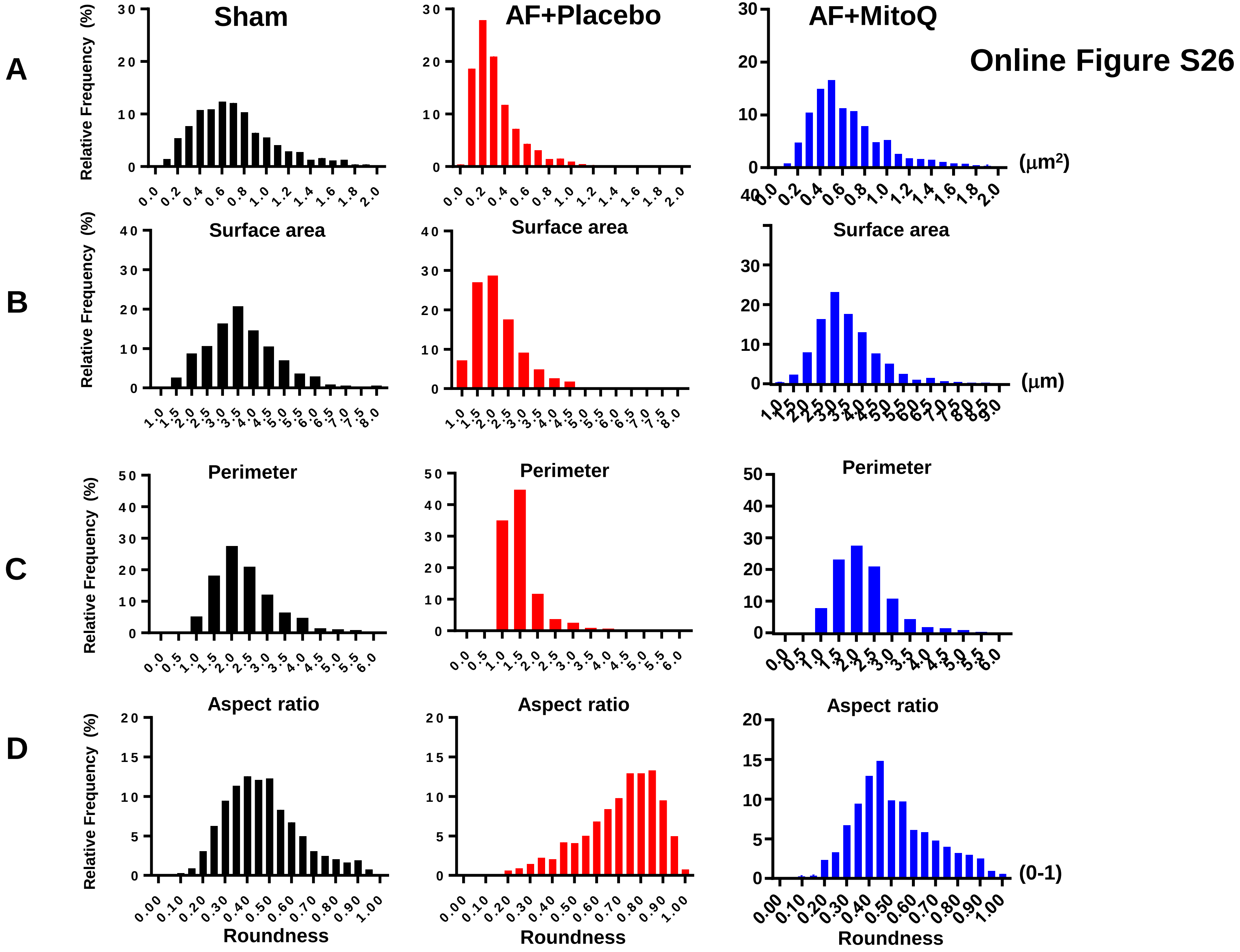


**Figure S26.** Histograms showing distribution frequency (% total mitochondria) of mitochondrial surface area, perimeter, aspect ratio and roundness from Sham, AF+Palcebo and AF+MitoQ left atria.

### Online Figure S27

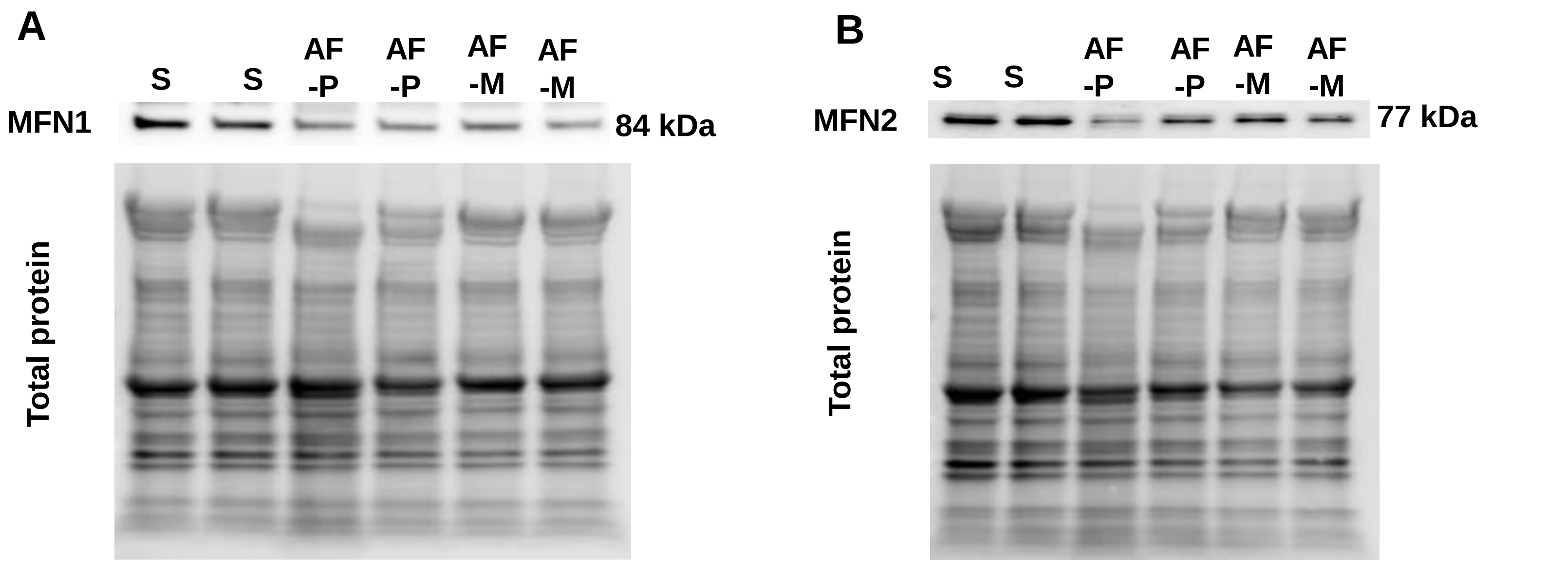


**Figure S27.** Western blot images of MFN1 (**A)**, MFN2 (**B)** and corresponding total protein from Sham, AF+Placebo and AF+MitoQ atrial CMs.

### Online Figure S28

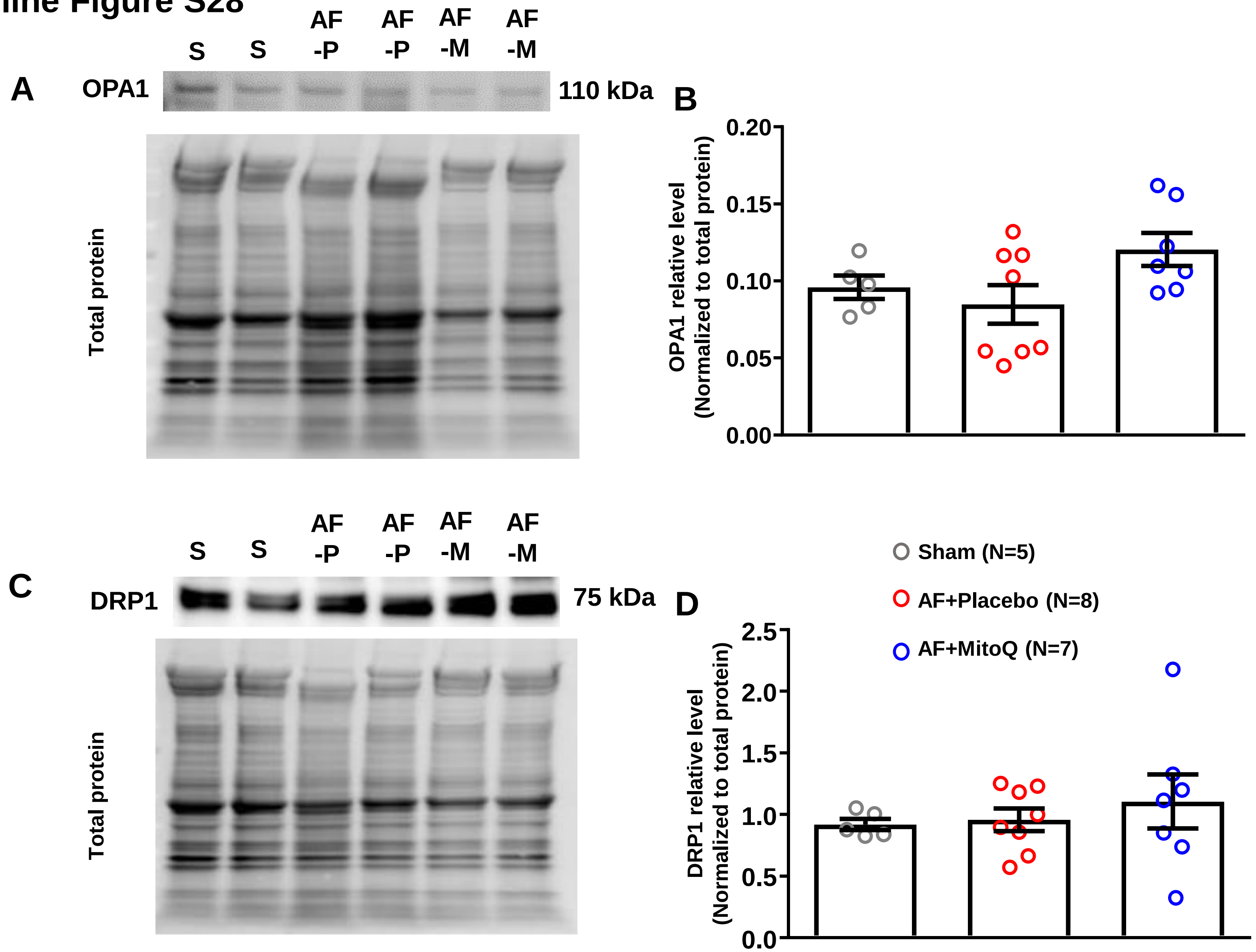


**Figure S28. A, C.** Western blot images of OPA1.DRP1 and total protein from Sham, AF+Placebo and AF+MitoQ atrial CMs; **B, D.** Individual-data point and mean±SEM OPA1 and DRP1 protein expression level.

#### Online Figure S29

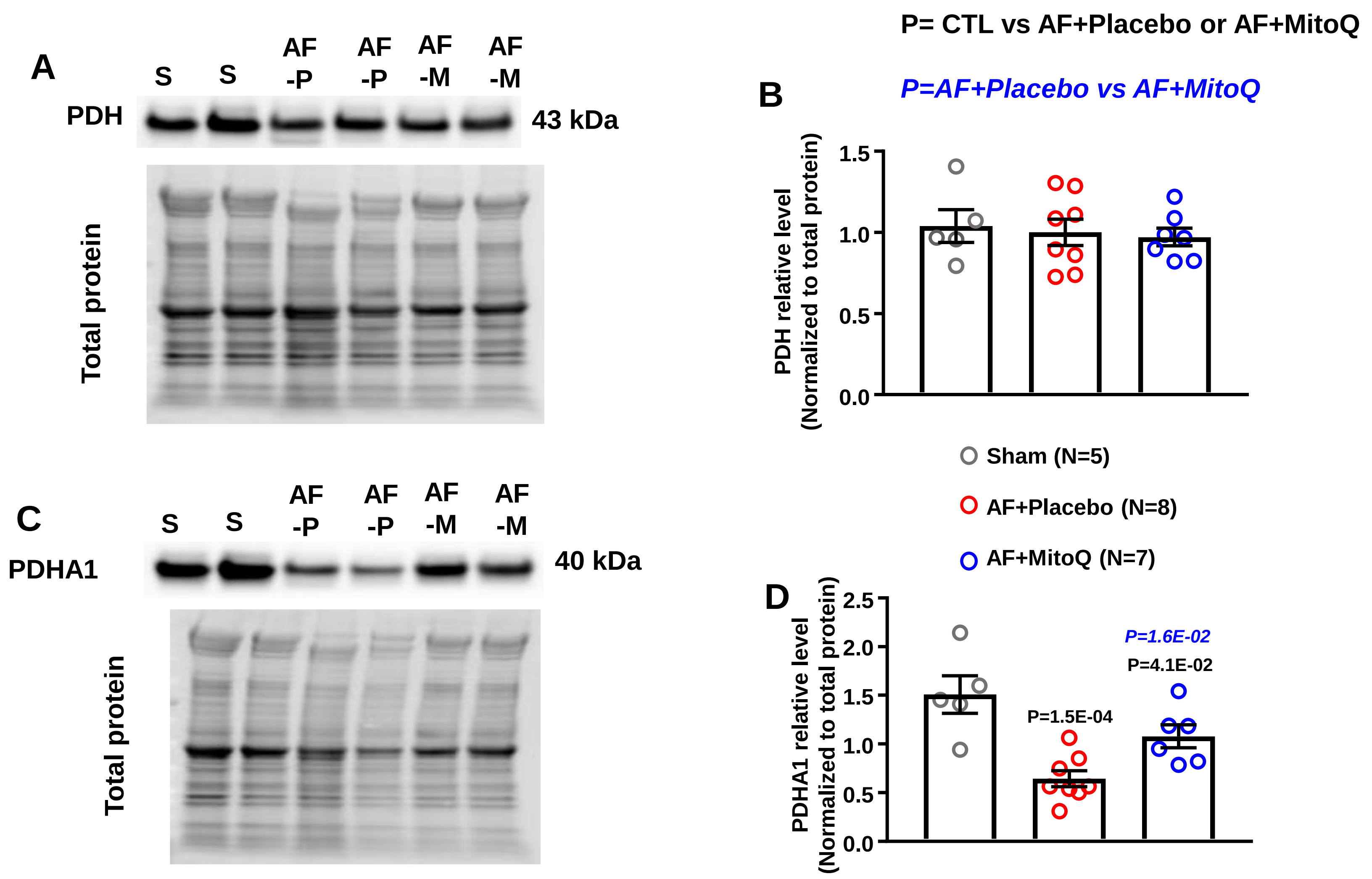


**Figure S29. A.** Western blot images of PDH and total protein from Sham, AF+Palcebo and AF+MitoQ atrial CMs; **B.** Individual-data point and mean±SEM PDH protein expression level; **C.** Western blot images of PDHA1 and total protein from Sham, AF+Palcebo and AF+MitoQ atrial CMs; **D.** Individual-data point and mean±SEM PDHA1 protein expression level.

### Online Figure S30

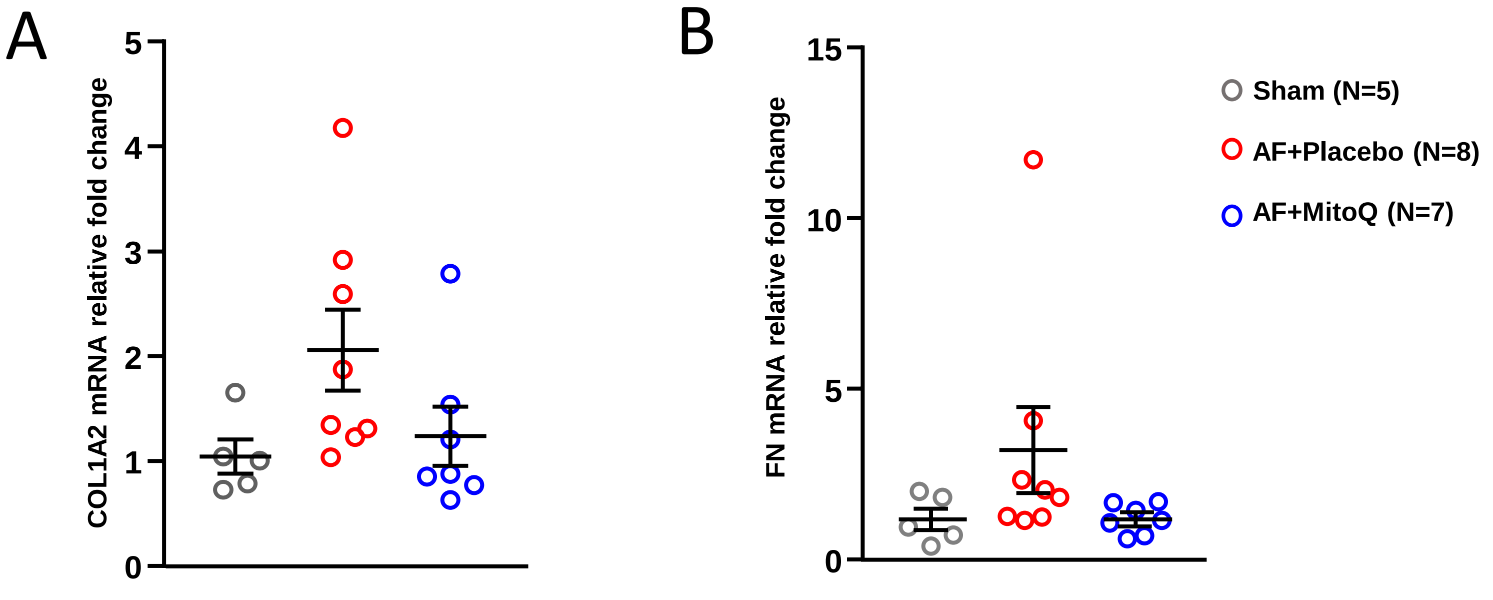


**Figure S30. A.** Fold-change (vs Sham mean) in mRNA expression of COL1A2; **B.** Fold-change in mRNA expression of FN; in Sham, AF+Placebo and AF+MitoQ atrial CMs. The differences among groups were not statistically significant.
