## Supplementary material for "Time-dependent Mitochondrial Remodeling in Experimental Atrial Fibrillation and Potential Therapeutic Relevance": Online method

### ONLINE METHODS SUPPLEMENT

#### Animal Model and In Vivo Study

Animal-care procedures were approved by the Animal Research Ethics Committee of the Montreal Heart Institute (protocol: 2020-47-07) and followed Canadian Council on Animal Care Guidelines. For the initial characterization study, a total of 58 adult Foxhound dogs were studied, divided into control (n=15, 23.4±3.4 kg, 2.0±0.3 years-old, Female/Male , F/M:7/8); 24-hour (n=11, 21.8±4.1 kg, 2.5±0.5 years, F/M:5/6 ), 3-day (n=10, 22.1±3.3 kg, 2.4±0.5 years, F/M:6/4), 1-week (n=11, 22.4±2.3 kg, 2.1±0.3 years, F/M:5/6) and 3-week AF (n=11, 21.8±2.7 kg, 2.4±0.5 years, F/M:5/6). No significant sex-based differences were noted in results, so results were combined for analysis. Dogs were anesthetized with ketamine (5.3 mg/kg i.v.)/diazepam (0.25 mg/kg, i.v.) and isoflurane (1.5%), intubated and ventilated. A unipolar pacing lead was inserted into the right-atrial (RA) appendage under fluoroscopic guidance and connected to a pacemaker in the neck. The pacemaker was programmed to maintain AF by pacing the RA at 600 bpm for 24 hours, 3 days, 1 week and 3 weeks. By maintaining AF electrically, this model mimics the atrial remodeling associated with spontaneous AF.[1] Dogs were anesthetized with morphine (2 mg/kg, s.c.) and α-chloralose (120 mg/kg, i.v., followed by 29.25 mg/kg/h) on day 1, 3, 7 and 21, and ventilated mechanically. Effective refractory periods (ERPs) were measured at basic cycle lengths of 150, 200, 250, 300, and 350 ms in the left-atrial (LA)-appendage, with 10 basic stimuli (S1) followed by a premature extrastimulus (S2) with 5-ms decrements. AF was induced by atrial burst pacing at 50 Hz and 10-V 2-ms square-wave output. Mean AF-duration was based on 10 AF-inductions in each dog. If the mean duration of the first 5 episodes of AF was >2 min , AF was induced only 5 times. When AF lasted more than 10 minutes it was considered sustained, and no further AF-inductions were performed. Hemodynamic data were obtained with fluid-filled catheters and transducers.[2,3]

For the in vivo mitochondrial-targeted treatment study, 20 additional Foxhound dogs (21.4±1.7 kg, 2.1±0.3 years, F/M:12/8) were studied in the following 3 groups ⑴ Sham group, instrumented but without atrial tachypacing (N=5); ⑵ AF+Placebo group, maintained in AF by atrial tachypacing and receiving placebo (empty capsule) once a day for 3 weeks (N=8); ⑶ AF+MitoQ group, maintained in AF with atrial tachypacing receiving MitoQ (5 $mg.kg^{-1}$ orally, once a day, N=7) for 3 weeks. The experimenter was blinded to dog therapy-assignment until the experiments were completed and data analyzed. MitoQ treatment was initiated 3 days before tachypacing onset. Mitoquinone (MitoQ) was kindly provided by MitoQ Ltd (New Zealand).

#### Canine ACM Isolation and Culture

ACMs were isolated with previously described methods.[4,5] Dogs were anesthetized with morphine (2 mg/kg s.c.) and α-chloralose (120 mg/kg i.v.) and mechanically ventilated. The heart was removed after intra-atrial injection of heparin (10,000 U), immersed in 1.8 mM $Ca^{2+}$-containing Tyrode's solution, the left coronary artery was cannulated, and left-atrial tissue perfused with 1.8 mM $Ca^{2+}$-containing Tyrode's solution (37°C, 100% $O_2$), then with $Ca^{2+}$-free Tyrode's solution (~10 minutes), followed by ~60-min perfusion with the same solution containing collagenase (~0.45 mg/mL, CLSII, Worthington, Lakewood, NJ) and 0.1% bovine serum albumin (BSA, Sigma–Aldrich, Oakville, ON). Tissues were minced and ACMs harvested. Isolated cardiomyocytes were stored in 200 µmol/L $Ca^{2+}$-containing Tyrode's solution for confocal imaging, $Ca^{2+}$-transient and action potential recording, and in Kraftbruhe storage solution for L-type calcium current recording. For primary culture, ACMs were kept in PCell-

100X medium (WISENT INC., 001-045-CL), concentrated by centrifugation at 300 rpm (1 min), and cell-pellets removed. Cells were plated at low density onto laminin coated glass coverslips (25*25 mm). PCell-100x medium supplemented with 1% penicillin/streptomycin (Invitrogen) and 1% Insulin-Transferrin-Selenium for cell culture. After 2 hours, dead and unattached myocytes were removed and fresh medium was added. Pacing was accomplished with square-wave, 1-ms pulses (C-pace Cell-Culture Stimulator, IonOptix). Atrial cardiomyocytes were subjected to 24-hr in vitro pacing at 1 Hz (P1) or 3 Hz (P3). The rest of the cells were fast-frozen for subsequent biochemical studies. The right atrium was dissected free and the right coronary artery was cannulated and perfused with Krebs solution for optical mapping experiments.

#### Human Atrial Samples and Isolation of Human Atrial Cardiomyocytes

For experiments with human samples, tissue-specimens from right atrial appendages were obtained from patients (>18 years) undergoing elective open-heart surgery. Patients were categorized according to a pre-existing diagnosis of longstanding persistent (chronic) AF (cAF) or absence of such diagnosis (SR). Written informed consent was obtained from every patient prior to cardiac surgery. Experimental protocols were approved by the local ethical review board of the University Duisburg-Essen, Germany (#12–5268-BO).

Human ACMs were isolated from right atrial appendage tissue using enzymatic digestion as previously described.[6,7] After isolation, cells were suspended in EGTA-free storage solution containing (in mM): KCl 20, $KH_2PO_4$ 10, D-glucose 10, L-glutamic acid 70, beta-OH-butyrate 10, taurine 10, butanedione monoxime 10, supplemented with 1% (w/v) bovine serum albumin, pH 7.4.

#### Confocal Imaging

**Mitochondrial ROS (mtROS).**

##### 1) Canine Atrial Cardiomyocytes

For mitochondrial ROS detection, the laminin-coated coverslips with adherent myocytes were incubated with MitoSOX Red (2.5 μM, Thermofisher Scientific, M36008) for 70 min at 37$^{o}$ C in the dark[8] and then washed free of extraneous MitoSOX with 1.8 mM $Ca^{2+}$-containing Tyrode's solution. Fluorescence was monitored at excitation/emission wavelength ~510/580 nm. Figure S1A shows autofluorescence recordings of unlabeled atrial cardiomyocytes. Fluorescence intensity and images were acquired using a confocal microscope (Olympus Corp, Tokyo, Japan, with FluoView 1000), equipped with a ×40 and ×60 oil-immersion objective lens (N.A. 1.3) and an argon-ion laser. The scanning parameters were unchanged for all the scans. More than 100 cells/group were recorded for intracellular fluorescent intensity measurement. Images were analyzed offline using ImageJ software (Wayne Rasband, National Institutes of Health).[9]

##### 2) Human Atrial Cardiomyocytes

After stepwise reintroduction of extracellular $Ca^{2+}$ to a final concentration of 0.2 mM, freshly isolated human ACMs were incubated with 2.5 µM MitoSOX Red (Thermofisher Scientific, M36008) in 2.0 mM $Ca^{2+}$-containing Tyrode's solution for 60 min at ambient temperature and darkness. After 60 min, ACMs were pellet ed and resuspended in Tyrode's solution twice to wash off extracellular dye. Control cells were incubated for the same duration in Tyrode's solution without MitoSox Red. After incubation, cells were transferred to a chambered slide (Ibidi) for subsequent confocal imaging. Imaging was performed using a confocal laser scanning microscope (Leica TCS SP8) equipped with a x40 oil immersion objective (Leica, N.A. 0.7).

MitoSox Red was excited at a wavelength of 500 nm, and emission was recorded at 570-640 nm using a highly sensitive hybrid pixel detector (Leica). Laser intensity, pixel size, and pixel dwell times were constant for all imaging sessions. The confocal plane was chosen at the nucleus level in all cells. All ACMs were imaged on the same day as they were isolated. Cytosolic regions of rod-shaped human ACMs showing an intact sarcomeric structure in brightfield view were analyzed using standard ImageJ functionalities. As typical for human cardiomyocytes, some cells contained large quantities of lipofuscin granules and high autofluorescence levels (Figure S1B). These regions and the nucleus were excluded from fluorescence analysis.

**Analysis of Mitochondrial Membrane Potential ($\Delta\Psi_m$)**

Mitochondrial membrane potential was visualized in cardiomyocytes stained with tetraethyl benzimidazolyl carbocyanine iodide (JC-1, 5 μM, Thermofisher Scientific, T3168). JC-1 is a cationic carbocyanine dye that shows potential-dependent accumulation in mitochondria, indicated by a green fluorescence emission at (~529 nm) for the monomeric form (low concentrations) of the probe and yields green fluorescence, which shifts to red (~590 nm) with a concentration-dependent formation of red fluorescent J-aggregates (higher concentrations).[10] These characteristics make JC-1 a sensitive marker for mitochondrial membrane potential.[11,12] Atrial cardiomyocytes were loaded with JC-1 for 70 min at 37° C in the dark. After incubation, ACMs were washed free of extraneous JC-1 with 1.8 mM $Ca^{2+}$-containing Tyrode's solution. Red and green emission was recorded in ACMs with an Olympus confocal microscope. More than 100 cells/group were recorded for intracellular fluorescent intensities. Images were analyzed offline using ImageJ software.

**Mitochondrial Permeability Transition Pore (mPTP) Opening**

We used the cell-permeant fluorescent dye tetramethylrhodamine methyl ester (TMRM) to record rapid changes in. [13] ACMs were incubated with TMRM (200 nM, Thermofisher Scientific, I34361) for 20 min at room temperature in the dark. ACMs were loaded with TMRM, then washed with 1.8 mM $Ca^{2+}$-containing Tyrode's solution. TMRM was excited at 548 nm and emitted fluorescence acquired at >570 nm. TMRM-stained mitochondria appear as red rectangles and depolarization of the $\Delta\Psi_m$ due to mPTP-opening leads to TMRM release from mitochondria to cytosol, causing a decrease in mitochondrial fluorescence.[14] mPTP-opening was quantified in a stack of time-sequence images which were acquired every 30 s for 30 min via time-series image capture.

To enhance the signal-to-noise ratio, we employed a combination of techniques, including partial differential equation-based processing, anisotropic diffusion, and median filtering. Small-gradient noisy areas were smoothed using fourth-order diffusion, while areas with higher gradients remained unaltered. Median filtering was subsequently applied to remove spikes caused by noise. The regions corresponding to mitochondria were identified using image segmentation. This involved performing morphology dilation, followed by the identification of connected components, and finally computing the number of mPTP-openings. The accuracy of mPTP-opening quantification was confirmed and cross-checked manually after the initial calculation of mPTP-openings for each frame. To determine the frequency of mPTP-opening events, we recorded the total number of mPTP-openings observed within the field of view of the cell area during the recording period. When an mPTP event occurred in the region of a previously detected mPTP in the preceding frame, it was considered a continuation of the previous event. The duration of each mPTP event was calculated as the time difference between the first frame in which the mPTP was detected and the last frame of the event.

**Mitochondrial $Ca^{2+}$ Transient Measurement**

A cold-warm incubation protocol was used to load the mitochondria with Rhod 2-AM.[15-17] ACMs were loaded with Rhod 2-AM (7.5 μM) for 3 hours at 4$^{o}$C in PCell-100X medium. After cold loading, ACMs were incubated for 16 hours at 37°C in the same medium, followed by 10-min ACM perfusion with 1.8 mM $Ca^{2+}$-containing Tyrode's solution (35$^{o}$C) to selectively remove cytosolic Rhod 2-AM so that Rhod 2 remained only in the mitochondria. An Olympus confocal microscope (Olympus Corp, Tokyo, Japan, with FluoView 1000) equipped with a ×40 oil-immersion objective lens (N.A. 1.3) and an argon-ion laser was used for two-dimensional (2D) confocal mitochondrial $Ca^{2+}$-imaging. Excitation at 524 nm and emission at 589 nm were used for Rhod 2-AM fluorescence measurements. ACMs were field-stimulated with 2 platinum electrodes at 35°C.

**Cytosolic $Ca^{2+}$ transient measurement**

Canine atrial CMs were loaded with the $Ca^{2+}$ indicator Fluo-4 AM (10 μM, Thermo Fisher Scientific, Saint-Laurent, QC) for 25-30 minutes and then washed for 30 minutes to allow for deesterification of the dye. Two-dimensional (2D) confocal $Ca^{2+}$-imaging was performed. Cytosolic $Ca^{2+}$-transients were recorded in CMs loaded with Fluo-4 AM. An Olympus confocal microscope (Olympus Corp, Tokyo, Japan, with FluoView 1000) equipped with a ×40 oil-immersion objective lens (N.A. 1.3) and an argon-ion laser were used. For Fluo-4, excitation and emission wavelengths were 488 nm and >515 nm, respectively. Images were acquired in linescan mode (3 or 6 ms scan-1; pixel size 0.103μm/Pixel). CMs were field-stimulated with 2 platinum electrodes.

**$Ca^{2+}$ spark measurement**

The methodology for the detection and evaluation of spontaneous $Ca^{2+}$ sparks is as previously described.[18] In brief, atrial cardiomyocytes (ACMs) were incubated with 10 μM Fluo-4 AM (Invitrogen, Thermo Fisher Scientific) for 15 minutes. Subsequently, ACMs were placed on glass coverslips and superfused with Tyrode's solution containing 1.8 mM $Ca^{2+}$ for 10 minutes to facilitate intracellular de-esterification. Fluorescence images were acquired using a ZEISS LSM 5 LIVE confocal fluorescence microscope equipped with a 63×/1.4 Plan Apochromat oil objective. Fluo-4 AM was excited with a 488-nm argon laser, and emission signals were collected at wavelengths above 505 nm. Background-subtracted fluorescence emission signals (F) were normalized to the baseline fluorescence (F0) by averaging 20 images. Changes in intracellular calcium concentration ($[Ca^{2+}]_i$) are presented as ΔF/F0, where ΔF = F – F0 . The spontaneous $Ca^{2+}$ sparks frequency was determined using previously established methodologies. Briefly, a region mask was applied to define the cell outline and minimize interference from the cell exterior. A combination of a nonlinear partial differential equation-based approach, anisotropic diffusion, and median filtering was employed to enhance the signal-to-noise ratio. Fourth-order diffusion was applied to smooth areas with small gradient noise, while regions with larger gradients remained unaffected. Median filtering was subsequently employed to eliminate impulsive spikes generated by noise. The detection process utilized image segmentation to identify spark regions. Initially, morphology dilation was performed, followed by labeling of connected components and subsequent calculation of spark numbers and size. Upon the initial computation of sparks for each frame, the spark count was verified and recorded. In cases where spark centers coincided with the region of a prior spark, newly detected sparks were considered a continuation of the event in the previous frame and assigned the same identification (ID) number. Spark frequency was determined as the total count of detected $Ca^{2+}$ events over the recording time within the cell's field of view. Spark amplitude was quantified as the maximum F/F0 ratio during a spark event, representing the highest relative fluorescence ratio.

**Nicotinamide Adenine Dinucleotide (NADH) Autofluorescence**

NADH autofluorescence reflects the activity of the mitochondrial electron transport chain (ETC) as well as substrate supply.[19] NADH autofluorescence was measured in AACMs adherent to 25x 25 mm laminin-coated glass coverslips with an inverted microscope equipped with a ×40 objective. Excitation was provided by a mercury arc lamp at a wavelength of 350 nm, with exposure to UV light controlled by an electronic shutter (Optikon, model T132, Vincent Associates) anchored between the arc lamp and epifluorescence attachment of an inverted Olympus microscope (x40 objective). Emission was collected using a 400-500 nm band pass filter.[20] We measured the basal levels of NADH and calculated redox indexes by representing basal NADH levels as a percentage of the difference between the maximally oxidized and maximally reduced signals. ACMs were exposed to the uncoupling agent 1 μM carbonyl cyanide 4-(Trifluoromethoxy phenylhydrazone) (FCCP, Cayman, #15218) to stimulate maximal respiration and induce minimum NADH autofluorescence, respectively. After the fluorescent signal stabilized, the complex IV inhibitor 5 mM Sodium cyanide (NaCN, Millpore Sigma, # 380970-5), which fully inhibits respiration, was added to record the autofluorescence signal at its maximum (Figure S2A).[19,21,22]

**Flavin Adenine Dinucleotide (FAD) Autofluorescence**

The green autofluorescence emitted by FAD in its oxidized form was excited with an argon laser at 488 nm and emission was collected at >510 nm. The laser power was limited to prevent ACM damage. The mitochondrial uncoupler FCCP (1 μM) was applied to stimulate maximal respiration and oxidizes the mitochondrial $FADH_2$ pool in ACMs, resulting in increased fluorescence (maximal FAD signal). The subsequent application of the Complex IV inhibitor, 5 mM NaCN, suppresses respiration, preventing $FADH_2$ oxidation and decreasing the fluorescence signal (minimum FAD signal). The method for calculating the FAD redox index and estimating the FAD pool are illustrated in Figure S2B.[19]

#### Transmission Electron Microscopy (TEM)

TEM was used to evaluate mitochondrial structure. Briefly, LA tissue was sectioned into 1 $mm^3$ pieces, which were fixed with 2.5% glutaraldehyde in 0.1M sodium cacodylate buffer and 4% sucrose (pH=7.4, at 4$^o$C). The slices were then exposed to 1% aqueous osmium tetroxide and 1.5% aqueous potassium ferrocyanide for 2 hours at 4$^o$C and the sample washed by aspiration three times with $ddH_2O$. After dehydration, samples were embedded in Lx-112 (McGill University Facility for Electron Microscopy Research). Sections (75 nm) were cut (Leica Microsystems EM UC6 Ultramicrotome), stained with uranyl-acetate and lead-citrate, and visualized (FEI Tecnai G2 Spirit BioTwin 120kV Cryo-TEM, FEI Company, U.S.A). Samples were viewed with a Tecnai Spirit 120 kV electron microscope and images captured with a Gatan Ultrascan 4000 camera. For each animal at least thirty randomly selected sections were used for the analysis of mitochondrial morphology.

#### Patch- clamp recording

##### 1) Canine Atrial Cardiomyocytes

All in-vitro recordings were obtained at 35°C. The whole-cell perforated-patch technique was used to record action potentials (APs) in current-clamp mode and tight-seal patch-clamp to record currents in voltage-clamp mode. Borosilicate glass electrodes (Sutter Instruments) filled with pipette solution were connected to a patch-clamp amplifier (Axopatch 200B, Axon). Electrodes had tip resistances of 2-4 MΩ. For perforated-patch recording, nystatin-free intracellular solution was placed in the tip of the pipette by capillary action (~30 s), then pipettes were back-filled with nystatin-containing (600-μg/mL) pipette solution. Currents are expressed as densities (pA/pF). Junction potentials between bath and pipette solutions averaged 10.5 mV and were corrected for APs only.[5] Tyrode's solution contained (mM) NaCl 136, $CaCl_2$1.8, KCl 5.4, $MgCl_2$ 1, $NaH_2PO_4$ 0.33, dextrose 10, and HEPES 5, titrated to pH 7.3 with NaOH. The pipette solution for AP-recording contained (mM) GTP 0.1, potassium-aspartate 110, KCl 20, $MgCl_2$ 1, MgATP 5, HEPES 10, sodium-phosphocreatine 5, and EGTA 0.005 (pH 7.4, KOH). The extracellular solution for L-type $Ca^{2+}$-current ($I_{CaL}$) measurement contained (mM) tetraethylammonium-chloride 136, CsCl 5.4, $MgCl_2$ 1, $CaCl_2$ 2, $NaH_2PO_4$ 0.33, dextrose 10, and HEPES 5 (pH 7.4, CsOH). Niflumic acid (50-μM) was added to inhibit $Ca^{2+}$-dependent $Cl^-$-current, and 4-aminopyridine (2-mM) to suppress $I_{to}$. The pipette solution for $I_{CaL}$-recording contained (mM) CsCl 120, tetraethylammonium chloride 20, $MgCl_2$ 1, EGTA 10, MgATP 5, HEPES 10, and Li-GTP 0.1 (pH 7.4, CsOH).

##### 2) Human Atrial Cardiomyocytes

Freshly-isolated human ACMs were pelletted and resuspended with storage solution supplemented with 10mM EGTA. Cells were there then divided into two fractions and incubated overnight (15-18 hrs), either with or without the addition of 200nM MitoTempo at ambient temperature. On the next day, human ACMs were pelletted and resuspended in storage solution without EGTA. After stepwise reintroduction of extracellular $Ca^{2+}$ to a final concentration of 0.2 mM, measurements of $I_{CaL}$ were performed at ambient temperature using identical solutions and protocols as for dog atrial cardiomyocytes in whole cell ruptured-patch configuration.

#### Cell Contraction

ACM contractility was measured with the IonOptix Calcium and Contractility System (Ionoptix.com). Isolated atrial cells were transferred into an FHD recording chamber mounted on an Olympus X71 inverted microscope with an attached CCD camera and superfused with 1.8 mM $Ca^{2+}$ Tyrode solution at 37°C. ACMs were paced at 1Hz with an IonOptix MyoPacer field stimulator (pulse duration 0.5 ms; 20 volts). Raw data for sarcomere shortening and relaxation was collected and stored using IonWizard software.[23]

#### Optical Mapping

Hearts were excised and the RA was dissected free. The right coronary artery was cannulated and perfused with Krebs solution (mM: 120 NaCl, 4 KCl, 1.2 $MgSO_4$ 0.7, 1.2 $KH_2PO_4$, 25 $NaHCO_3$, 5.5 glucose, 1.25 $CaCl_2$, 95% $O_2$/5% $CO_2$) at a flow rate of 20 mL/min and a temperature of 37°C.  Silk ligatures were used to ligate atrial branches to stop leaks and maintain effective perfusion.[32] The preparation was loaded with di-4-ANEPPS following ten minutes of stabilization and electromechanical uncoupling with blebbistatin (15 μM, Biotium, CA). The tissue was then excited with a laser (Thorlabs, Newton, NJ) with a wavelength of 520±45 nm and emission recorded at 700±35 nm. Fluorescence images were recorded with a charge-coupled device camera (CardioCCD, Redshirt Imaging) focused on a 1.5×1.5-cm square. A pair of bipolar electrodes was placed on the superior right-atrial appendage to pace the tissue with 2-ms pulse-width, 1.5×threshold square-wave current pulses. Conduction velocity and AP-duration (APD) were recorded at basic cycle lengths (BCLs) of 500, 400, 300, and 200 ms. Time from the maximum upstroke velocity ($dF/dt_{max}$) until 80% repolarization was used to determine APD80. CV was calculated from the gradient of the scalar field of the isochronal activation maps. AF was induced by burst pacing at 20 Hz (2 ms, 4Xthreshold-current, 10 inductions), after which optical recording continued for 5 seconds. A spontaneously maintained fast (>600 bpm) irregular atrial rhythm lasting for 5 seconds was defined as inducible AF. When AF persisted for >20 minutes, it was stopped by pouring cold Kreb solution on the RAA to cease electrical activity, and a 15-minute rest time was allowed for recovery. Data were processed with a custom analysis-routine written in Matlab (The MathWorks).[24]

#### Quantitative Real-Time PCR

Total RNA was extracted from dog LA tissue with the miRNeasy Mini Kit (217004, Qiagen, Germany) according to the manufacturer's instructions. RNA concentration was quantified using Thermo Scientific NanoDrop 2000 spectrophotometer. The primary cDNA was synthesized with TaqMan Transcription Kit (4368813, Applied Biosystems, Lithuania) for mRNA. The following settings were used for reverse transcription: reverse transcription 25°C 10 min, 37°C 120 minutes, stop reaction 85°C 5 min, and hold 4°C. Quantitative PCR was performed with TaqMan probes (Table SII) and TaqMan Universal PCR Master Mix (4444557, Applied Biosystems, Lithuania) on an Applied Biosystems StepOnePlus Real-Time PCR System with the following cycling mode: enzyme activation at 50°C for 2 minutes and 95°C 2 minutes and 40 cycles of denaturation 95°C 1 s and annealing/extension 60°C 20 s. All TaqMan assays were designed on StepOne™ Software (Applied Biosystems), with threshold at 0.2. Gene expression was calculated with $2^{-\triangle Ct}$ method and normalized to β2-microglobulin.

#### Western Blot

LA cardiomyocytes were lysed with three times volume of lysis buffer (150mM NaCl, 50mM Tris-HCl pH7.5, 0.1% TritonX-100,10% glycerol, 0.1% SDS, 1 mM phenylmethane sulfonyl fluoride and 10ul Halt™ Protease Inhibitor Cocktail (100X) (Thermo Scientific, # 78429) in 1 ml buffer, 1% phosphatase inhibitor cocktail 3 (Thermo Scientific, # P0044)). Equivalent amounts of protein were separated by sodium dodecyl sulfate polyacrylamide gel electrophoresis (SDS-PAGE) and transferred to polyvinylidene difluoride (PVDF) membranes. Total protein on membrane was determined with No-Stain Protein Labeling Reagent (Invitrogen, #A4449) and detected with the Bio-Rad ChemiDoc Imaging System. Membranes were blocked for 1 hour at room temperature and probed with primary antibodies overnight at 4°C. After extensive washing, membranes were further incubated with secondary antibodies conjugated to horseradish peroxidase and washed extensively at room temperature. Membranes were exposed to the superSignal™ West pico PLUS chemiluminescent Substrate (Thermo Scientific, #34577) and chemiluminescent signal was detected with the Bio-Rad ChemiDoc Imaging System or X-ray film (Radiomat LS, # XC6A2). After stripping in ReBlot Plus Strong Antibody Stripping Solution, membranes were re-probed with appropriate primary and secondary antibodies. Band intensities were quantified with Bio-Rad Image Lab software. Protein expressions were normalized to the total protein on the membrane.[4]

Proteins were isolated from human RA tissue homogenates, and protein levels were determined with Western blot according to standard protocols. Antibodies and working concentration used are listed in the Major Resources Table. We used the appropriate near-infrared fluorophore dyes as secondary antibodies and imaged using an Odyssey Infrared Imaging System (LI-COR Biosciences).

#### Histology

Atrial tissues were fixed in 10% neutral-buffered formalin solution, embedded in paraffin blocks, transverse sections (6 µm) were cut at room temperature and stained with Masson's Trichrome. Stained images were digitized, and the fibrotic area was analyzed with Image Pro 9.3 (Media Cybernetics, Rockville, Maryland). LA fibrosis (expressed as percent cross-sectional area) was quantified by an observer blinded to experimental group.[25]

#### Data Analysis

#### Canine Studies

Custom-made software (available at https://github.com/FengXiongCA/Nucleus-Cytosolic-Ca) was used to analyze $Ca^{2+}$-transients. Image J was used to analyze confocal images. GraphPad Prism 8.0, Origin 5.0 and SAS release 9.4 (SAS Institute Inc., Cary, NC, USA) were used for data analysis. Wherever multiple ACMs were measured per dog, numbers are given as n/N, where n indicates number of ACMs studied, from N dogs.

Mixed effects model was used for all analyses. This analysis allows the use of heterogeneous variance-covariance matrices for fixed effect and this was done when appropriate. Some of the analyses were multilevel; the level used for the data were defined by (1) the individual dog and (2) the cell within dog and random effect within the model was the intercept to take into account correlation of multiple cells within dogs. Some of the analyses involved fixed effect as repeated factor(s) and the unstructured covariance structure was used to model the within-dog errors. For multilevel model involving repeated factor(s), the random effects within the model were the intercept and the repeated factor to take into account correlation within dogs and within repeated factor * dogs. The prevalence of sustained-AF induction was compared between groups with Fisher's exact test.

All variables with a non-normal distribution were transformed using a natural logarithm for analyses. $P$ values < 0.05 were considered to be statistically significant. Analyses were performed with SAS release 9.4 (SAS Institute Inc., Cary, NC, USA).

#### Human ACM Studies

For biochemical experiments and clinical parameters, for which each patient contributed a single data-point, unpaired two-tailed Student's t-tests were used when the data satisfied normal-distribution criteria. Non-normally distributed continuous data or data for which normality could not be assessed, were compared with Mann-Whitney tests. Categorical data were analyzed by Fisher's exact test.

For $I_{CaL}$ recordings, MitoTempo, and MitoSox experiments in which each patient may contribute multiple data points, data were defined (1) by the individual patient ID and (2) the ID of the cardiomyocyte within each patient. Sample sizes are given as n/N, where n=cardiomyocytes and N=patients. Multilevel mixed-effect models were employed to compare groups, as previously described.[26] The random effect within the model was the intercept to account for non-independent measurements in multiple cells from individual patients. Multilevel models were implemented in RStudio (Boston, MA) using lme4 with *P*-values derived using the Kenward-Roger approximation. Non-normally distributed data were log-transformed.
